# Discovery and characterization of cyclic peptides selective for the *C*-terminal bromodomains of BET family proteins

**DOI:** 10.1101/2022.12.25.521885

**Authors:** Charlotte Franck, Karishma Patel, Louise J Walport, Mary Christie, Alexander Norman, Toby Passioura, Hiroaki Suga, Richard J Payne, Joel P Mackay

## Abstract

**SUMMARY:** DNA encoded cyclic peptide libraries offer unique opportunities to discover high-potency, high-specificity ligands directed against a target protein. We set out to explore the potential for such libraries to provide ligands that can distinguish between bromodomains from the closely related paralogues of the Bromodomain and ExtraTerminal domain (BET) family of epigenetic regulators. Analysis of peptides isolated from a screen against the *C*-terminal bromodomain of family member BRD2, together with new peptides discovered in previous screens against the corresponding domain from BRD3 and BRD4, reveals peptides with nanomolar and subnanomolar affinities. X-ray crystal structures of several of these bromodomain-peptide complexes reveal diverse structures and binding modes, which nevertheless display several conserved binding features. A subset of the peptides demonstrates significant paralogue-level specificity, though structural analysis does not reveal clear physicochemical explanations for this specificity. Our data demonstrate the power of cyclic peptides to discriminate between highly similar proteins with high potency and hint that differences in conformational dynamics between BET-family bromodomains might modulate binding affinities amongst family members for particular ligands.

## INTRODUCTION

Cyclic peptides are a distinct and versatile group of bioactive lead molecules with potential for application as pharmaceuticals, agrichemicals, and biochemical tools (Banting et al., 1922; Montesinos and Bardají, 2008). Although a range of clinically valuable cyclic peptides, including cyclosporin and the vancomycin, actinomycin and polymyxin families of antibiotics, are derived directly from natural products, library screening strategies have the potential to uncover new classes of high affinity peptide ligands.

The <u>Ra</u>ndom nonstandard <u>P</u>eptide Integrated <u>D</u>iscovery (RaPID) platform utilizes mRNA display to express designer peptide libraries of 10^12^ or more members (**Supplementary Figure 1A**) (Passioura and Suga, 2017). By reassigning one or more codons to unnatural amino acids using a flexible aminoacylation ribozyme (Murakami et al., 2006; Murakami et al., 2003; Niwa et al., 2009; Saito and Suga, 2001; Saito et al., 2001), library diversity can be increased and altered in a targeted manner. Peptides are frequently produced as stable macrocycles *via* reprogramming of the initiating methionine with an *N*-terminal chloroacetylated amino acid to allow formation of a thioether with a cysteine residue that appears later in the sequence (often at the *C*-terminus). Libraries can thus comprise peptides with varying ring sizes and compositions, with cyclisation offering conformational stabilization and thus higher affinity and selectivity (Goto et al., 2009; Kawakami et al., 2009; Sako et al., 2008).

The ability of RaPID-derived cyclic peptides to provide such properties facilitates demanding applications such as distinguishing between closely related targets (Kawamura et al., 2017; Vinogradov et al., 2022). The gene regulatory proteins BRD2, BRD3, BRD4 and BRDT (the BET family) are paralogues that each contain two highly similar bromodomains (BDs); these domains feature a deep pocket that allows them to recognize *N*^*ε*^*-*acetyl-lysine (AcK) residues in post-translationally modified histones and other transcriptional regulators (**Supplementary Figure 2**) (Zaware and Zhou, 2019). BET proteins control the expression of distinct set of genes that are important for cell proliferation, cell cycle progression and apoptosis and have been identified as promising therapeutic targets for diseases that range from cancer to inflammatory disorders, heart disease and diabetes (Belkina and Denis, 2012; Filippakopoulos et al., 2010; Gilham et al., 2016; Miyoshi et al., 2008; Nicodeme et al., 2010; Padmanabhan et al., 2016).

Because BET proteins play roles in many cellular processes, specificity is likely to be important for realizing the therapeutic potential of BET inhibitors. Hundreds of small molecules that bind BET BDs have been described but only very rarely has selectivity for one paralogue been demonstrated over the others (Liu et al., 2020), a direct consequence of the high degree of sequence identity between the AcK binding sites in each family member. As a result, most small-molecule BET inhibitors that have entered clinical trials have been associated with adverse effects. Indeed, only recently have inhibitors emerged that display good selectivity between the first (BD1) and second (BD2) BDs of each BET protein, which have lower sequence similarity (Gilan et al., 2020; Yu et al., 2021). An inhibitor that can robustly distinguish between BD1s or between BD2s from different BET paralogues has not yet been discovered.

We have begun to explore the potential for RaPID to provide BET BD ligands with high affinity and specificity and recently reported the selection of macrocyclic peptide ligands for BRD3-BD2 and BRD4-BD2 (Patel et al., 2020). Here, we report a new RaPID screen directed against BRD2-BD2 and provide a deeper analysis and comparison of the peptide families that arose from this and the previous two screens. Although RaPID selection against BRD2-BD2 yields an enriched library of AcK-containing peptides, the cyclic peptides selected for characterisation do not display significant specificity for their cognate BD or for the BD2 domains over BD1s and, in certain cases, bind with stronger affinity to the BD1s. In contrast, expanding our analysis to explore peptide families arising from our previous BRD3-BD2 and BRD4-BD2 selections identified binders that demonstrate significant selectivity for BD2 domains over BD1 domains and even peptides that display a clear preference for the BD against which they were selected. Structural characterisation of these cyclic peptides in complex with BDs reveals a wide range of structural features and binding modes and helps identify conserved features and motifs when compared to previously characterised BET BD cyclic peptide ligands.

## RESULTS

### A RaPID screen against BRD2-BD2 enriches macrocyclic peptides with diverse sequences

Following our recent studies on the selection of cyclic peptide ligands for the BD2 domains of BRD3 and BRD4, we sought to identify cyclic peptide binders of BRD2-BD2. We performed an mRNA-based RaPID screen using a ∼10^12^-member peptide library in which genetic reprogramming permitted the incorporation of one or more AcK residues (**Supplementary Figure 1B**). One AcK was fixed in the centre of the peptide, given the known AcK-binding properties of BET BDs, to assist in targeting the AcK binding pocket of the BD. Initiation with a *N*-terminal *N*-chloroacetyl-L-tryptophan allowed spontaneous cyclisation with the thiol side chain of a *C*-terminal cysteine residue. Peptides in the library ranged from 10 to 17 amino acids with a randomized region of 7–14 residues split either side of the fixed AcK.

This library of peptides (ligated to the encoding RNA-DNA hybrid, **Supplementary Figure 1**), was screened against biotinylated BRD2-BD2 immobilised on streptavidin-magnetic beads. An enriched DNA library was recovered *via* reverse transcription and PCR for use as a new input library and in this manner a total of five rounds of selection was carried out. Next-generation DNA sequencing of the library after each round revealed a high degree of sequence enrichment, with the top sequence accounting for 2.3% of total sequencing reads by round 5 of the RaPID selection (**Supplementary Dataset 1**). All sequences contained one or more AcK residues, consistent with the library design.

We selected five peptides with diverse sequences for characterization, taken from the 15 most abundant peptides from the final enriched BRD2-BD2 library; these peptides are listed in **Figure 1A**. All peptides were synthesised by automated solid-phase peptide synthesis, cyclised using a basic solution comprising 5 vol% *i*Pr_2_NEt in dimethylformamide and purified using reverse-phase HPLC to afford the final peptides in isolated yields ranging from 1% to 16% based on the loading of the *C*-terminal amino acid to resin.

**Figure 1.**
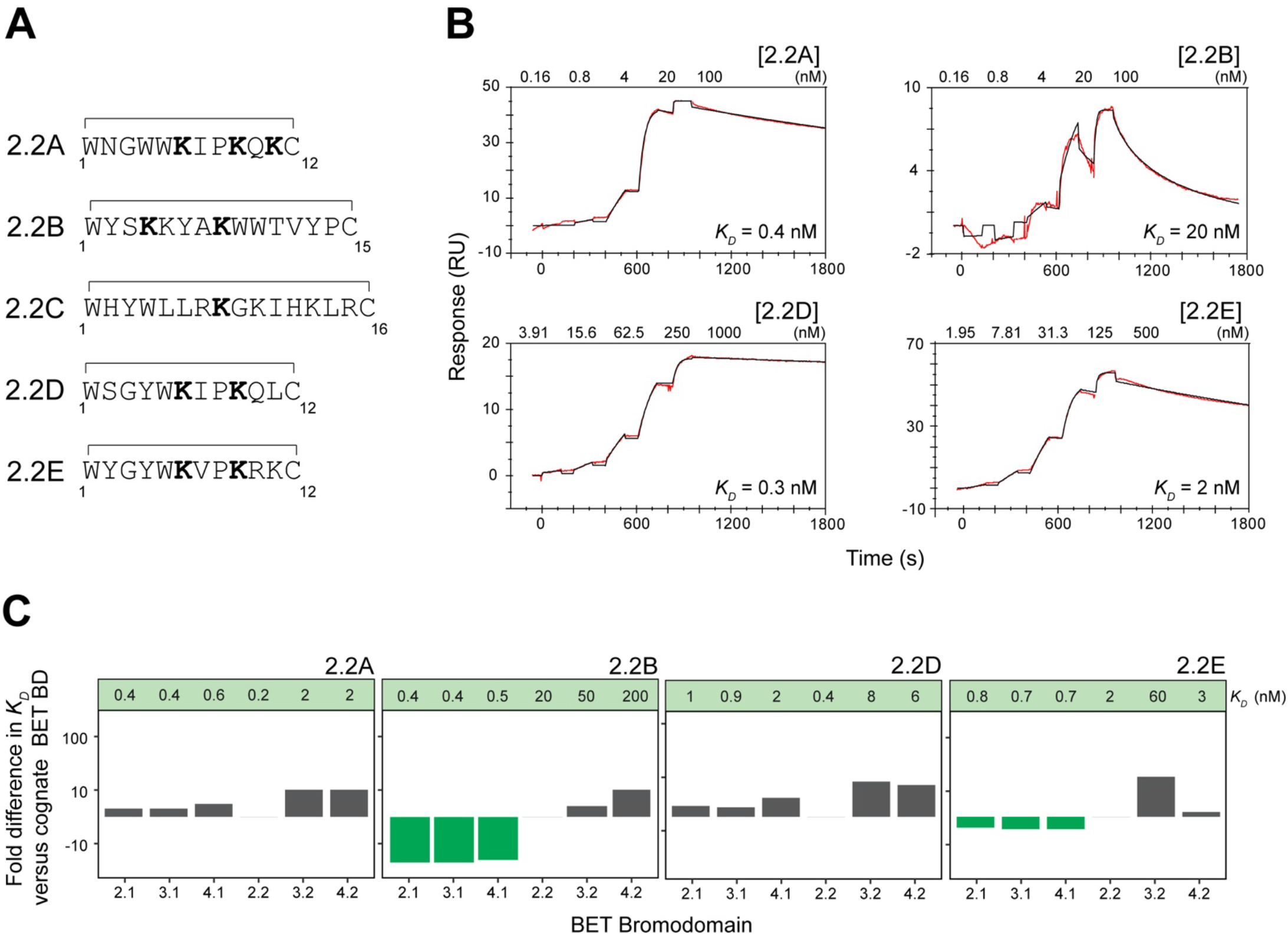
Cyclic peptides examined in this study from the RaPID selection against BRD2-BD2. **A**. Sequences of the peptides selected for study from the BRD2-BD2 RaPID screen. AcK residues are shown in bold and the residues that are cyclized are indicated. Each peptide name is prefixed with 2.2 to indicate which BD the peptide was selected against and suffixed with a unique letter to differentiate individual peptides. **B**. Representative SPR sensorgrams (red) for binding of the indicated peptides to BRD2-BD2. Fits to a 1:1 binding model (black) are shown for each sensorgram. The *K*_*D*_ values of the representative traces are provided. **C**. Fold change in *K*_*D*_ (measured by SPR) for each of the BRD2-BD2 selected peptides binding to all BDs from BRD2, BRD3, and BRD4. The *K*_*D*_ values for each interaction are given in a panel above the graph. The fold changes were calculated by taking the affinity of each peptide against BRD2-BD2, the BD against which they were selected, as equal to 1. Increases in affinity relative to BRD2-BD2 are shown in green and decreases in affinity are shown in grey. All *K*_*D*_ values are given as the geometric mean of a minimum of three independent measurements. Uncertainties are shown in **Supplementary Table 1**.

### RaPID peptides selected against BRD2-BD2 bind BET-family BD2 domains with nanomolar dissociation constants

We used surface plasmon resonance (SPR) to assess the affinity of each of the five chosen cyclic peptides for BRD2-BD2, BRD3-BD2 and BRD4-BD2, as well as for the three corresponding BD1 domains. Biotinylated BDs were immobilized on a streptavidin coated sensor chip and treated with each peptide in turn. With the exception of **2.2C**, which showed poor behaviour on the SPR chip, all peptides exhibited *K*_*D*_ values of between 0.2 nM and 20 nM for BRD2-BD2, the domain against which they were selected. Binding to the other BD2 domains was ∼2–30-fold weaker, though still in the nanomolar *K*_*D*_ range (**Figure 1B-C** and **Supplementary Table 1**). In addition, all four peptides also bound to the BD1 domains. Peptides **2.2A** and **2.2D** showed strong binding, comparable to the binding of the molecules to BRD2-BD2. Surprisingly, the remaining two peptides, **2.2B** and **2.2E**, bound even more tightly to the BD1 than BD2 domains by factors of ∼2–50-fold.

### 2.2E binds BRD2-BD2 in a compact but irregular conformation

We next used X-ray crystallography to delineate the structural mechanisms by which a subset of the BRD2-BD2 selected cyclic peptides recognized the BET BDs. First, the structure of peptide **2.2E** in complex with BRD2-BD2 was determined to a resolution of 1.6 Å (PDB ID: 8CV7; **Figure 2A, Supplementary Table 2**). Two essentially identical copies of the complex are observed in the asymmetric unit (RMSD of 0.06 Å^2^ over backbone atoms); we focus on one. In this complex, BRD2-BD2 adopts a conformation that is indistinguishable from previously reported structures of this domain [*e*.*g*., PBD ID 3ONI (Filippakopoulos et al., 2010)]. **2.2E** does not form an element of regular secondary structure, although it still forms a highly ordered and compact structure that features six internal backbone hydrogen bonds (including two that are bifurcated, **Figure 2B**) and an additional hydrogen bond between the sidechain amide of AcK6 and the carbonyl group of Cys12.

**Figure 2.**
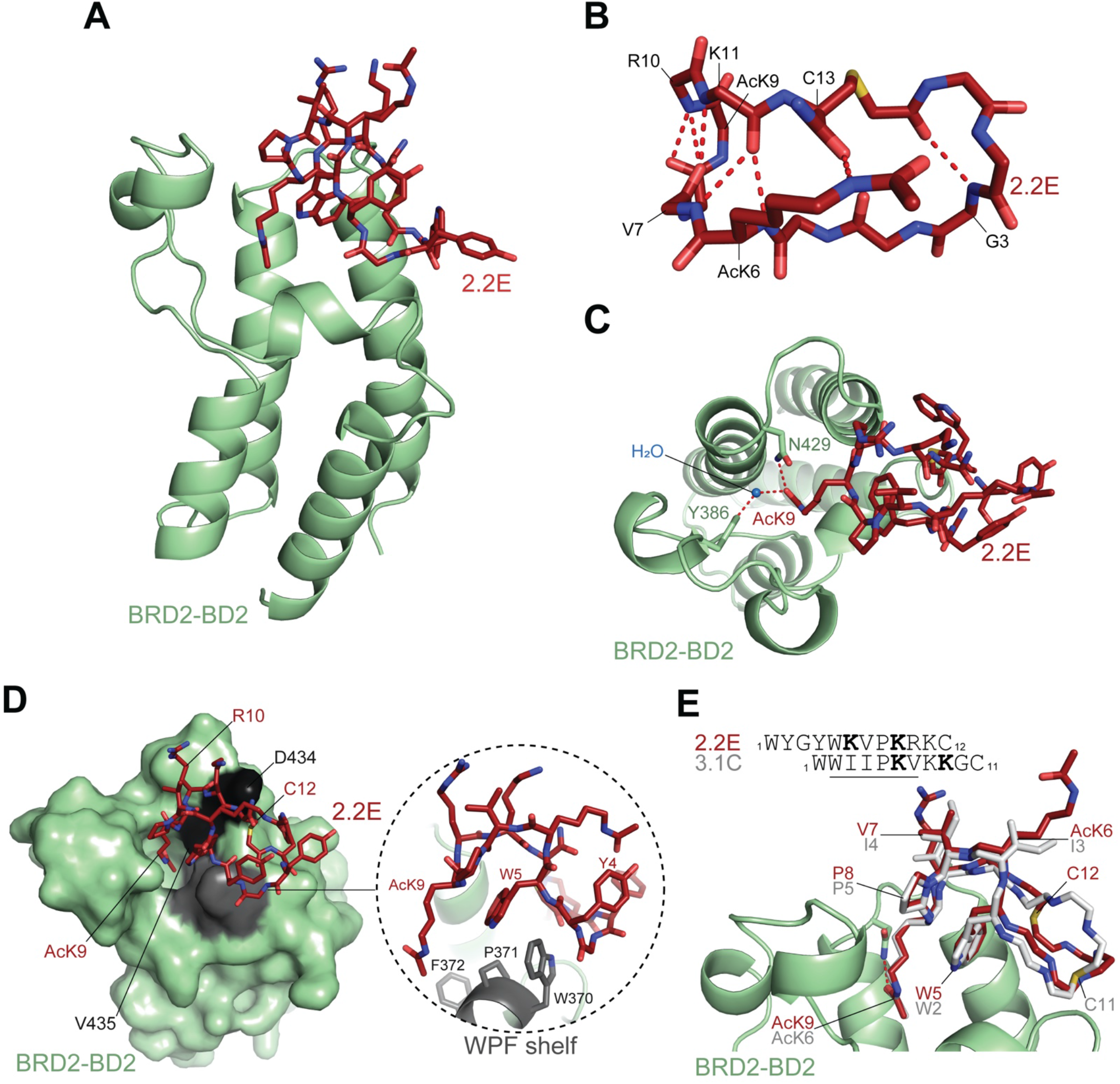
Structure of cyclic peptide 2.2E in complex with BRD2-BD2. The peptide in shown in red and BRD2-BD2 is shown in pale green. **A**. Ribbon/stick representation of the X-ray crystal structure of BRD2-BD2 in complex with **2.2E** (1.6 Å resolution; PDB ID: 8CV7). **B**. Backbone of **2.2E** in the structure of BRD2-BD2. The internal hydrogen bonds within the mainchain are indicated by red dashed lines. **C**. Hydrogen bonds made by AcK9 of **2.2E** in the binding pocket of BRD2-BD2 with conserved residues N429 and Y386 (mediated via a water molecule in the latter case). The hydrogen bonds are represented as red dashed lines. **D**. The key contact surface (shown in grey on BRD2-BD2; the WPF shelf is shown in light grey) made by **2.2E** in its interaction with BRD2-BD2. A close-up of the interaction between the WPF shelf and **2.2E** is shown in the right circled panel. **E**. Comparison of the BD binding mode conserved between **2.2E** and **3.1C** in complex with BRD2-BD2 (PDB ID: 6U71; only a single copy of BRD2-BD2 is shown for clarity). The structurally conserved motif is underlined in the sequence line-up at the top.

AcK9 contacts BRD2-BD2 in the AcK binding pocket (**Figure 2C**), entering at the canonical angle and forming the commonly observed hydrogen bonds with Asn429 (direct) and Tyr386 (water-mediated). Additional hydrogen bonds are also observed between the carbonyls of AcK9 and Arg10 of **2.2E** and the amide protons of Val435 and Asp434, respectively. Finally, the sidechain carboxyl of Asp434 interacts with the amide proton of Cys12 of **2.2E**. Extensive Van der Waals interactions are made between the peptide and the BRD2-BD2, in particular with the WPF shelf, a hydrophobic surface on the edge of the binding pocket (**Figure 2D**). This position is typically occupied by the sidechain of the more *N*-terminal AcK found in KXXK motifs of natural BET BD ligands (as illustrated by the interaction between BRD4-BD1 and diacetylated histone H4 peptide in **Supplementary Figure 2B**), but is here occupied by Trp5 as we have observed previously for other cyclic peptides (Patel et al., 2020).

The structure of this complex also revealed the existence of a BD-binding motif that is conserved between **2.2E** and the previously described peptide **3.1C** (Patel et al., 2020). Alignment of the sequences and structures of these peptides (**Figure 2E**) shows highly similar backbone and sidechain conformations for a six-residue stretch (Trp5–Arg10 in **2.2E**). Outside this motif, the structures diverge significantly, including the location of the cyclizing cysteine residue.

### 2.2B engages two BDs

An X-ray crystal structure was also obtained of peptide **2.2B** in complex with BRD2-BD1 (1.84 Å – PDB ID: 8DNQ; **Figure 3A, Supplementary Table 3**). Strikingly, this peptide was highly enriched in the RaPID selection for BRD2-BD2 but showed a ∼40–50-fold higher affinity for BD1 domains over BRD2-BD2 and ∼2.5–10-fold higher affinity over the remaining BD2s by SPR (**Figure 1C**). **2.2B** engages simultaneously with two BDs in the structure. Each BD adopts its canonical conformation, and the peptide displays an irregular conformation featuring five internal hydrogen bonds - two of these *via* the hydroxyl sidechain of Ser3. The peptide conformation (**Figure 3B**) is unusually ‘open’ among the seven cyclic peptides for which we have previously determined BD-bound structures (Patel et al., 2020).

**Figure 3.**
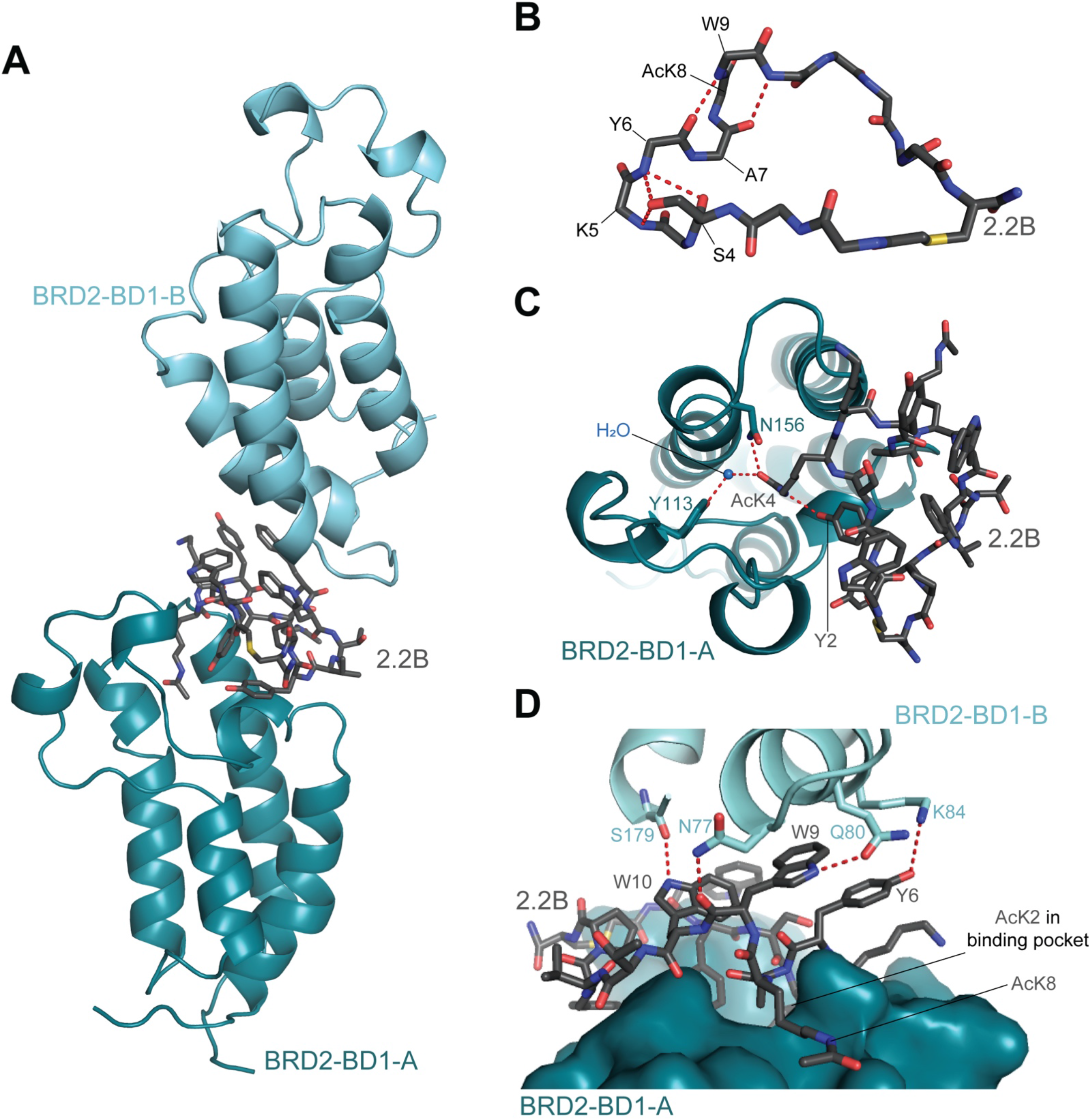
Structure of cyclic peptide 2.2B in complex with BRD2-BD1. The peptide is shown in dark grey and BRD2-BD1 is shown in blue. **A**. Ribbon/stick representation of BRD2-BD1 in complex with **2.2B** (1.84 Å resolution; PDB ID: 8DNQ). The peptide contacts two copies of the BD (BRD2-BD1-A and BRD2-BD1-B). **B**. Backbone of **2.2B** in the structure of BRD2-BD1. Internal hydrogen bonds within the mainchain and between the sidechain hydroxyl group of S4 and the mainchain of the peptide are represented as red dashed lines. **C**. Key interactions between **2.2B** and the BRD2-BD1-A (the BD that engages **2.2B** via the AcK-binding pocket). The hydrogen bonds formed between AcK4 of **2.2B** with N156 and Y113 (mediated via a water molecule in the latter case) in the binding pocket of BRD2-BD1 are indicated by red dashed lines. The hydrogen bond formed between the sidechains of AcK4 and Y2 of **2.2B** is also displayed. **D**. Key interactions between **2.2B** and BRD2-BD1-B. The hydrogen bonds formed between residues of **2.2B** and BRD2-BD2-B are indicated by red dashed lines.

The interaction with one BD occurs *via* the binding of AcK4 (entering the pocked vertically) and the formation of the typical hydrogen bonds between AcK4 and the Asn156 and Tyr113 sidechains on the BD (**Figure 3C**). Tyr2 in the peptide also partially enters the pocket and is sandwiched between the WPF shelf and the aliphatic portion of the AcK4 sidechain, with which it forms a hydrogen bond (**Figure 3C**). Despite **2.2B** having a second AcK residue (AcK8), the peptide does not engage the second BD *via* the binding pocket but rather with a surface on the opposite end of the BD. The interaction is mediated by a cluster of one Tyr (Tyr6) and two Trp residues (Trp9 and Trp10) in **2.2B** and Asp77, Gln80, Lys84, and S129 in the BRD (**Figure 3D**).

### Examination of additional hits from all RaPID screens against the BD2s of the BET family

Because significant selectivity for BRD2-BD2 was not observed among the peptides selected from the BRD2-BD2 RaPID screen, we expanded our previous analysis of BRD3-BD2 and BRD4-BD2 selections. In the original study (Patel et al., 2020), we examined only three of the several thousand peptides that were enriched. We therefore considered together the 500 most enriched peptides from each of the BD2 screens in order to further assess paralogue specificity. Alignment of these 1500 sequences (**Supplementary Table 4**) revealed multiple peptide families with high sequence similarity, many of which were common to all three selections and some that were unique to a single screen. We focused on peptides that were (a) highly enriched in the RaPID selections, (b) belonged to unique families of sequences compared to previously analysed sequences, and (c) possessed a short or mainly hydrophobic sequence to ultimately favour cell permeability. Four further peptides from the BRD3-BD2 selection and BRD4-BD2 screen were chosen for synthesis and characterisation based on these criteria (listed in **Figure 4A, Figure 5A**, and **Supplementary Table 1**).

**Figure 4.**
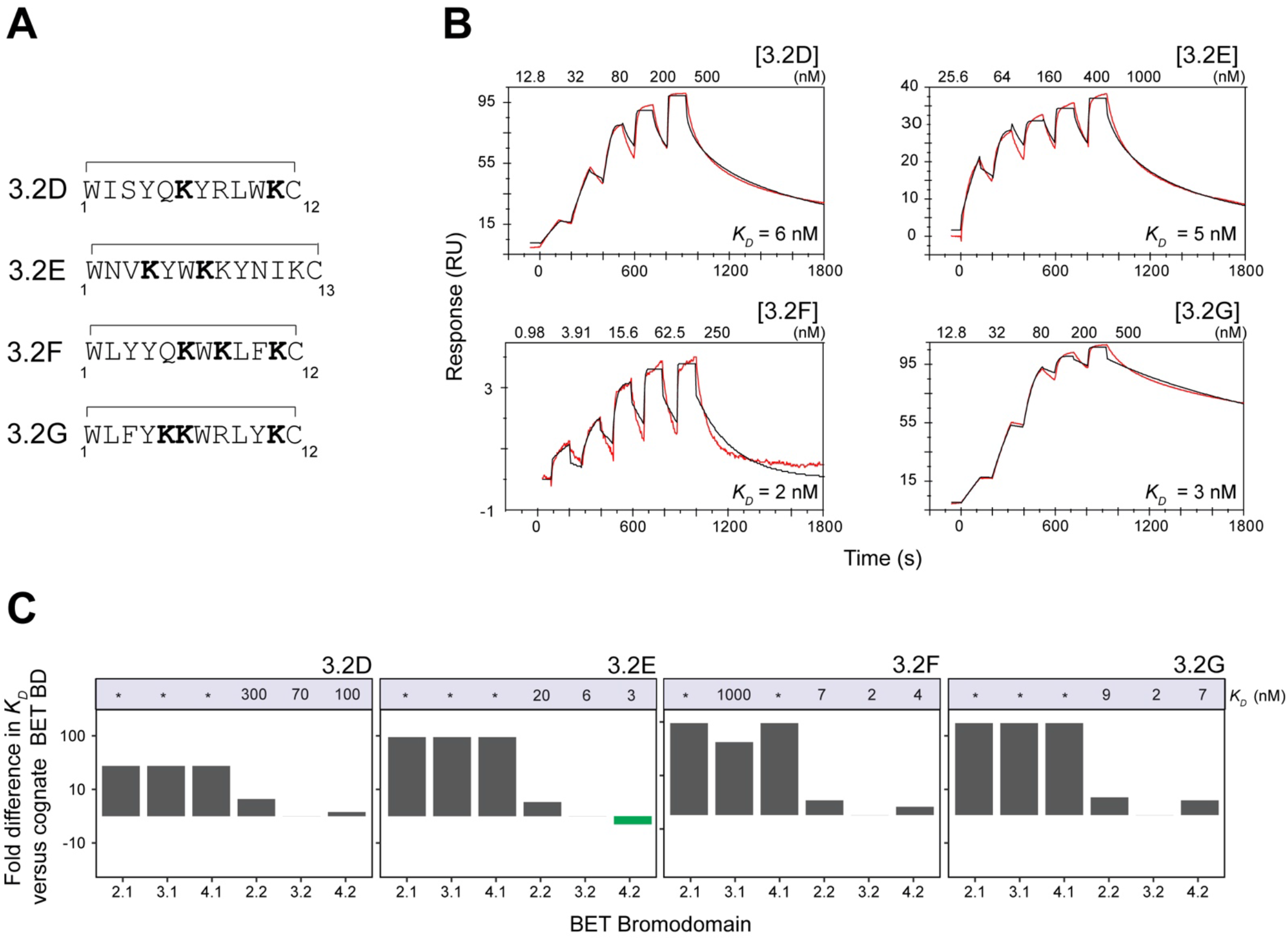
Cyclic peptides examined in this study from the RaPID selection against BRD3-BD2. **A**. Sequences of the peptides selected for study from the published BRD3-BD2 RaPID screen (Patel et al., 2020). AcK residues are shown in bold and the residues that are cyclized are indicated. As with the BRD2-BD2 screen peptides, the new peptides were prefixed to indicate the screen from which they were derived (*e*.*g*., 3.2 and 4.2) and suffixed with a unique letter identifier continuing from the naming system of our original report of the BRD3-BD2 and BRD4-BD2 screens where we looked at three peptides (*e*.*g*., A, B and C) from those selections. **B**. Representative SPR sensorgrams (red) for binding of the BRD3-BD2 RaPID peptides against BRD3-BD2, the bromodomain against which they were selected. Fits to a 1:1: binding model (black) are shown for each sensorgram. All *K*_*D*_ values are given as the geometric mean of a minimum of three independent measurements. The *K*_*D*_ values of the representative traces are provided **C**. Fold change in *K*_*D*_ (measured by SPR) for each of the BRD3-BD2 selected peptides binding to all BDs from BRD2, BRD3, and BRD4. The *K*_*D*_ values for each interaction are given in a panel above the graph. The fold changes were calculated taking the affinity of each peptide against BRD3-BD2, the BD against which the peptides were selected, as equal to 1. Increases in affinity relative to BRD3-BD2 are shown in green and decreases in affinity are shown in grey. All *K*_*D*_ values are given as the geometric mean of a minimum of three independent measurements. Asterisks (*) indicate no observed binding *via* SPR. Uncertainties are shown in **Supplementary Table 1**.

**Figure 5.**
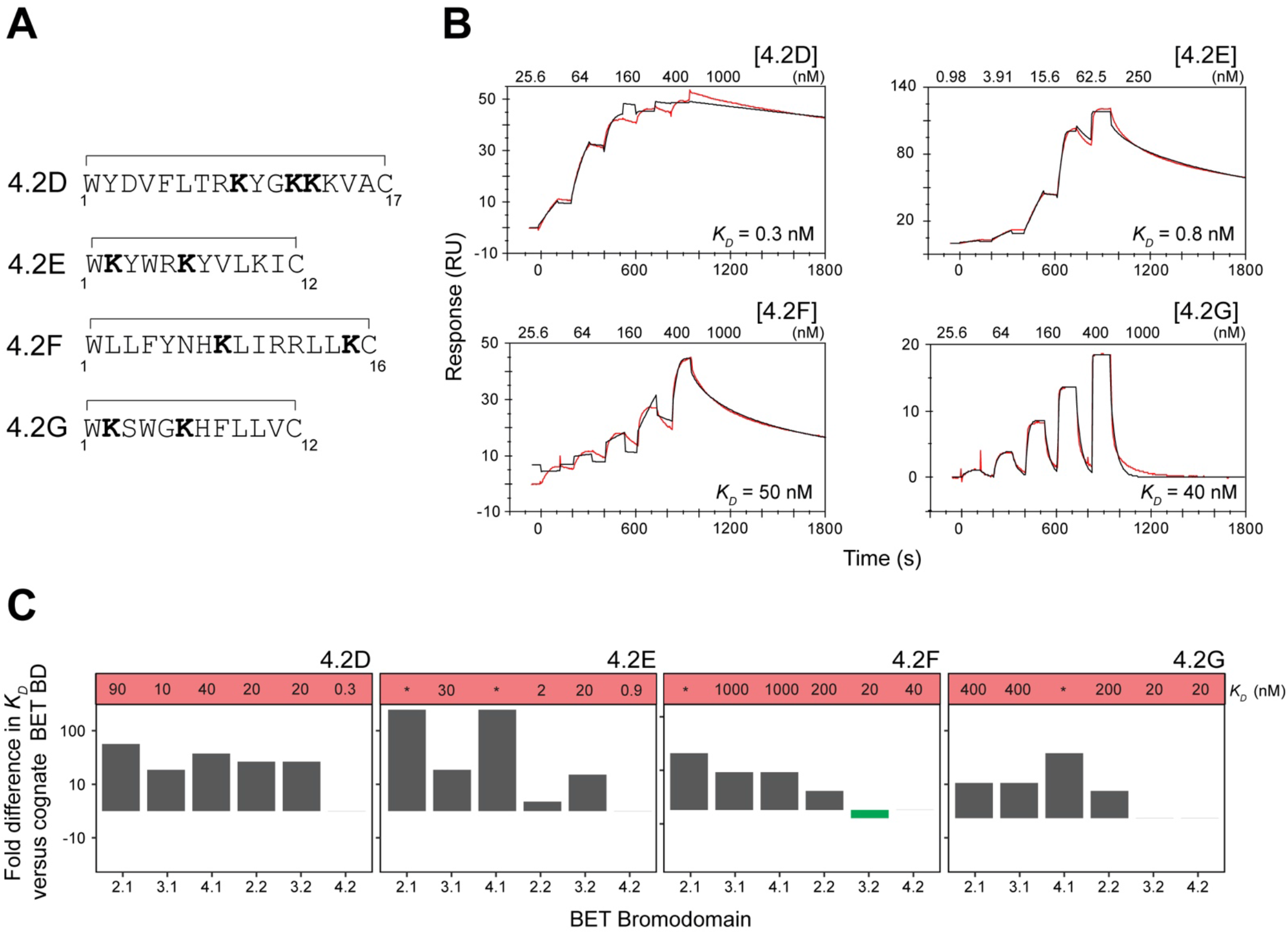
Cyclic peptides examined in this study from the RaPID selection against BRD4-BD2. **A**. Sequences of the peptides selected for study from the BRD4-BD2 RaPID screen (Patel et al., 2020). AcK residues are shown in bold and the residues that are cyclized are indicated. Each peptide name is prefixed with 4.2 to indicate which BD the peptide was selected against and suffixed with a unique letter to differentiate individual peptides. **B**. Representative SPR sensorgrams (red) for binding of the BRD4-BD2 RaPID peptides against BRD4-BD2, the bromodomain against which they were selected. Fits to a 1:1: binding model (black) are shown for each sensorgram. All *K*_*D*_ values are given as the geometric mean of a minimum of three independent measurements. The *K*_*D*_ values of the representative traces are provided. **C**. Fold change in *K*_*D*_ (measured by SPR) for each of the BRD4-BD2 selected peptides binding to all BDs from BRD2, BRD3, and BRD4. The *K*_*D*_ values for each interaction are given in a panel above the graph. The fold changes were calculated taking the affinity of each peptide against BRD4-BD2, the BD against which the peptides were selected, as equal to 1. Increases in affinity relative to BRD3-BD2 are shown in green and decreases in affinity are shown in grey. All *K*_*D*_ values are given as the geometric mean of a minimum of three independent measurements. Asterisks (*) indicate no observed binding *via* SPR. Uncertainties are shown in **Supplementary Table 1**.

The new set of peptides displayed significant selectivity for BD2 over BD1 domains in our SPR binding assay. This effect was particularly marked for the BRD3-BD2 selected peptides; essentially no binding was detected for the peptides to any of the BD1s, representing a selectivity of at least 250-fold or more for **3.2E, 3.2F**, and **3.2G**, and ∼15-fold for **3.2D** (**Figure 4B** and **Figure 4C**). **3.2D, 3.2F**, and **3.2G** further demonstrated at ∼2–5-fold stronger binding to the domain against which they were selected over the other BD2s. Peptides derived from the BRD4-BD2 selection, namely **4.2E, 4.2F**, and **4.2G** similarly bound the BD1 domains with between ∼2-3-fold weaker affinity than the BD2s (**Figure 5C** and **Figure 5B)**. Even more striking was the ∼60-fold selectivity of peptide **4.2D** for BRD4-BD2 over the paralogous BD2 domains from BRD2 and BRD3; this is the highest paralogue-level selectivity observed among the set of peptides examined here.

### 4.2E binds its target as a highly ordered β-hairpin with a ‘side-entry’ AcK

We used X-ray crystallography to delineate the molecular mechanisms by which a subset of the newly selected peptides recognized BDs. First, the structure of **4.2E** bound to BRD4-BD2, the domain against which it was selected, was determined to a resolution of 1.9 Å (PDB ID: 8CV4, **Figure 6A**; **Supplementary Table 5**). The asymmetric unit revealed two copies of the BD, each bound to a molecule of **4.2E** in an identical fashion (**Supplementary Figure 3A**). An additional molecule of **4.2E** was located adjacent to one of these BD-bound copies of **4.2E**, making three peptide-peptide hydrogen bonds but almost no interactions with the BD (**Supplementary Figure 3B**). We therefore focus our analysis on the 1:1 complex (the interaction of **4.2E** with BRD4-BD2-A shown in **Supplementary Figure 3A**). In this complex, BRD4-BD2 adopts a conformation that is essentially identical to previously reported structures of this domain (Shi et al., 2014; Zou et al., 2014). Peptide **4.2E** forms a regular and highly ordered β-hairpin that features six intramolecular hydrogen bonds (**Figure 6B**). One of the two AcK residues in **4.2E** (AcK2) engages the canonical AcK binding pocket of the BD, whereas the other (AcK6) is oriented away from the BD (**Figure 6C**). However, the angle at which AcK2 enters the pocket is very different to that observed for naturally occurring AcK-BD complexes; **Figure 6D** shows a comparison with the structure of BRD4-BD1 bound to a diacetylated peptide from histone H4 (PDB ID: 3UVW) (Shi et al., 2014).

**Figure 6.**
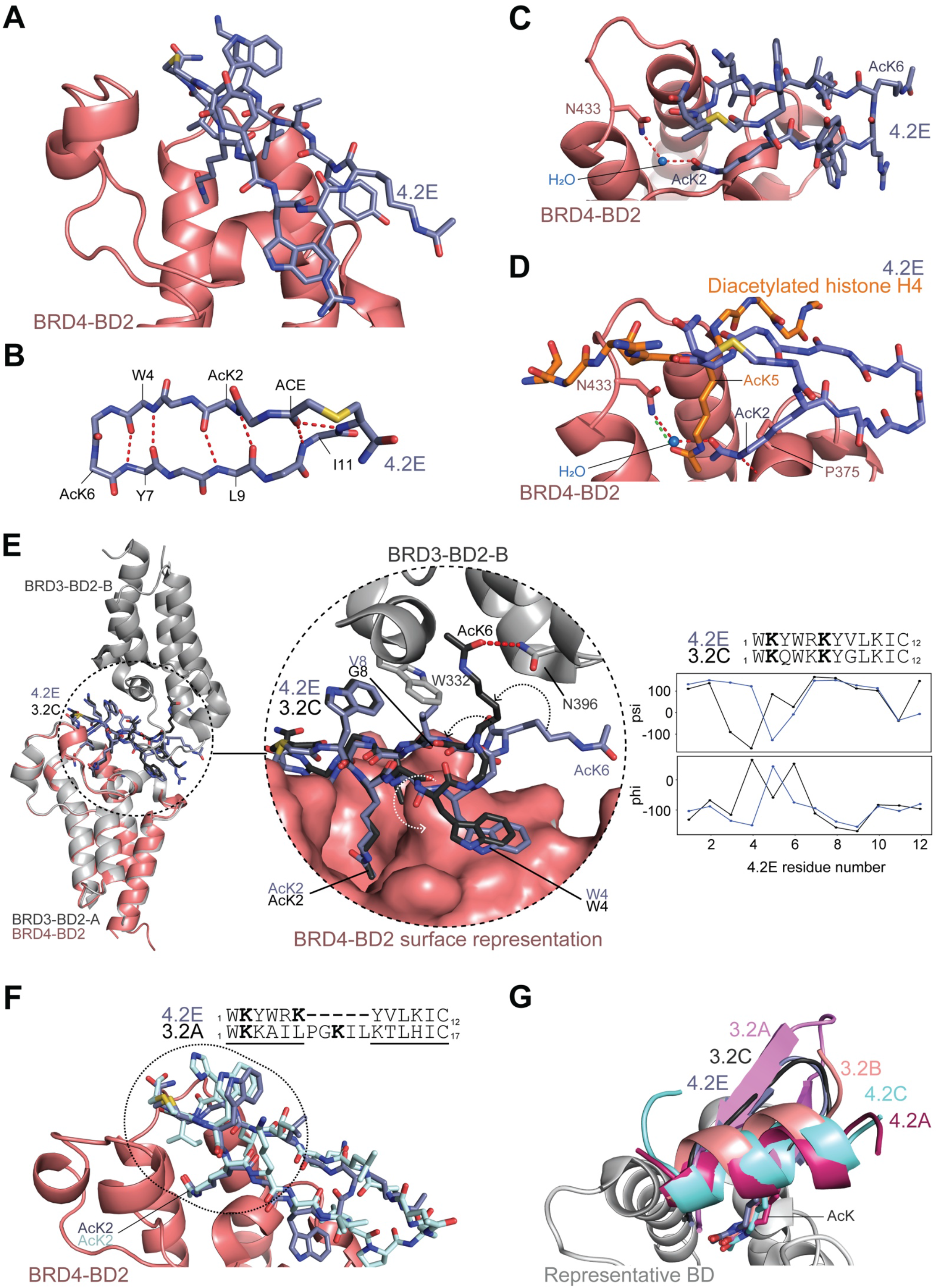
Structure of cyclic peptide 4.2E bound to the BRD4-BD2 bromodomain. The peptide is shown in mid-blue and BRD4-BD2 is shown in salmon. **A**. Ribbon/stick representation of the X-ray structure of BRD4-BD2 bound to 4.2E (1.9 Å resolution; PDB ID: 8CV4). **B**. Backbone of **4.2E** in the structure of BRD4-BD2. The internal hydrogen bonds formed within the mainchain are indicated by red dashed lines. **C**. The water mediated hydrogen bond formed between AcK2 of **4.2E** with Asn433 of BRD4-BD2. The hydrogen bonds are indicated by red dashed lines. **D**. Overlay of the BRD4-BD:**4.2E** structure with the structure of BRD4-BD1 in complex with a diacetylated histone H4 peptide (PDB ID: 3UVW). The diagonal entry of the AcK residue in **4.2E** into the binding pocket of the BD contrasts the vertical entry of the AcK residue (AcK5) observed for histone H4. **E**. *Left panel*: Overlay of the previously determined structure of BRD3-BD2 bound to cyclic peptide **3.2C** (Patel et al., 2020) onto the structure of the BRD4-BD2:**4.2E** complex. The **4.2E** and **3.2C** have closely related sequences and the two complex structures are highly similar. *Centre panel*: A close-up of the overlayed interactions between **4.2E** and **3.2C** in complex with their cognate BDs. Crankshaft motion of the backbone (indicated by dashed arrows) interconverts the conformations of **4.2E** and **3.2C**, involving a rotation of the AcK (AcK6) that allows **3.2C** to bind a second copy of BRD3-BD2. Residue V8 in **4.2E** that clashes with the Trp residue of the WPF shelf of BDs (W332 in BRD3-BD2) is shown. The presence of a Gly residue in this position in **3.2C** likely enables the engagement of a second BRD4-BD2 molecule by eliminating the clash. *Right panel*: Sequence alignment of **4.2E** and **3.2C** and the phi and psi angles of the two peptides are shown highlighting the differences in the backbone conformation. **F**. Overlay of **4.2E** and **3.2A**, highlighting the conformational similarity surrounding the cyclisation moiety. Similar regions in the sequence alignment of the two peptides are underlined and circled in the dashed ellipse. **G**. Comparison of the BD-binding pocket entry angle of the AcK residues in the β-hairpin structures of peptides **4.2E, 3.2A**, and **3.2C** with the entry angle of several α-helical peptides that we have previously described (Patel et al., 2020).

In line with this different entry angle, the highly conserved hydrogen bonding pattern that is observed in BET-ligand complexes is substantially disrupted. The water mediated hydrogen bond often observed with Tyr390 in other structures is absent in the **4.2E** complex, whereas the canonical hydrogen bond with Asn433, which is normally direct, is instead water mediated (**Figure 6C**). The ‘diagonal’ pocket entry angle does, however, result in the formation of a direct hydrogen bond between the Pro375 carbonyl and the AcK2 amide NH – an interaction that has not been observed in previous structures (**Figure 6D**).

We note that this ‘diagonal’ orientation of the pocket-binding AcK residue has been seen not only for AcKs embedded in β-hairpins but for a range of structurally diverse peptides. The helical **3.2B, 4.2A** and **4.2C** peptides from our recent study adopt this mode of binding (**Figure 6G**). It is notable that despite the lack of the direct and water-mediated hydrogen bonds with the conserved Asn and Tyr residues, respectively, our data show that the entry angle of the AcK residue into the pocket still allows a high-affinity interaction to take place.

Peptide **4.2E** also displays substantial selectivity for BRD4-BD2 over BD1 domains, with ∼30-fold selectivity over BRD3-BD1 and over a thousand-fold selectivity over the other BD1 domains. Further, **4.2E** displays a ∼20-fold selectivity over BRD3-BD2. The pattern is similar to that observed previously for the peptide **3.2C** (Patel et al., 2020), which was selected against BRD3-BD2 and has close sequence similarity to **4.2E** (**Figure 6E**; **Supplementary Table 1**). The BRD3-BD2:**3.2C** complex (PDB ID: 6ULP) has 2:1 stoichiometry (**Figure 4E**) and an overlay of the structures reveals that the backbone conformation of the two peptides is very similar for most residues (**Figure 6E**). However, a four-residue segment of **4.2E** (Gln3–AcK6) undergoes a crankshaft-style rotation that, remarkably, alters the position of only a single sidechain in the peptide: AcK6. This residue effectively rotates ∼120° in the **3.2C** complex to a position that allows it to enter the AcK binding site of a second copy of BRD3-BD2. Why does **4.2E** not take up this same conformation? The substitution of Gly8 in **3.2C** to valine in **4.2E** gives rise to a significant steric clash with Trp332 in the ZA loop of BRD3-BD2, blocking the binding of a second BD (**Figure 6E**) and demonstrating that stoichiometry can be controlled by subtle changes in the peptide sequence. Furthermore, **4.2E** binds BRD4-BD2 and BRD2-BD2 with 15 times and 45 times higher affinity, respectively, than does **3.2C**. The basis for this difference is, however, not clear. All residues in BRD4-BD2 that are within 5 Å of **4.2E** are conserved in both BRD2 and BRD3, suggesting the possibility that the differences in affinity might reflect differences in dynamics between the paralogues.

There is also significant conformational similarity between **4.2E** and another β-sheet peptide (**3.2A**) reported in our previous study (Patel et al., 2020), which was selected in the screen against BRD3-BD2. Like **3.2C**, the conformation of the **3.2A** peptide in the vicinity of the cyclization moiety closely resembles **4.2E**, as does the position and binding mode of the BD-binding AcK residue. **Figure 6F** demonstrates this structural similarity and highlights the residues for which this conformational similarity holds. A nine-residue sequence encompassing C- and N-terminal residues is the structurally conserved motif, and the structures show that this motif can be linked by sequences ranging from 2–7 residues in length.

### 4.2D has a short helical structure

Finally, we determined the X-ray crystal structure of **4.2D** bound to BRD4-BD2 (1.7 Å, PDB ID: 8CV6, *K*_*D*_ = 0.3 nM) in an effort to understand its observed preference for BRD4-BD2 over the other BDs (**Figure 7A**; **Supplementary Table 6**). **4.2D** features a short α-helix and displays a high degree of organisation, with 16 internal hydrogen bonds, including a bifurcated hydrogen bond between the carbonyl of Thr7 and the amide nitrogens of AcK12 and Gly11v(**Figure 7B**). Although **4.2D** bears three AcK residues, only AcK13 engages the BD target. This residue enters the binding pocket vertically, allowing for the canonical hydrogen bonding pattern with Asn433 and Tyr390 of the BD (**Figure 7C**). Additionally, Tyr2 in the peptide also takes up the same position that is observed for Tyr2 in the complex of **2.2B** with BRD2-BD2 described above – that is, lying between the pocket-binding AcK and the hydrophobic WPF shelf (**Figure 7D**). This similarity is striking given that the Tyr and AcK residues are embedded in completely different peptide backbone conformations in the two structures (**Figure 7D**). The Tyr and AcK are separated by a single residue in **2.2B**, but by 10 residues in **4.2D**. Several other intermolecular hydrogen bonds are formed, together with extensive van der Waals contacts.

**Figure 7.**
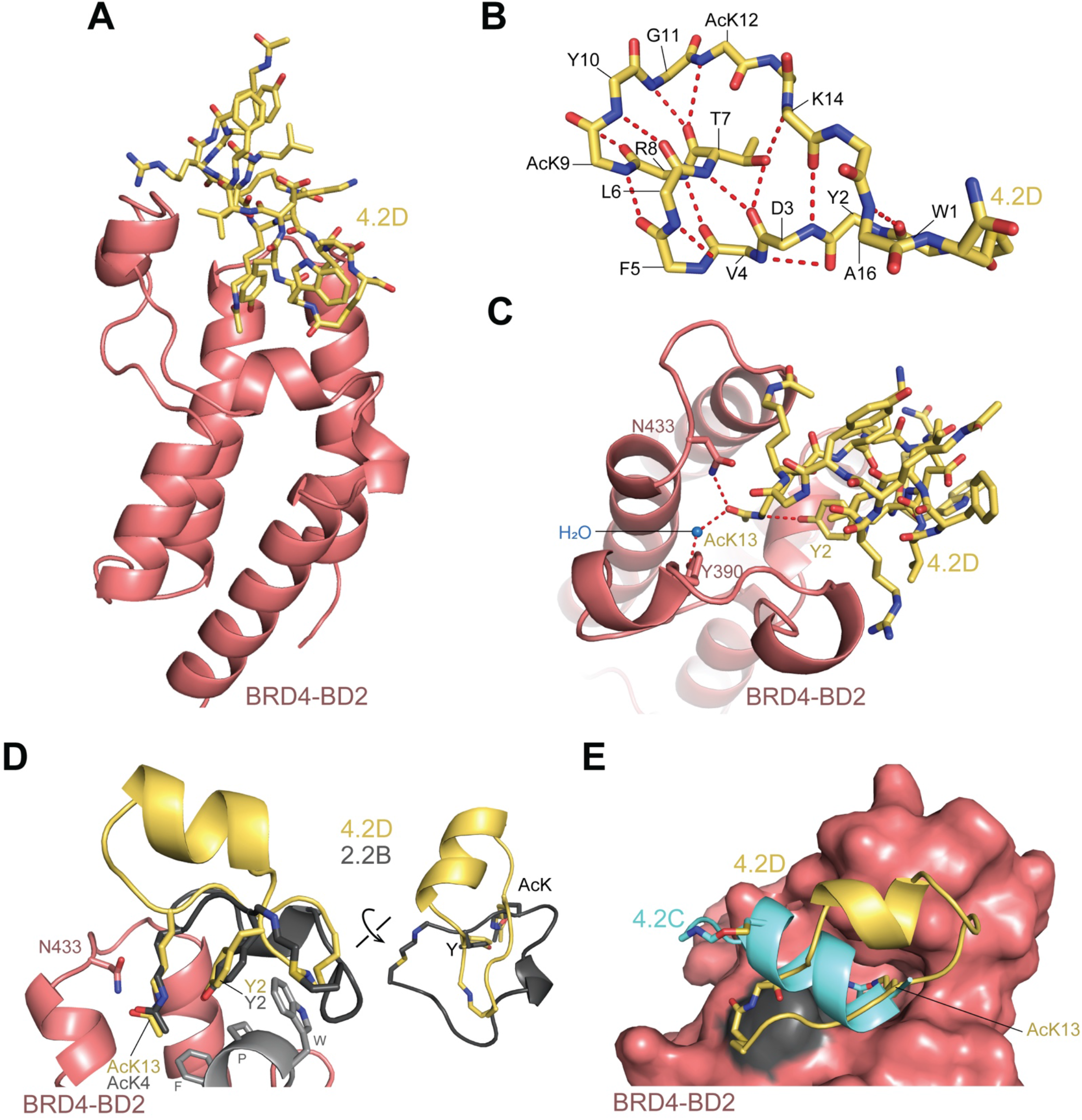
Structure of cyclic peptide 4.2D in complex with BRD4-BD2. The peptide in shown in yellow and BRD4-BD2 is shown in salmon. **A**. Ribbon/stick representation of the complex formed between **4.2D** and BRD4-BD2 (1.7 Å resolution; PDB ID: 8CV6). **B**. Backbone of **4.2D** in the structure of BRD4-BD2. Internal hydrogen bonds within the mainchain and between the sidechain hydroxyl group of Thr7 and the mainchain of the peptide are represented as red dashed lines. **C**. Key interactions between **4.2D** and BRD4-BD2. The hydrogen bonds formed between AcK13 of **4.2D** and N443 and Y390 (mediated via a water molecule) in the binding pocket of BRD4-BD2 are indicated by red dashed lines. The hydrogen bond formed between the sidechain of AcK13 and Y2 of **4.2D** is also displayed. **D**. Overlay of the structure of **2.2B** (dark grey) on the structure of **4.2D** showing the positioning of the structurally conserved Tyr in both peptides. The conserved Tyr is sandwiched between the primary AcK residue that enters the BD-binding pocket in both **4.2D** and **2.2B** and the WPF shelf (depicted in light grey) of the BDs with which it directly interacts. The inset shows a rotated view of the two peptides emphasising the very different backbone structures of the two peptides. **D**. Relative alignment of the α-helices observed in **4.2E** and **4.2C**. Both the AcK residue that engages the binding pocket and the cyclisation moieties are shown.

Several peptides that we have recently discovered feature α-helices that are oriented on the BD target in a similar manner (as seen in the overlay of the α-helical BET-BD binding cyclic peptides **3.2B, 4.2A**, and **4.2C** in **Figure 6G**); however, the helix in **4.2D** lies at a significantly different angle (**Figure 7E**). In addition, the pocket-binding AcK residue does not lie in the helix, but rather in an extended part of the peptide (**Figure 7E**), indicating that a variety of peptide topologies and orientations are consistent with high-affinity BD binding.

Peptide **4.2D** demonstrated exceptional affinity for the BD against which it was selected compared to the remaining five BDs (**Supplementary Table 1**). Thus, we also determined the structure of the peptide in complex with BRD3-BD2 (1.5 Å, PDB ID: 8CV5, **Supplementary Figure 4A, Supplementary Table 7**, *K*_*D*_ = 20 nM; ∼60-fold weaker binding compared to BRD4-BD2) to uncover any structural differences that contribute to the observed selectivity. The structure is highly similar to the structure of **4.2D** bound to BRD4-BD2 (**Supplementary Figure 4A** and **4B**). The RMSD over 109 Cα atoms across the two complexes is 0.35 Å and only one BD residue within 4 Å of the peptide in either complex differs between the two paralogues: a glutamate in BRD3-BD2 (E344) that is a glycine in BRD4-BD2 (**Supplementary Figure 4C**). This residue can form a salt bridge with Arg8 in **4.2D**, which, if anything would be envisaged to increase the affinity of **4.2D** for BRD3-BD2, the converse of what is observed. Again, therefore the basis for this significant difference in affinity is not apparent from our structural data.

## DISCUSSION

### A wide range of binding modes that feature conserved elements

Here we report the characterisation of macrocyclic peptides that were selected in RaPID screens against the BD2 domains of the BET proteins BRD2, BRD3 and BRD4. Typically, the peptides form highly organised structures with substantial intramolecular hydrogen bonding networks. Indeed, all peptides described previously and most here display an extensive network of single, bifurcated, and even trifurcated internal hydrogen bonding. By contrast, peptide **2.2B** displays a relatively open structure with only five hydrogen bonds, all in one half of the peptide, suggesting that a compact structure is not a prerequisite for tight binding.

In line with our previous observations (Patel et al., 2020), these peptides display a diverse range of secondary structural motifs, including α-helices (**4.2D**) and β-hairpins (**4.2E**); others display irregular backbones with no elements of regular secondary structure, despite being highly ordered (**2.2E** and **2.2B**). Interestingly, although it lacked regular secondary structure, peptide **2.2E** shares a conserved motif with a peptide from our previous work, peptide **3.1C**. These two peptides overlay closely in the shared six-residue motif, but with no structure or sequence conservation in the other half of the peptides, demonstrating that different macrocyclic scaffolds can present similar binding epitopes. Indeed, not only do distinct folds present similar motifs, but we also observed the converse: the same secondary structural element docking onto a BD in distinct manners [*e*.*g*., the helical peptides **4.2C** vs **4.2D** or the β-sheet peptides **4.2E** vs **3.1B** from (Patel et al., 2020)], further underscoring the diversity represented in RaPID peptide libraries.

Despite this high structural diversity, some conserved interactions were observed. The most prominent feature of all these diverse peptide ligands is the binding of an AcK residue in the canonical binding pocket of the BDs. We previously observed that macrocyclic ligands for BD2 domains (namely BRD3-BD2 and BRD4-BD2) were often α-helical and displayed a diagonal entry angle of the AcK residue into the binding pocket (Patel et al., 2020). This angle contrasted with the vertical insertion observed both for native AcK ligands and for β-hairpin-based ligands of BD1 domains. In the current work, an AcK in the helical **4.2D** peptide also addresses the AcK pocket via a diagonal entry angle. Peptide **4.2E**, however, is the first β-hairpin that we have observed to engage with the BD2 through this non-canonical AcK entry angle, again emphasizing that favourable interactions can be implemented by distinct peptides in a wide range of structural contexts.

We also noted that all peptides are characterised by the presence of a hydrophobic residue that directly contacts the aliphatic sidechain of the interacting AcK, effectively forming a wedge at the top of the binding pocket (*e*.*g*., Tyr2 in **2.2B** and Trp5 in **2.2E**) and perhaps reducing the conformational entropy of the long AcK sidechain to promote stronger binding. Further examination of our full set of BD-peptide complex structures shows Leu/Ile/Trp/Tyr/His residues in a comparable position in nine of the structures (though in a variety of distinct sequence settings). In cases where the sidechain has a polar moiety (*e*.*g*., the Tyr hydroxyl group), water mediated hydrogen bond networks often connect that moiety with the NE of the nearby AcK. This location, which is adjacent to the WPF shelf, is thus both a key element of substrate-BD interactions and a favoured target site for high-affinity ligands.

### Selectivity without structural differences

Perhaps most strikingly, we discovered peptides (*e*.*g*., **3.2D** and **4.2D**) that displayed significant selectivity between BD2 paralogues. These affinity differences demonstrate that even within the series of highly homologous BD2 domains, selectivities of ≥ 50-fold can be achieved, and furthermore that there are multiple peptide scaffolds that can support such specificities. Surprisingly, however, although structures were obtained for several complexes of BDs and peptides, we were unable to account for these differences in specificity. We made a similar observation for peptide 2.2B, which bound BD1 domains with significantly higher affinity than BD2 domains, but for reasons that remain cryptic. We speculate that differences in conformational dynamics between BD paralogues might play an important role in determining binding affinities. Indeed, a previous study has shown that differences in binding-loop dynamics between the BDs of BRD7 and BRD9 influences interactions with small molecules (Wang et al., 2022).

In summary, our study has revealed a range of new macrocyclic peptides that recognize BET BD2 bromodomains, adding to an already diverse collection of structures observed in our previous work. These peptides can be based on diverse structural scaffolds and use very different binding modes to the BD2 domains to achieve high potency and substantial selectivity, emphasizing the diverse chemical space available for exploitation in drug discovery (although cell permeability of such peptides remains an issue to be addressed). We have also demonstrated that these macrocycles can achieve significant paralogue-level selectivity *in vitro*, a relatively elusive property in the BET bromodomain space. Further studies will be required to understand the origins of this selectivity, which might in the longer term provide insight into routes to rationally develop paralogue selective BET inhibitors.

## DATA AVAILABILITY

Coordinates and structure factors for all structures reported in this study have been deposited to the PDB under the following accession codes: 8CV7 (BRD2-BD2:**2.2E**), 8DNQ (BRD2-BD1:**2.2B**), 8CV4 (BRD4-BD2:**4.2E**), 8CV6 (BRD4-BD2:**4.2D**), and 8CV5 (BRD3-BD2:**4.2D**).

## ACKNOWLEDGEMENTS

We acknowledge Sydney Analytical for access to the SPR infrastructure. This research was conducted using the MX1 and MX2 beamlines at the Australian Synchrotron (a part of ANSTO) and made use of the Australian Cancer Research Foundation (ACRF) Eiger 16M detector. J.P.M., L.J.W., and R.J.P. received funding from the National Health and Medical Research Council (APP 1161623). R.J.P. is supported by an Investigator Grant (APP 1174941) from the National Health and Medical Research Council. This work was also supported by the Francis Crick Institute which receives its core funding from Cancer Research UK (CC2030), the UK Medical Research Council (CC2030), and the Wellcome Trust (CC2030).

For the purpose of Open Access, the authors have applied a CC BY public copyright licence to any Author Accepted Manuscript version arising from this submission.

## AUTHOR CONTRIBUTIONS

Conceptualization: C.F., K.P., L.J.W., T.P., H.S., R.J.P., J.P.M.; Data curation: C.F., K.P., L.J.W., M.C., A.N.; Investigation: C.F., K.P., L.J.W., M.C.; Methodology: C.F., K.P., L.J.W., M.C.; Resources: R.J.P, H.S., J.P.M.; Supervision: R.J.P., J.P.M.; Writing – original draft: C.F., K.P., L.J.W., T.P., J.P.M.; Writing – review & editing: C.F., K.P., L.J.W., M.C., R.J.P., J.P.M.

## METHODOLOGY

### Protein expression and purification

The bromodomains from human BRD2 (BD1: 65-194; BD2: 347-455), BRD3 (BD1: 25-147; BD2: 307-419), and BRD4 (BD1: 42-168; BD2: 348-464) were cloned for bacterial expression into the pGEX-6P and pQE80L-NAvi plasmids for expression as *N*-terminal GST-fusion proteins and *N*-terminally His-tagged and biotinylated proteins, respectively.

All bromodomain-containing proteins, except BRD4-BD1, were transformed into competent BL21(DE3) Escherichia *coli* (*E. coli*) cells. Overnight saturated starter cultures, grown from single colonies from fresh transformation plates, were used to inoculate LB expression cultures (1:100). Expression cultures were grown at 37 °C, with shaking at 150 rpm, until cultures reached mid-log phase (OD600 of ∼0.6-0.8). At this point, cultures were cooled to room temperature and protein expression was induced by the addition of 250 µM isopropyl β-d-1-thiogalactopyranoside (IPTG) and supplemented with 200 µM biotin for pQE80L-NAvi constructs. Expression cultures were transferred to 18 °C for expression for a further ∼20–24 h before harvesting. Biotinylated BRD4-BD1 was expressed using Rosetta2(DE3) *E. coli* cells. To this end, starter cultures, as described above, were used to inoculate autoinduction expression cultures, which were grown at 37 °C with shaking at 150 rpm. Upon reaching mid-log phase, cultures were cooled to room temperatures, supplemented with 200 µM biotin and grown for a further 20-24 h at 18 °C before harvesting. All media were supplemented with the appropriate antibiotics and cultures were harvested via centrifugation at 5000 ×g at 4 °C for 25 min and cell pellets were stored at –20 °C.

Cell pellets expressing biotinylated proteins for SPR were resuspended in a buffer composed of 50 mM Tris pH 8.0, 500 mM NaCl, 20 mM imidazole, 5 mM β-mercaptoethanol (β-ME), 0.1% Triton X-100, 1× cOmplete EDTA-free protease inhibitor, 10 µg mL^-1^ DNase I, 10 µg mL^-1^ RNase, and 100 µg mL^-1^ lysozyme and lysed via sonication. Centrifugation at 18,000 ×g for 30–60 min separated the soluble from insoluble fraction. The proteins (present in the soluble fraction) were initially purified *via* immobilised nickel ion affinity chromatography using a 1-mL HisTrap column. The bound protein was eluted using a 20–250 mM imidazole gradient elution. The protein-containing fractions were pooled and concentrated to a small volume prior to loading on a HiLoad 16/600 Superdex 75 column. Proteins were eluted from the column using 20 mM HEPES pH 7.5, 150 mM NaCl, and 1 mM tris(2-carboxyethyl)phosphine (TCEP). Protein-containing eluates were pooled and either aliquoted directly or concentrated before aliquoting. Aliquots were snap frozen in liquid nitrogen and stored at –80 °C. Protein purification was analysed by SDS-PAGE and monitoring UV absorbance at 280 nm.

Cell pellets expressing GST-fused proteins for crystallography were lysed in a buffer composed of 50 mM Tris pH 7.2, 500 mM NaCl, 5 mM β-ME, 0.1% Triton X-100, 1× cOmplete EDTA-free protease inhibitor, 10 µg mL^-1^ DNase I, 10 µg mL^-1^ RNase, and 100 µg mL^-1^ lysozyme via sonication. Lysate was clarified via centrifugation at 18,000 ×g for 30–60 min and the soluble fraction was subjected to reduced glutathione (GSH)-affinity chromatography using a 5-mL GSTrap column. Bound protein was eluted stepwise with 50 mM Tris pH 7.2, 150 mM NaCl, 10 mM GSH, and 5 mM β-ME. Protein-containing fractions were pooled and incubated overnight at 4 °C with HRV-3C protease to enable cleavage of the GST-tag. The cleaved protein was concentrated prior to size exclusion chromatography using a HiLoad 16/600 Superdex 75 column. The protein was eluted from the column in a buffer comprising 20 mM HEPES pH 7.5, 150 mM NaCl, and 1 mM TCEP. Fractions containing the desired protein were pooled and concentrated before aliquoting. Protein was concentrated to 10–15 mg mL^-1^. Protein aliquots were snap frozen in liquid nitrogen and stored at – 80 °C.

### RaPID screening

Transcribed mRNA libraries were produced using T4 RNA polymerase from DNA ljbraries with the following sequence: TAATACGACTCACTATAGGGTTGAACTTTAAGTAGGAGATATATCCATG(NNK)m= 3-7ATG(NNK)n=4-7TGTGGGTCTGGGTCTGGGTCTTAGGTAGGTAGGCGGAAA

Library mRNA was ligated to a puromycin-PEG-DNA splint using T4 RNA ligase following standard reaction conditions. For the first selection round libraries were mixed in the following proportions: 0.01425(m=3,n=4): 0.45(m=4,n=4): 10(m=4,n=5): 10(m=5,n=5): 7.5(m=5,n=6): 7.5(m=6,n=6): 7.5(m=6,n=7): 7.5(m=7,n=7).

RaPID screens were carried out as previously described (Patel et al., 2020). Briefly, puromycin-ligated randomised mRNA libraries were in vitro translated (30 min, 37 °C then 12 min, 25 °C) using a custom transcription/translation mixture lacking methionine and 10-formyl-5,6,7,8-tetrahydrofolic acid and containing additional 12.5 µM ClAc-L-Trp-tRNA^fMet^_CAU_ and 25 µM AcK-tRNA^Asn^_CAU_ prepared as described previously (ref 1 and 2 from the SI of our PNAS paper). First-round translations were carried out on a 150-µL scale, and all subsequent rounds on a 5-µL scale. Following translation, 200 mM EDTA, pH 8.0 (15 µL) was added and reverse transcription with M-MLV RTase, RNase H minus (Promega) performed. Subsequently blocking buffer was added (50 mM HEPES, 150 mM NaCl, 2 mM DTT, 0.1% Tween-20, 0.2% (w/v) acetylated bovine serum albumin, pH 7.5) and the library incubated with magnetic streptavidin bead-immobilised BRD2-BD2 (Promega) (200 nM, 30 min). Three washing steps were performed (156 µL ice-cold 50 mM HEPES, 150 mM NaCl, 2 mM DTT, 0.1% Tween-20, pH 7.5, 3 × 5 min), after which 400 µL PCR solution was added and retained peptide-mRNA/DNA hybrids eluted from the beads by heating (95 °C, 5 min). Library enrichment was determined by quantitative real-time PCR relative to standards and the input DNA library using primers T7g10M_F46 and CGS3-CH.R22. Enriched pools were amplified using the same primers and used as the input DNA for subsequent selection rounds.

T7g10M_F46 -TAATACGACTCACTATAGGGTTGAACTTTAAGTAGGAGATATATCC CGS3-CH.R22 – TTTCCGCCTACCTACCTAAGAC

Enriched pools from rounds 3-5 were also used to prepare double indexed libraries (Nextera XT indices) for sequencing on a MiSeq platform (Illumina) using a v3 chip as single 151 cycle reads. Sequencing reads were converted into peptide sequences and ranked by total read number (**Supplementary Table 1**).

### Peptide synthesis

All peptides were assembled by stepwise fluorenylmethyloxycarbonyl (Fmoc)-based solid-phase peptide chemistry on a Syro I automated synthesiser (Biotage). Peptides were synthesised as *C*-terminal amides using NovaPEG Rink Amide resin (novabiochem). Couplings were performed with DIC/Oxyma (1:1) and 5 equivalents (equiv.) of each amino acid. Upon coupling of the *N*-terminal Fmoc-Trp(Boc)-OH, the Fmoc group was removed (20 % (v/v) piperidine in DMF) and the resin was incubated in DMF with choloroacetic acid (4 equiv.), Oxyma (4 equiv.) and Oxyma (4 equiv.). Resin was washed with DMF and CH_2_Cl_2_ (each 5 times) and linear peptides were cleaved from resin by treatment with an acidic cocktail of trifluoroacetic acid (TFA):triisopropyl silane (*i*Pr_3_SiH):H_2_O (90:5:5 v/v/v) for 2 h at room temperature with agitation. The cleavage solution was filtered, concentrated in vacuo and crude linear peptide was precipitated with ice-cold diethyl ether and dried under a gentle nitrogen flow. All crude linear peptides, except peptides 2.2A, 2.2B and 2.2C, were resuspended in 5 vol% *i*Pr_2_NEt in DMF to a final concentrated of 5 mM to allow for macrocyclisation. Peptides 2.2A, 2.2B and 2.2C were resuspended in DMSO and the pH was raised to >8 using triethylamine to allow cyclisation. Upon completion of the cyclisation (1 h, rt), peptides were reacidified with TFA for downstream purification by reverse-phase high-performance liquid chromatography using a Xbridge BEH C18 column (300 Å – Waters) on a Waters HPLC system (Solvent A: 0.1% TFA in H_2_O, Solvent B: 0.1% TFA in acetonitrile). Purified peptides (as determined by mass spectrometry) were reconstituted in DMSO and their concentrations determined from their absorbance at 280 nm.

### Surface plasmon resonance (SPR)

SPR measurements were conducted using a Biacore™ T200 (GE Healthcare) with a running buffer comprised of 10 mM HEPES pH 7.4, 150 mM NaCl, 0.01% (w/v) Tween-20. The CAP chip (GE Healthcare) was used to immobilise biotinylated BET bromodomains with a target density of 1000-1500 response units (RU). Regeneration of the chip was performed between cycles as per the manufacturer’s protocol. All experiments were performed at 4 °C in single cycle kinetics mode. Data analysis was conducted using the Biacore™ Insight Evaluation Software.

### X-ray crystallography

X-ray crystallography of the bromodomain-peptide complexes was performed using the sitting-drop vapour-diffusion technique. The peptide (1.5 molar equiv.) was added to the purified BD proteins (10-15 mg mL-1) and incubated on ice for 30 min to allow for complex formation. Initial crystallisation trials were conducted using the commercially available 96-well screens PACT (Molecular Dimensions), JCSG+ (Molecular Dimensions), PEGion (Hampton Research), PEGRx (Hampton Research), and Index HT (Hampton Research). A Mosquito crystallisation robot (SPT Labtech) dispensed the protein-peptide mixtures into MRC two-drop chamber, 96-well crystallisation plates (Hampton Research) and each condition was screened at a protein to precipitant ratio of 1:1 or 2:1 with a final drop volume of 300 nL. All 96-well plates were incubated at 18 °C and crystals generally appeared after days to weeks. 10 vol% glycerol in the mother liquid from which the crystals were grown was added to the crystal to allow for cryoprotection prior to plunge freezing in liquid nitrogen.

The Macromolecular Crystallography MX1 and MX2 beamlines (at 100 K and wavelength of 0.9537 Å) were used at the Australian Synchrotron for X-ray diffraction data collection of frozen crystals (Aragão et al., 2018; Cowieson et al., 2015). XDS was used for data integration and further processing was performed using the CCP4i suite (Potterton et al., 2003; Winn et al., 2011). Indexing, scaling, and merging of the data was performed using AIMLESS and the initial phases were calculated by the molecular replacement program Phaser MR using existing X-ray crystal structures of BRD-BD proteins as the molecular replacement models (PDB IDs: 4UYF (BRD2-BD1), 3ONI (BRD2-BD2), 3S92 (BRD3-BD2), and 5UVV (BRD4-BD2)) (Filippakopoulos et al., 2010; Gosmini et al., 2014). COOT was used for manual model building and for iterative rounds of manual building followed by refinement using Phenix (Adams et al., 2010; Emsley et al., 2010; Murshudov et al., 2011). Structure diagrams were generated using PyMol and protein:peptide interfaces were analysed using PDBePISA. The data collection and refinement statistics for all structures described in this study are outlined in **Supplementary Table 2, Supplementary Table 3**, and **Supplementary Tables 5-7**.

