## SupplementaryData for "Discovery and characterization of cyclic peptides selective for the *C*-terminal bromodomains of BET family proteins"

### SUPPLEMENTARY FIGURES

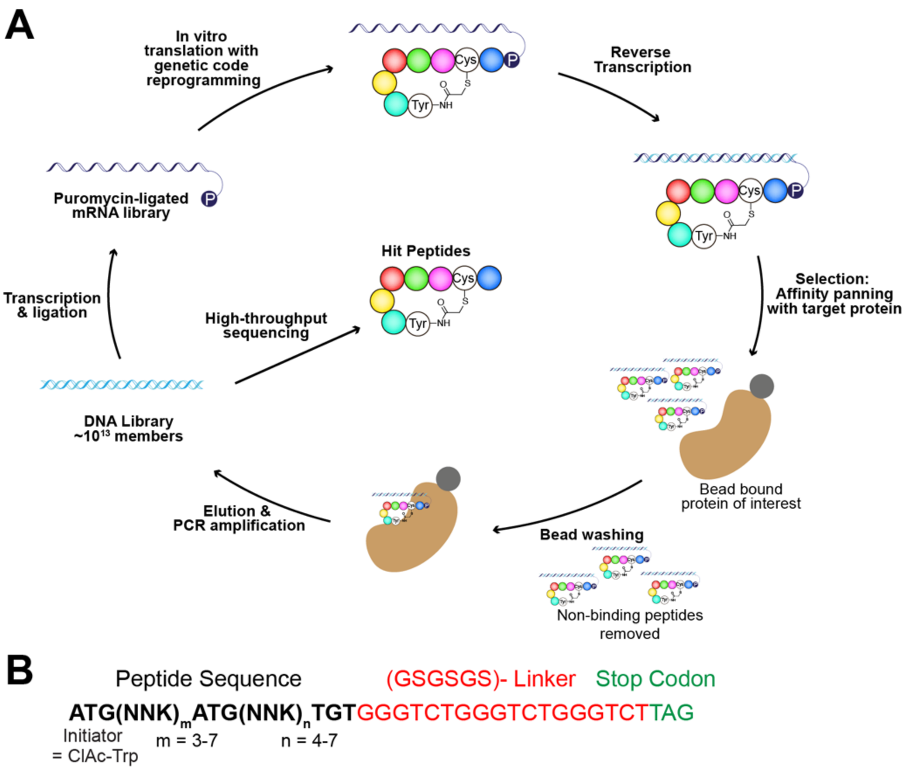


| 1st | 2nd: U | 2nd: C | 2nd: A | 2nd: G | 3rd |
|---|---|---|---|---|---|
| U | Phe | Ser | Tyr | Cys | U |
| | | | | | C |
| | | | | | A |
| | Leu | Ser | Stop | Trp | G |
| C | Leu | Pro | His | Arg | U |
| | | | | | C |
| | | | | | A |
| | Leu | Pro | Gln | Arg | G |
| A | Ile | Thr | Asn | Ser | U |
| | | | | | C |
| | | | | | A |
| | **KAc** | Thr | Lys | Arg | G |
| G | Val | Ala | Asp | Gly | U |
| | | | | | C |
| | | | | | A |
| | Val | Ala | Glu | Gly | G |

**Supplementary Figure 1. Overview of the RaPID system. A.** Schematic of the RaPID selection scheme. **B.** Library design and codon assignment used in the RaPID selections.

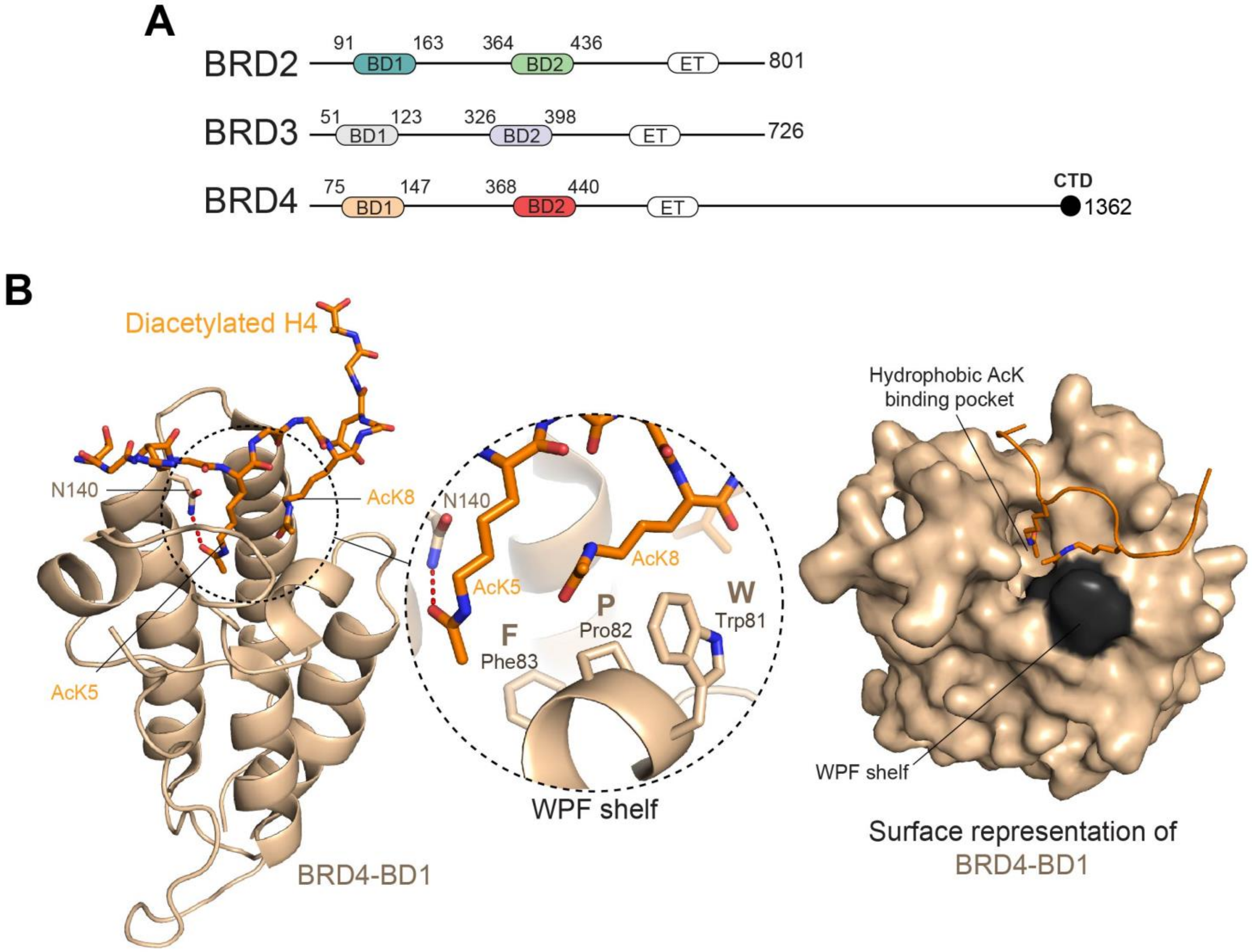


**Supplementary Figure 2. Domain architecture of the Bromodomain and ExtraTerminal domain (BET) family of proteins. A.** Domain topology of human BRD2, BRD3, and BRD4. The first (BD1) and second (BD2) bromodomains, extraterminal domains (ET), and C-terminal domains (CTD) are indicated. Residue ranges of the BDs are indicated, and the colouring shown for each individual BD is used throughout the text. **B.** The X-ray crystal structure of BRD4-BD1 (*wheat*) in complex with a diacetylated histone H4 peptide (*orange*, PDB ID: 3UVW). *Left*: Ribbon representation of the BRD4-BD1:diacetylated histone H4 complex. The conserved asparagine (Asn140) and the *N*-terminal AcK residue from the diacetylated histone H4 motif (AcK5) with which Asn140 makes a hydrogen bond are displayed as sticks and the hydrogen bond formed between them is indicated by the orange dashed line. A close-up of the WPF shelf that forms significant van der Waals interactions with *C*-terminal AcK (AcK8) of the diacetylated motif in histone H4 is shown in the diagram enclosed in the dashed circle. *Right*: Surface representation of BRD4-BD1 showing the hydrophobic cavity of the BD that is occupied by the *N*-terminal AcKs of diacetylated motifs. The WPF shelf with which the *C*-terminal AcKs of diacetylated motifs interact is coloured in *dark grey*.

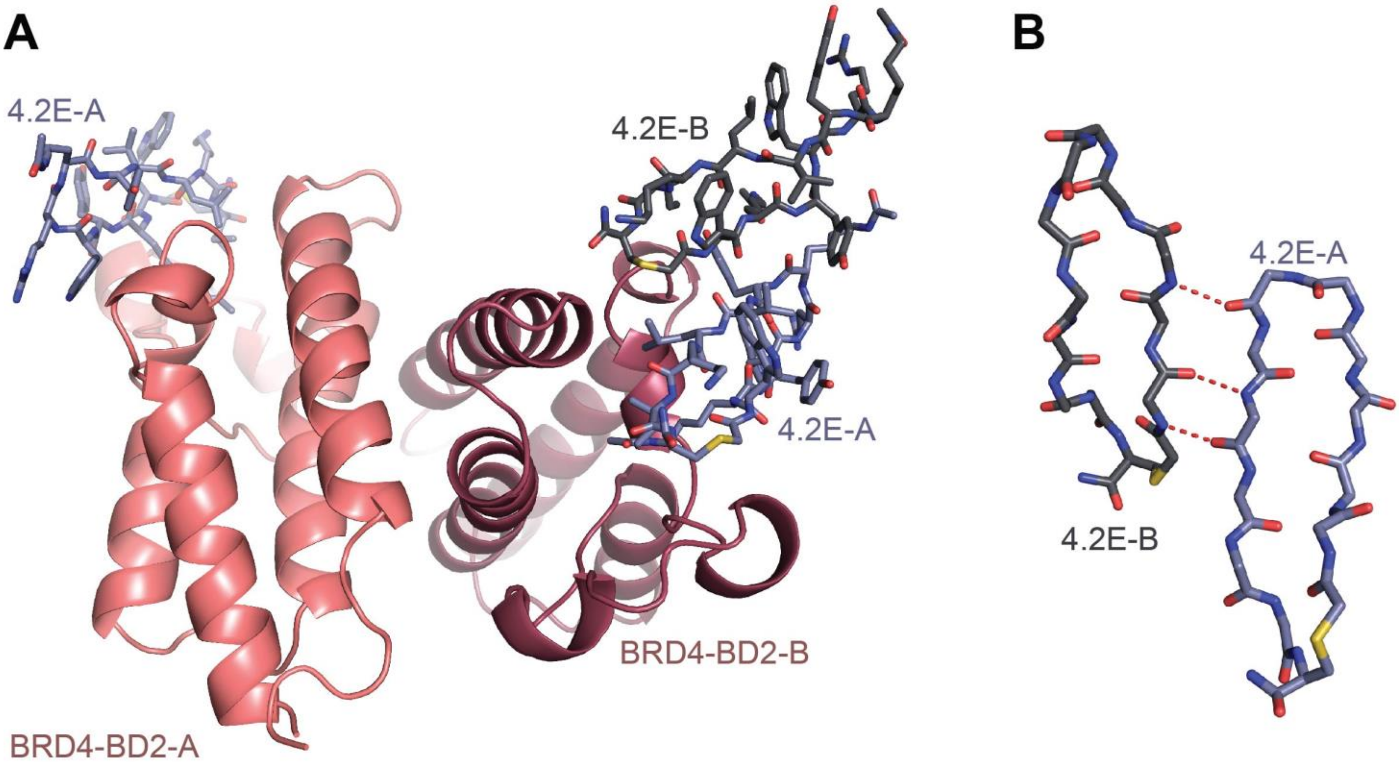


**Supplementary Figure 3. Asymmetric unit of the BRD4-BD2:4.2E complex.** **A**. The asymmetric unit shows two copies of BRD4-BD2, each bound in an identical fashion to a molecule of **4.2E** (**4.2-A**). An additional peptide, **4.2E-B**, is observed interacting with the **4.2-A** molecule that is bound to BRD4-BD2-B in the asymmetric unit. **B**. Inter-mainchain hydrogen bonding between **4.2E-A** and **4.2E-B** (red dashed lines).

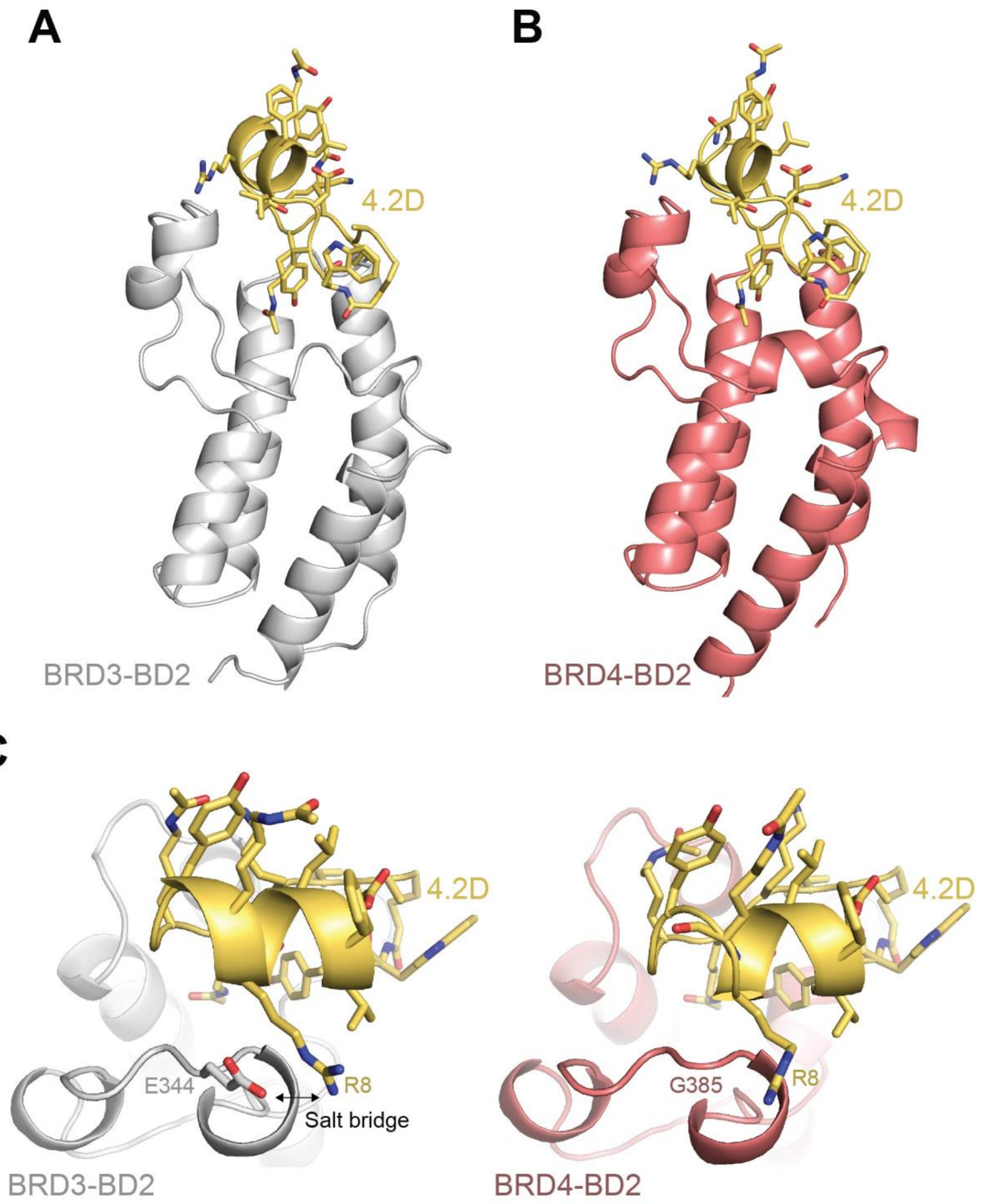


**Supplementary Figure 4. Comparison of the BRD3-BD2:4.2D and BRD4-BD2:4.2D complexes.** The peptide is shown in yellow, BRD3-BD2 is shown in grey, and BRD4-BD2 is shown in salmon. **A**. Ribbon representation of the structure of BRD3-BD2 bound of **4.2D** (1.5 Å resolution; PDB ID: 8CV5). **B**. Ribbon representation of the structure of BRD4-BD2 in complex with BRD4-BD2 shown in the same orientation as the structure in **A**. An overlay of α-carbons of both structures shown in **A** and **B** has an RMSD of 0.32 Å). **C**. *Left panel:* Close-up view of the BRD3-BD2:**4.2D** interaction. The sole residue, E344, in BRD3-BD2 that differs from BRD4-BD2 within the binding sphere of the interaction is displayed. This residue forms a salt bridge with R8 of **4.2D**. *Right panel*: Close-up view of the BRD4-BD2:**4.2D** interaction. The position of the sole residue, G385, in BRD4-BD2 that differs from BRD3-BD2 within the binding sphere of the interaction is shown. This residue does not form any contacts with **4.2D**.

### SUPPLEMENTARY DATASET

#### Supplementary Dataset 1.

Sequences recovered from each round of RaPID selection against BRD2-BD2 (attached as a separate spreadsheet).

### SUPPLEMENTARY TABLES

**Supplementary Table 1.** Peptide details and dissociation constants for the cyclic peptides selected for study from RaPID selections against BRD2-BD2, BRD3-BD2, and BRD4-BD2.

| RaPID Selection | Peptide | Sequence | Enrichment | $K_D$ (nM)[a] | | | | | |
|---|---|---|---|---|---|---|---|---|---|
| | | | | BRD2-BD1 | BRD3-BD1 | BRD4-BD1 | BRD3-BD1 | BRD3-BD2 | BRD4-BD2 |
| BRD2-BD2 | 2.2A | WNGWW**K**IP**K**Q**K**C | 2.3% | 0.4 ± 0.2 | 0.4 ± 0.1 | 0.6 ± 0.3 | 0.2 ±0.08 | 2 ± 3 | 2 ±0.6 |
| BRD2-BD2 | 2.2B | WYS**K**KYA**K**WWTVYPC | 1.4% | 0.4 ± 0.2 | 0.4 | 0.5 | 20 | 50 | 200 |
| BRD2-BD2 | 2.2C | WHYWLLR**K**GKIHKLRC | 1.3% | nd[c] | nd[c] | nd[c] | nd[c] | nd[c] | nd[c] |
| BRD2-BD2 | 2.2D | WSGYW**K**IP**K**QLC | 1.2% | 1 ± 0.4 | 0.9 | 2 | 0.4 | 8 | 6 |
| BRD2-BD2 | 2.2E | WYGYW**K**VP**K**RKC | 0.6% | 0.8 ± 0.1 | 0.7 | 0.7 | 2 | 60 | 3 |
| BRD3-BD2 | 3.2A | W**K**KAILPG**K**ILKTLHIC | 3.1% | [b] | [b] | 9 ± 20 | [b] | [b] | [b] |
| BRD3-BD2 | 3.2B | WSWLCK**K**YNLIHC | 2.6% | 20 ± 20 | 30 ± 30 | 20 ± 8 | 40 ± 50 | 10 ± 8 | 30 ± 20 |
| BRD3-BD2 | 3.2C | W**K**QWK**K**YGLKIC | 2.0% | [b] | [b] | [b] | 90 ± 50 | 30 ± 10 | 10 ± 10 |
| BRD3-BD2 | 3.2D | WISYQ**K**YRLW**K**C | 0.2% | [b] | [b] | [b] | 300 ± 8 | 70 ± 8 | 100 ± 20 |
| BRD3-BD2 | 3.2E | WNV**K**YW**K**KYNIKC | 0.1% | [b] | [b] | [b] | 20 ± 2 | 6 ± 3 | 3 ± 2 |
| BRD3-BD2 | 3.2F | WLYYQ**K**W**K**LF**K**C | 0.1% | [b] | 1000 ± 1000 | [b] | 7 ± 2 | 2 ± 3 | 4 ± 2 |
| BRD3-BD2 | 3.2G | WLFY**KK**WRLY**K**C | 0.2% | [b] | [b] | [b] | 9 ± 7 | 2 ± 0.1 | 7 ± 0.8 |
| BRD4-BD2 | 4.2A | W**K**NWCWL**K**R**K**LLLRC | 20.0% | 50 ± 30 | 3 ± 1 | 30 ± 30 | 100 ± 100 | 20 ± 10 | 30 ± 10 |
| BRD4-BD2 | 4.2B | WAYHTIRL**K**WR**W**PSISC | 9.5% | 20 ± 0.03 | [b] | [b] | 3 ± 6 | 100 ± 10 | 1 ± 0.04 |
| BRD4-BD2 | 4.2C | W**K**GYLCLR**K**RIQRTYNC | 4.6% | [b] | [b] | [b] | 20 ± 30 | 20 ± 100 | 40 ± 40 |
| BRD4-BD2 | 4.2D | WYDVFLTR**K**YG**KK**KVAC | 3.8% | 90 ± 20 | 10 ± 7 | 40 ± 50 | 20 ± 7 | 20 ± 3 | 0.3 ± 0.05 |
| BRD4-BD2 | 4.2E | W**K**YWR**K**YVLKIC | 2.5% | [b] | 30 ± 10 | [b] | 2 ± 0.4 | 20 ± 0.7 | 0.9 ± 0.2 |
| BRD4-BD2 | 4.2F | WLLFYNH**K**LIRRLL**K**C | 1.5% | [b] | 1000 ± 1000 | 1000 ± 1000 | 200 ± 80 | 20 ± 3 | 40 ± 20 |
| BRD4-BD2 | 4.2G | W**K**SWG**K**HFLLVC | 1.0% | 400 ± 400 | 400 ± 200 | [b] | 200 ± 30 | 20 ± 0.8 | 20 ± 0.1 |

[a]The $K_D$ values are given as the geometric mean of a minimum of three independent measurements. We estimate the uncertainty in each $K_D$ value to be ~25%.
[b]No binding observed (meaning any interaction must have a $K_D$ >1 μM).
[c]No data (nd) due to non-specific binding of the peptide to the SPR sensor chip.
[d]Coloured boxes in the peptide detail section indicate the peptides from each selection that are unique to this study. Details of previously studied peptides (unhighlighted) are also given. Affinities of the peptides for their cognate BDs are highlighted by coloured boxes in the $K_D$ values section.

**Supplementary Table 2.** Data collection and refinement statistics for the crystal structure of BRD2-BD2 in complex with **2.2E** (PDB ID: 8CV7) solved using molecular replacement.

| | BRD2-BD2:2.2E |
|---|---|
| **Data collection** | |
| Space group | P 1 21 1 |
| Cell dimentions | |
| *a, b, c* (Å) | 52.95, 52.42, 55.99 |
| *α, β, γ* (°) | 90.00, 96.67, 90.00 |
| Resolution (Å) | 1.6 (1.6-1.63) |
| $R_{merge}$ | 0.054 (0.427) |
| $I/\sigma I$ | 18.3 (4.2) |
| CC(1/2) | 0.999 (0.933) |
| Completeness (%) | 98.3 (97.0) |
| Redundancy | 7.0 (7.0) |
| **Refinement** | |
| Resolution (Å) | 1.6 |
| No. reflections | 39545 |
| $R_{work}/R_{free}$ | 0.1756/0.1983 |
| No. atoms | 2330 |
| Protein | 1812 |
| Ligand/ion | 248 |
| Water | 270 |
| *B*-factors | 23.94 |
| R.m.s. deviations | |
| Bond lengths (Å) | 0.0069 |
| Bond angles (°) | 1.0250 |

*Values in the parantheses are for the highest resolution shell.
All data were collected on a single crystal.

**Supplementary Table 3.** Data collection and refinement statistics for the crystal structure of BRD2-BD1 in complex with **2.2B** (PDB ID: 8DNQ) solved using molecular replacement.

| | **BRD2-BD1:2.2B** |
|---|---|
| **Data collection** | |
| Space group | P 1 21 1 |
| Cell dimentions | |
| *a, b, c* (Å) | 41.68. 54.37, 76.93 |
| *α, β, γ* (°) | 90.00, 93.85, 90.00 |
| Resolution (Å) | 1.84 (1.84-1.88) |
| $R_{merge}$ | 0.095 (0.701) |
| $I/\sigma I$ | 6.9 (1.3) |
| CC(1/2) | 0.996 (0.733) |
| Completeness (%) | 98.2 (97.4) |
| Redundancy | 3.7 (3.6) |
| **Refinement** | |
| Resolution (Å) | 1.84 |
| No. reflections | 29500 |
| $R_{work}/R_{free}$ | 0.2115/0.2614 |
| No. atoms | 2482 |
| Protein | 1950 |
| Ligand/ion | 330 |
| Water | 196 |
| *B*-factors | 22.00 |
| R.m.s. deviations | |
| Bond lengths (Å) | 0.0070 |
| Bond angles (°) | 0.8270 |

*Values in the parantheses are for the highest resolution shell.
All data were collected on a single crystal.

**Supplementary Table 4.** Sequence alignment of the 500 most enriched sequences from the BRD2-BD2, BRD3-BD2, and BRD4-BD2 RaPID selections.

| | | |
|---|---|---|
| BD32 0.07% | WIYY RXWRLYXCGA GSGS* | 19 |
| BD32 0.03% | WIYY RXWRLYXCGA GSGA* | 19 |
| BD32 0.05% | WIYY RXWRLYXCGS GSGA* | 19 |
| BD32 0.03% | WIYY RXWRLYXCGS GAGA* | 19 |
| BD42 0.01% | WIYY QXWRIKXCGA GSGS* | 19 |
| BD42 0.01% | WIYW RXYRLRXCGA GSGS* | 19 |
| BD42 0.01% | WLYY KXWRLHXCGA GSGS* | 19 |
| BD42 0.01% | WIYW RXYRLRXCGS GSGP* | 19 |
| BD42 0.01% | WLYY KXWRLHXCGS GSGA* | 19 |
| BD42 0.01% | WIYY QXWRIKXCGS GSGA* | 19 |
| BD32 0.02% | WLYY KXWRLHXCGS GS* | 17 |
| BD42 0.01% | WLYY KXWRLHXCGS GS* | 17 |
| BD42 0.01% | WLYY KXWRLHXYGS GSGS* | 19 |
| BD42 0.01% | WLYY KXWRLHXWGS GSGS* | 19 |
| BD42 0.01% | WLYY KXWRLHXCGS WSGS* | 19 |
| BD42 0.04% | WIYW RXYRLRXCGS GS* | 17 |
| BD42 0.01% | WIYW RXYRLRXCGP GSGS* | 19 |
| BD42 0.01% | WIYW RXYRLRXCGS GLGLR* | 20 |
| BD42 0.01% | WIYW RXYRLRXCGF GSGS* | 19 |
| BD42 0.01% | WIYY QXWRIKXCGS GPGS* | 19 |
| BD42 0.01% | WIYY QXWRIKXGGS GSGS* | 19 |
| BD42 0.01% | WXYW RXYVLKICGS GS* | 17 |
| BD42 0.01% | WXYW RXYVLKICGS GSWS* | 19 |
| BD42 0.01% | WXYW RXYVLKICGF GSGS* | 19 |
| BD42 2.47% | WXYW RXYVLKIC* | 13 |
| BD42 0.01% | WXYW RXYVLKICGL GLGLR* | 20 |
| BD42 0.01% | WXVFYHGGX NYSILRICGS GSGA* | 24 |
| BD42 0.01% | WXVFYHGGX NYSILRICGS GSWS* | 24 |
| BD42 0.01% | WXVFYHGGX NYSILRICGL GLGLR* | 25 |
| BD42 1.53% | WXVFYHGGX NYSILRIC* | 18 |
| BD42 0.01% | WSGW WXVPXKICGL GLGLR* | 20 |
| BD32 0.11% | WXFY QXWRLQIC* | 13 |
| BD42 0.48% | WXFY QXWRLQIC* | 13 |
| BD42 0.01% | WXFY QXWRLQICGS WSGS* | 19 |
| BD42 0.01% | WXVWR KXLALQICGS GS* | 18 |
| BD42 0.01% | WXVWR KXLALQICGS GYGS* | 20 |
| BD42 0.01% | WXVWR KXLALQICGS GPGS* | 20 |
| BD42 0.01% | WXVWR KXLALQICGP GSGS* | 20 |
| BD42 0.01% | WXVWR KXLALQICGS GLGLR* | 21 |
| BD42 2.12% | WXVWR KXLALQIC* | 14 |
| BD42 0.87% | WXVY GXLALKIC* | 13 |
| BD42 0.01% | WXVY GXLALKICGS GSGP* | 19 |
| BD42 0.01% | WXVWR KXLALQI*GS GSGS* | 20 |
| BD42 0.01% | WSGW WXVPXKICGS GS* | 17 |
| BD42 0.01% | WSGW WXVPXKICGS GSGP* | 19 |
| BD42 0.01% | WSGW WXVPXKICGS GYGS* | 19 |
| BD42 0.01% | WSGW WXVPXKICGS GTGP* | 19 |
| BD42 0.01% | WSGW WXVPXKICGS GLGLR* | 20 |
| BD42 0.01% | WLYY KXWRLHXCGA GAGS* | 19 |
| BD22 0.88% | WNGW WXVPXKIC* | 13 |
| BD42 4.72% | WSGW WXVPXKIC* | 13 |
| BD42 0.01% | WRGW WXVPXKIC* | 13 |
| BD42 0.01% | WSGW WXAPXKIC* | 13 |
| BD42 0.01% | WSGW WXEPXKIC* | 13 |
| BD32 1.96% | WIYY RXWRLRXC* | 13 |
| BD32 1.45% | WLYY RXWRLRXC* | 13 |
| BD32 1.08% | WIYY RXWRLFXC* | 13 |
| BD32 1.03% | WIYY KXWRLKXC* | 13 |
| BD32 0.96% | WIYY RXWRLWXC* | 13 |
| BD32 0.70% | WLYY RXWRLNXC* | 13 |
| BD32 0.70% | WLYY RXWRLKXC* | 13 |
| BD32 0.64% | WIYY RXWRLKXC* | 13 |
| BD32 0.59% | WIYY KXWRLLXC* | 13 |
| BD32 0.56% | WIYF RXWRLHXC* | 13 |
| BD32 0.56% | WIYY QXWRLNXC* | 13 |
| BD32 0.31% | WLYY RXWRLFXC* | 13 |
| BD32 0.26% | WIYY RXWRLYXC* | 13 |
| BD32 0.16% | WIYW KXYRLRXC* | 13 |
| BD32 0.14% | WIYY AXWRLHXC* | 13 |
| BD32 0.12% | WIYY AXWRLRXC* | 13 |
| BD32 0.09% | WIYY LXWRLKXC* | 13 |
| BD32 0.09% | WIYF RXWRLRXC* | 13 |
| BD32 0.08% | WIYY KXWRLHXC* | 13 |
| BD32 0.07% | WXYY RXWRLRXC* | 13 |
| BD32 0.05% | WXYY RXWRLXIC* | 13 |
| BD32 0.05% | WIYW NXYRLRXC* | 13 |
| BD32 0.05% | WIYF NXWRLKXC* | 13 |
| BD32 0.05% | WTYY RXWRLRXC* | 13 |
| BD32 0.04% | WAYY RXWRLKXC* | 13 |
| BD32 0.03% | WIYY GXWRLLXC* | 13 |
| BD32 0.03% | WAYY RXWRLLXC* | 13 |
| BD32 0.03% | WQYY RXWRLLXC* | 13 |
| BD32 0.03% | WTYY RXWRLYXC* | 13 |
| BD32 0.02% | WLYY RXYRLRXC* | 13 |
| BD32 0.02% | WXYY RXWRLHXC* | 13 |
| BD42 4.48% | WIYW RXYRLRXC* | 13 |
| BD42 0.01% | WIYW RLYRLRXC* | 13 |
| BD42 0.01% | WIYC RXYRLRXC* | 13 |
| BD42 0.01% | WIYW RXYRLRLC* | 13 |
| BD42 0.01% | WIYW RXYRLRC* | 12 |
| BD32 1.26% | WLYY KXWRLHXC* | 13 |

| | | |
|---|---|---|
| BD32 1.15% | WLYY QXWRLWXC* | 13 |
| BD32 0.97% | WLYY QXWRLNXC* | 13 |
| BD32 0.59% | WLYY KXWRLQXC* | 13 |
| BD32 0.42% | WLYF QXWRLYXC* | 13 |
| BD32 0.41% | WLYY KXWRLYXC* | 13 |
| BD32 0.25% | WLYY HXWRLYXC* | 13 |
| BD32 0.24% | WLYY KXWRLKXC* | 13 |
| BD32 0.24% | WLYF RXWRLYXC* | 13 |
| BD32 0.21% | WLYF QXWRLRXC* | 13 |
| BD32 0.21% | WLYF KXWRLYXC* | 13 |
| BD32 0.21% | WLYF RXWRLNXC* | 13 |
| BD22 0.06% | WLYHPYN XAXRLLXC* | 16 |
| BD32 0.26% | WLYW QXYRLLXC* | 13 |
| BD32 0.20% | WLYY QXWRLLXC* | 13 |
| BD32 0.17% | WLYW KXYRLLXC* | 13 |
| BD32 0.16% | WLYY NXWRLLXC* | 13 |
| BD32 0.06% | WLYY XXYRLLXC* | 13 |
| BD32 0.03% | WLYW QXFRLLXC* | 13 |
| BD32 0.03% | WLYW HXYRLLXC* | 13 |
| BD22 1.29% | WHYWL LRXGKIHKLR C* | 17 |
| BD42 0.01% | WXNWCWL LRXLLLRC* | 16 |
| BD42 0.01% | WKYWL YITXSFRLLL RC* | 18 |
| BD42 0.01% | WXNWYWL XRXLLLRC* | 16 |
| BD32 0.16% | WLYY LXWRLYXC* | 13 |
| BD32 0.15% | WLYF XXWRLKXC* | 13 |
| BD32 0.13% | WLYY HXWRLFXC* | 13 |
| BD32 0.13% | WLYF KXWRLRXC* | 13 |
| BD32 0.07% | WLYF KXWRLXXC* | 13 |
| BD32 0.07% | WLYY QXYRLRXC* | 13 |
| BD32 0.07% | WLYY HXWRLKXC* | 13 |
| BD32 0.06% | WLYF QXYRLRXC* | 13 |
| BD32 0.06% | WLYW QXWRLRXC* | 13 |
| BD32 0.06% | WLYY NXWRLXXC* | 13 |
| BD32 0.05% | WLYY XXYRLRXC* | 13 |
| BD32 0.04% | WLYY KXYRLRXC* | 13 |
| BD32 0.03% | WLYF QXWRLQXC* | 13 |
| BD32 0.03% | WLYH RXWRLFXC* | 13 |
| BD32 0.02% | WLYY AXWRLWXC* | 13 |
| BD32 0.02% | WLYW KXYRLRXC* | 13 |
| BD32 0.02% | WLYY LXWRLWXC* | 13 |
| BD42 1.61% | WLYY KXWRLHXC* | 13 |
| BD22 0.04% | WYWLYK ESXWLHWRHI C* | 18 |
| BD42 0.01% | WLYY KXWRLHIC* | 13 |
| BD42 0.01% | WLYY NXWRLQXC* | 13 |
| BD32 1.11% | WLYW RXYRLHXC* | 13 |
| BD32 0.31% | WLYY QXYRLHXC* | 13 |
| BD32 0.20% | WLYY RXYRLHXC* | 13 |
| BD32 0.16% | WLYW KXFRLHXC* | 13 |
| BD32 0.13% | WLYY QXFRLHXC* | 13 |
| BD32 0.09% | WLYW QXFRLHXC* | 13 |
| BD32 0.06% | WLYW KXYRLHXC* | 13 |
| BD32 0.05% | WLYY LXYRLHXC* | 13 |
| BD32 0.04% | WXYY KXWRLLIC* | 13 |
| BD32 0.04% | WXYY KXWRLRXC* | 13 |
| BD32 0.04% | WLYY KXYRLHXC* | 13 |
| BD32 0.03% | WQYY KXWRLKXC* | 13 |
| BD32 0.03% | WLYW HXFRLHXC* | 13 |
| BD42 0.01% | WPYY KXWRLHXC* | 13 |
| BD32 0.16% | WXYY QXWRLKVC* | 13 |
| BD32 0.15% | WXYY QXWRLLXC* | 13 |
| BD32 0.07% | WVYY QXWRLRXC* | 13 |
| BD32 0.06% | WXYY QXWRLRXC* | 13 |
| BD32 0.04% | WLYW KXFRLWXC* | 13 |
| BD32 0.03% | WLYW RXYRLWXC* | 13 |
| BD32 0.77% | WLYW KXYRLKXC* | 13 |
| BD32 0.34% | WLYW QXYRLNXC* | 13 |
| BD32 0.16% | WIYW RXYRLLXC* | 13 |
| BD32 0.13% | WLYY KXYRLKXC* | 13 |
| BD32 0.08% | WLYW HXYRLKXC* | 13 |
| BD32 0.07% | WLYW QXYRLQXC* | 13 |
| BD32 0.06% | WLYW GXYRLQXC* | 13 |
| BD32 0.06% | WLYY QXYRLKXC* | 13 |
| BD32 0.05% | WLYY XXYRLYXC* | 13 |
| BD32 0.05% | WLYY RXYRLXXC* | 13 |
| BD32 0.04% | WLYY RXYRLFXC* | 13 |
| BD32 0.03% | WLYW XXYRLKXC* | 13 |
| BD32 0.03% | WLYY QXYRLXXC* | 13 |
| BD32 0.03% | WLYF QXYRLXXC* | 13 |
| BD32 0.03% | WLYY QXYRLQXC* | 13 |
| BD32 0.02% | WLYW GXYRLYXC* | 13 |
| BD32 0.02% | WLYW AXYRLYXC* | 13 |
| BD32 0.02% | WLYY KXYRLFXC* | 13 |
| BD32 0.02% | WLYW KXYRLFXC* | 13 |
| BD42 0.01% | WIYW RXYRLWXC* | 13 |
| BD42 0.01% | W THYRVEL*VG DISXXNWCWL XRXLLLRC* | 30 |
| BD42 0.01% | WVEL*VG DISXXNWCWL XRXLLLRC* | 26 |
| BD42 0.01% | W THYRVEL*VG DISXIYWRXY RLRXC* | 27 |
| BD42 0.34% | WIYSLYL TXKIRIQIKC * | 18 |
| BD42 0.01% | W THYRVEL*VG DISXIYSLYL TXKIRIQIKC * | 32 |
| BD32 0.30% | WIYW KXYRLKXC* | 13 |
| BD32 0.06% | WIYW NXYRLKXC* | 13 |
| BD32 0.02% | WIYW QXYRLKXC* | 13 |
| BD32 0.03% | WIYW NXYRLWXC* | 13 |

| | | |
|---|---|---|
| BD32 0.80% | WIYW QXYRLNXC* | 13 |
| BD32 0.64% | WIYW QXYRLYXC* | 13 |
| BD32 0.38% | WIYW QXYRLHXC* | 13 |
| BD32 0.20% | WIYW KXYRLLXC* | 13 |
| BD32 0.16% | WIYW KXYRLXXC* | 13 |
| BD32 0.13% | WQYW QXYRLWXC* | 13 |
| BD32 0.08% | WVYW QXYRLHXC* | 13 |
| BD32 0.06% | WIYY RXYRLYXC* | 13 |
| BD32 0.06% | WIYY QXYRLLXC* | 13 |
| BD32 0.04% | WIYF QXYRLHXC* | 13 |
| BD32 0.04% | WIYW HXYRLLXC* | 13 |
| BD32 0.03% | WAYW QXYRLHXC* | 13 |
| BD32 0.03% | WVYW QXYRLXXC* | 13 |
| BD32 0.03% | WIYY QXYRLXXC* | 13 |
| BD42 0.01% | WIYW QXYRLYXC* | 13 |
| BD32 0.23% | WLYW RXFRLKXC* | 13 |
| BD32 0.20% | WXYW KXXRLKIC* | 13 |
| BD32 0.08% | WLYW QXFRLKXC* | 13 |
| BD32 0.04% | WLYY XXFRLKXC* | 13 |
| BD32 0.03% | WXYW KXYRLNVC* | 13 |
| BD32 0.05% | WQYF RXWRLHXC* | 13 |
| BD32 0.03% | WXYF RXWRLKIC* | 13 |
| BD32 0.06% | WLYY QXFRLYXC* | 13 |
| BD32 0.06% | WLYW KXFRLYXC* | 13 |
| BD32 0.04% | WLYY KXFRLYXC* | 13 |
| BD32 0.13% | WLYY QXFRLFXC* | 13 |
| BD32 0.03% | WLYF QXFRLFXC* | 13 |
| BD32 0.13% | WXYY HXWRLKIC* | 13 |
| BD32 0.04% | WLYW KXFRLXXC* | 13 |
| BD32 0.06% | WXYW KXFRLQVC* | 13 |
| BD32 0.15% | WLYY QXWRIXXC* | 13 |
| BD32 0.14% | WLYY KXWRILXC* | 13 |
| BD32 0.13% | WLYY XXWRIRXC* | 13 |
| BD32 0.06% | WLYY RXWRIQXC* | 13 |
| BD32 0.04% | WLYY KXWRIHXC* | 13 |
| BD32 0.04% | WLYY RXWRIFXC* | 13 |
| BD32 0.03% | WLYY *XWRIKXC* | 13 |
| BD32 0.03% | WLYW QXWRIKXC* | 13 |
| BD42 0.45% | WIYY QXWRIKXC* | 13 |
| BD32 0.02% | WIYY QXFRLQXC* | 13 |
| BD42 0.04% | WAYH TIRLXWRXPS ISCGSGS* | 22 |
| BD42 0.01% | WAYH TIRLXWRXPS ISCGSGSGP* | 24 |
| BD42 0.01% | WAYH TIRLXWRXPS ISCGSGSES* | 24 |
| BD42 0.01% | WAYH TIRLXWRXPS ISCGSGSGA* | 24 |
| BD42 0.01% | WAYH TIRLXWRXPS ISCGSGSWS* | 24 |
| BD42 0.01% | WAYH TIRLXWRXPS ISCGSGSGT* | 24 |
| BD42 0.01% | WAYH TIRLXWRXPS ISCGFGSGS* | 24 |
| BD42 0.01% | WAYH TIRLXWRXPS ISCGSGSGLR * | 25 |
| BD42 0.02% | WAYH TIRLXWRXPS ISCGSGLGLR * | 25 |
| BD42 0.02% | WAYH TIRLXWRXPS ISCGLGLGLR * | 25 |
| BD42 0.01% | WAYH TIRLXWRXPS ISCGSGPGS* | 24 |
| BD42 0.01% | WAYH TIRLXWRXPS IRCGSGSGA* | 24 |
| BD42 0.01% | WAYH TIRLXWRXPS ISCGSGTGS* | 24 |
| BD42 0.01% | WAYH TIRLXWRXPS ISCGSGAGS* | 24 |
| BD42 0.01% | WAYH TIRLXWRXPS ISWGSGSGS* | 24 |
| BD42 0.01% | WAYH TIRLXWRXPS ISFGSGSGS* | 24 |
| BD42 9.47% | WAYH TIRLXWRXPS ISC* | 18 |
| BD42 0.03% | WAYH TIRLXRRXPS ISC* | 18 |
| BD42 0.02% | WAYH TIRLXWRXPS IRC* | 18 |
| BD42 0.01% | WAYH TIRLXWRXPP ISC* | 18 |
| BD42 0.01% | WAYH TIRLXGRXPS ISC* | 18 |
| BD42 0.01% | WAYH TIRLXWRXPA ISC* | 18 |
| BD42 0.01% | WAYH TIRLXCRXPS ISC* | 18 |
| BD42 0.01% | WAYH TIRLLWRXPS ISC* | 18 |
| BD42 0.01% | WAYH TIRLXWRXPT ISC* | 18 |
| BD42 0.01% | WAYH TISLXWRXPS ISC* | 18 |
| BD42 0.01% | WAYH TIRLXWRLPS ISC* | 18 |
| BD42 0.01% | WSYH TIRLXWRXPS ISC* | 18 |
| BD42 0.01% | WAYH AIRLXWRXPS ISC* | 18 |
| BD42 0.01% | WAYH TIRLXWRIPS ISC* | 18 |
| BD42 0.01% | WAYH TIRLTWRXPS ISC* | 18 |
| BD42 0.01% | WVEL *VGDISXAYH TIRLXWRXPS ISC* | 28 |
| BD42 0.01% | WAYH TIRLKWRXPS ISC* | 18 |
| BD42 1.55% | WAYS WLCSXRLLLK KXC* | 18 |
| BD42 0.01% | WAYS WLCSXRLLLK KXCGSGSGA* | 24 |
| BD42 0.01% | WLLFYNHX LIRRLLXCGS GS* | 21 |
| BD42 0.01% | WLLFYNHX LIRRLLXRGS GSGS* | 23 |
| BD42 0.01% | WLLFYNHX LIRRLLXCGS GSGT* | 23 |
| BD42 1.53% | WLLFYNHX LIRRLLXC* | 17 |
| BD42 0.01% | WPLFYNHX LIRRLLXC* | 17 |
| BD32 0.34% | WLFY RXWRLLXC* | 13 |
| BD32 0.19% | WLFY XXWRLYXC* | 13 |
| BD32 0.10% | WLFW KXYRLLXC* | 13 |
| BD32 0.08% | WLFY QXWRLNXC* | 13 |
| BD32 0.06% | WLFY KXWRLLXC* | 13 |
| BD32 0.04% | WLFY QXWRLQXC* | 13 |
| BD32 0.02% | WLFY QXYRLLXC* | 13 |
| BD32 0.02% | WLFF RXWRLXXC* | 13 |
| BD42 0.27% | WLFY XXWRLYXC* | 13 |
| BD32 0.16% | WLFW RXYRLYXC* | 13 |
| BD32 0.03% | WLFY QXYRLYXC* | 13 |
| BD32 0.13% | WLFWQXY RLKXC* | 13 |
| BD32 0.06% | WLFWKXY RLQXC* | 13 |

```
BD32 0.05% ---------- ---------- ---------- ------WLFY RXYRLHXC*- ---------- ---------- ---------- 13
BD32 0.05% ---------- ---------- ---------- ---WXFWRXY RLQIC*---- ---------- ---------- ---------- 13
BD32 0.04% ---------- ---------- ---------- ------WLFY QXYRLRXC*- ---------- ---------- ---------- 13
BD32 0.03% ---------- ---------- ---------- ------WLFW QXYRLXXC*- ---------- ---------- ---------- 13
BD32 0.04% ---------- ---------- ---------- ------WIFY QXWRLKXC*- ---------- ---------- ---------- 13
BD32 0.10% ---------- ---------- ---------- ------WXYW QXFRLKIC*- ---------- ---------- ---------- 13
BD32 0.09% ---------- ---------- ---------- ------WIYW QXFRLQXC*- ---------- ---------- ---------- 13
BD32 0.09% ---------- ---------- ---------- ------WXRW RXYRLQIC*- ---------- ---------- ---------- 13
BD32 0.03% ---------- ---------- ---------- ------WLRW RXYRLWXC*- ---------- ---------- ---------- 13
BD32 0.07% ---------- ---------- ---------- ------WIFY KXWRLLXC*- ---------- ---------- ---------- 13
BD32 0.03% ---------- ---------- ---------- ------WQFY KXWRLLXC*- ---------- ---------- ---------- 13
BD32 0.03% ---------- ---------- ---------- ------WIFY HXWRLKXC*- ---------- ---------- ---------- 13
BD42 0.03% ---------- ---------- ---------- ------WYDV FLTRXYGXXK VACGSGS*-- ---------- ---------- 22
BD42 0.01% ---------- ---------- ---------- ------WYDV FLTRXYGXXK VACGSGLGLR *--------- ---------- 25
BD42 0.01% ---------- ---------- ---------- ------WYDV FLTRXYGXXK VACGSGSWS* ---------- ---------- 24
BD42 0.01% ---------- ---------- ---------- ------WYDV FLTRXYGXXK VACGSGSGP* ---------- ---------- 24
BD42 0.01% ---------- ---------- ---------- ------WYDV FLTRXYGXXK VACGAGSGS* ---------- ---------- 24
BD42 0.01% ---------- ---------- ---------- ------WYDV FLTRXYGXXK VACGLGLGLR *--------- ---------- 25
BD42 3.84% ---------- ---------- ---------- ------WYDV FLTRXYGXXK VAC*------ ---------- ---------- 18
BD42 0.01% ---------- ---------- ---------- ------WYDE FLTRXYGXXK VAC*------ ---------- ---------- 18
BD42 0.01% ---------- ---------- ---------- ------WYDV FLTRXHGXXK VAC*------ ---------- ---------- 18
BD42 0.01% ---------- ---------- ---------- ------WYDV LLTRXYGXXK VAC*------ ---------- ---------- 18
BD42 0.01% ---------- ---------- ---------- ------WYDV FLTRXYGXXE VAC*------ ---------- ---------- 18
BD42 0.01% ---------- ---------- ---------- ------WYDV FLTHXYGXXK VAC*------ ---------- ---------- 18
BD42 0.01% ---------- ---------- ---------- ------WHDV FLTRXYGXXK VAC*------ ---------- ---------- 18
BD42 0.01% ---------- ---------- ---------- ------WYDV FLTRXYGXXX VAC*------ ---------- ---------- 18
BD42 0.01% ---------- ---------- ---------- ------WYDV F*RVCXV**R LRVGLGLGLR *--------- ---------- 25
BD42 0.01% ---------- ---------- ---------- -------WXC F*RVCXV**R LRVGLGLGLR *--------- ---------- 24
BD42 0.03% ---------- ---------- ---------- ------WIWN RSSIXKLCLL VRVGLGLGLR *--------- ---------- 25
BD42 0.01% ---------- ---------- ---------- ---------W XHRPCXPRFC FRVGLGLGLR *--------- ---------- 22
BD42 1.16% ---------- ---------- ---------- ------WYSY FLSRXXRTXV LAC*------ ---------- ---------- 18
BD42 0.01% ---------- ---------- ---------- ------WYSY FLSRXXRTXA LAC*------ ---------- ---------- 18
BD42 0.01% ---------- ---------- ---------- ------WYSY FLSRXXRTXV LACGLGLGLR *--------- ---------- 25
BD42 0.01% ---------- ---------- ---------- ------WYSY FCRV*CGRWC WLVGLGLGLR *--------- ---------- 25
BD42 0.05% ---------- ---------- ---------- ---WXNWCWL XRXLLLRCGS GSWS*----- ---------- ---------- 22
BD42 0.01% ---------- ---------- ---------- ---WXNWCWL XRXLLLRCGS GSWVLGR*-- ---------- ---------- 25
BD42 0.17% ---------- ---------- ---------- ---WXNWCWL XRXLLLRCGS GS*------- ---------- ---------- 20
BD42 0.03% ---------- ---------- ---------- ---WXNWCWL XRXLLLRCGS GAGS*----- ---------- ---------- 22
BD42 0.01% ---------- ---------- ---------- ---WXNWCWL XRXLLLRWGS GAGS*----- ---------- ---------- 22
BD42 0.03% ---------- ---------- ---------- ---WXNWCWL XRXLLLRCGP GSGS*----- ---------- ---------- 22
BD42 0.01% ---------- ---------- ---------- ---WXNWCWL XRXLLLRCGP GSGP*----- ---------- ---------- 22
BD42 0.01% ---------- ---------- ---------- ---WXNWCWL XRXLLLRCGP GSGT*----- ---------- ---------- 22
BD42 0.01% ---------- ---------- ---------- ---WXHWCWL XRXLLLRCGP GSGS*----- ---------- ---------- 22
BD42 0.03% ---------- ---------- ---------- ---WXNWCWL XRXLLLRGGS GSGS*----- ---------- ---------- 22
BD42 0.02% ---------- ---------- ---------- ---WXNWCWL XRXLLLRRGS GSGS*----- ---------- ---------- 22
BD42 0.01% ---------- ---------- ---------- ---WXNWCWL XRXLLLRSGS GSGS*----- ---------- ---------- 22
BD42 0.01% ---------- ---------- ---------- ---WXNWCWL XRXLLLRWGS GSGS*----- ---------- ---------- 22
BD42 0.01% ---------- ---------- ---------- ---WXNWCWL XRXLLLR*GS GSGS*----- ---------- ---------- 22
BD42 0.01% ---------- ---------- ---------- ---WXNWCWL XRXLLLRYGS GSGS*----- ---------- ---------- 22
BD42 0.01% ---------- ---------- ---------- ---WXNWCWL XRXLLLRGGS GSGA*----- ---------- ---------- 22
BD42 0.02% ---------- ---------- ---------- ---WXNWCWL XRXLLLRCGS GSGP*----- ---------- ---------- 22
BD42 0.02% ---------- ---------- ---------- ---WXNWCWL XRXLLLRCGS GSGY*----- ---------- ---------- 22
BD42 0.01% ---------- ---------- ---------- ---WXNWCWL XRXLLLRCGS GSGLR*---- ---------- ---------- 23
BD42 0.01% ---------- ---------- ---------- ---WXNWCWL XRXLLLRCGS GSGT*----- ---------- ---------- 22
BD42 0.01% ---------- ---------- ---------- ---WXNWCWL XRXLLLRCGS GSGF*----- ---------- ---------- 22
BD42 0.02% ---------- ---------- ---------- ---WXNWCWL XRXLLLRCGS GLGLR*---- ---------- ---------- 23
BD42 0.02% ---------- ---------- ---------- ---WXNWCWL XRXLLLRCGS GYGS*----- ---------- ---------- 22
BD42 0.01% ---------- ---------- ---------- ---WXNWCWL XRXLLLRCGS ESGS*----- ---------- ---------- 22
BD42 0.01% ---------- ---------- ---------- ---WXNWCWL XRXLLLRCGS RSGS*----- ---------- ---------- 22
BD42 0.01% ---------- ---------- ---------- ---WXNWCWL XRXLLLRCGS WSGS*----- ---------- ---------- 22
BD42 0.01% ---------- ---------- ---------- ---WXNWCWL XRXLLLRCGS ASGS*----- ---------- ---------- 22
BD42 0.01% ---------- ---------- ---------- ---WXHWCWL XRXLLLRCGS VSGS*----- ---------- ---------- 22
BD42 0.01% ---------- ---------- ---------- ---WXNWCWL XRXLLLRCGY GSGS*----- ---------- ---------- 22
BD42 0.01% ---------- ---------- ---------- ---WXNWCWL XRXLLLRCGS GSRS*----- ---------- ---------- 22
BD42 0.01% ---------- ---------- ---------- ---WXNWCWL XRXLLLRCGS GTGS*----- ---------- ---------- 22
BD42 0.01% ---------- ---------- ---------- ---WXNWCWL XRXLLLRCGS *--------- ---------- ---------- 18
BD42 0.01% ---------- ---------- ---------- ---WXNWCWL XRXLLLRCGS GSVS*----- ---------- ---------- 22
BD42 0.01% ---------- ---------- ---------- ---WXNWCWL XRXLLLRCRS GSGS*----- ---------- ---------- 22
BD42 0.01% ---------- ---------- ---------- ---WXNWCWL XRXLLLRCGT GSGS*----- ---------- ---------- 22
BD42 0.01% ---------- ---------- ---------- ---WXNWCWL XRXLLLRCGF GSGS*----- ---------- ---------- 22
BD42 0.01% ---------- ---------- ---------- ---WXNWCWL XRXLLLRCGS GPGS*----- ---------- ---------- 22
BD42 0.01% ---------- ---------- ---------- ---WXNWCWL XRXLLLRCGA GSGS*----- ---------- ---------- 22
BD42 0.01% ---------- ---------- ---------- ---WXNWCWL XRXLLLRCAS GSGS*----- ---------- ---------- 22
BD42 0.02% ---------- ---------- ---------- --WCNRWCWL XXXLLLLNCG SGS*------ ---------- ---------- 22
BD42 0.01% ---------- ---------- ---------- --WCNRWCWL XXXLLLLNCG SGYGS*---- ---------- ---------- 24
BD42 0.01% ---------- ---------- ---------- --WCNRWCWL XXXLLLLNCG SGSWS*---- ---------- ---------- 24
BD42 2.33% ---------- ---------- ---------- --WCNRWCWL XXXLLLLNC* ---------- ---------- ---------- 18
BD42 0.03% ---------- ---------- ---------- --WCNRRCWL XXXLLLLNC* ---------- ---------- ---------- 18
BD42 0.01% ---------- ---------- ---------- --WCNRGCWL XXXLLLLNC* ---------- ---------- ---------- 18
BD42 0.01% ---------- ---------- ---------- --WCNRWCWL XXXLLLLNCG LGLGLR*--- ---------- ---------- 25
BD42 0.01% ---------- ---------- ---------- ---WXSWCCL XRXLLLRRGS GSGS*----- ---------- ---------- 22
BD42 0.03% ---------- ---------- ---------- ---WXNWCWL XRXLLLRCGL GLGLR*---- ---------- ---------- 23
BD42 0.01% ---------- ---------- ---------- ---WWKXHRC XLITIRRCGS GS*------- ---------- ---------- 20
BD42 0.01% ---------- ---------- ---------- ---WWKXHRC XLITIRRCGS GLGLR*---- ---------- ---------- 23
BD42 0.78% ---------- ---------- ---------- ---WWKXHRC XLITIRRC*- ---------- ---------- ---------- 16
BD42 0.01% ---------- ---------- ---------- ---WWKXHRS XLITIRRC*- ---------- ---------- ---------- 16
BD42 0.02% ---------- ---------- ---------- ---WXGYLCL RXRIQRTYNC GSGS*----- ---------- ---------- 22
BD42 0.01% ---------- ---------- ---------- ---WXGYLCL RXRIQRTYNC GSGSGP*--- ---------- ---------- 24
BD42 0.01% ---------- ---------- ---------- ---WXGYLCL RXRIQRTYNC GPGSGS*--- ---------- ---------- 24
BD42 0.01% ---------- ---------- ---------- ---WXGYLCL RXRIQRTYNC GSGSGY*--- ---------- ---------- 24
BD42 0.01% ---------- ---------- ---------- ---WXGYLCL RXRIQRTYNC GSGSES*--- ---------- ---------- 24
BD42 0.01% ---------- ---------- ---------- ---WXGYLCL RXRIQRTYNC GSGSWS*--- ---------- ---------- 24
BD42 0.01% ---------- ---------- ---------- ---WXGYLCL RXRIQRTYNC GSGSGT*--- ---------- ---------- 24
```

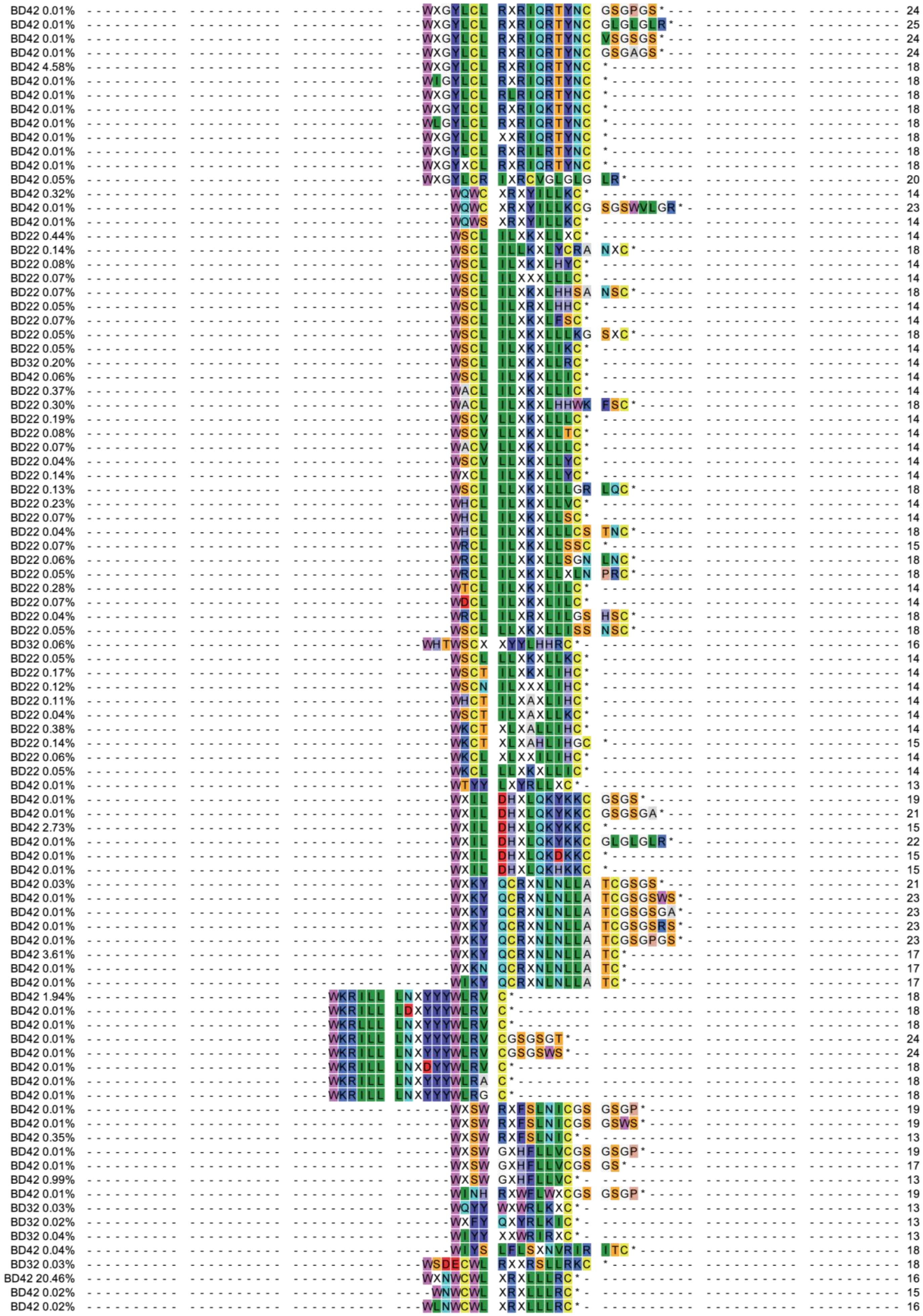
BD42 0.01% WXGYLCL RXRIQRTYNC GSGPGS* 24
BD42 0.01% WXGYLCL RXRIQRTYNC GLGLGLR* 25
BD42 0.01% WXGYLCL RXRIQRTYNC VSGSGS* 24
BD42 0.01% WXGYLCL RXRIQRTYNC GSGAGS* 24
BD42 4.58% WXGYLCL RXRIQRTYNC * 18
BD42 0.01% WIGYLCL RXRIQRTYNC * 18
BD42 0.01% WXGYLCL RLRIQRTYNC * 18
BD42 0.01% WXGYLCL RXRIQKTYNC * 18
BD42 0.01% WLGYLCL RXRIQRTYNC * 18
BD42 0.01% WXGYLCL XXRIQRTYNC * 18
BD42 0.01% WXGYLCL RXRILRTYNC * 18
BD42 0.01% WXGYXCL RXRIQRTYNC * 18
BD42 0.05% WXGYLCR IXRCVGLGLG LR* 20
BD42 0.32% WQWC XRXYILLKC* 14
BD42 0.01% WQWC XRXYILLKCG SGSWVLGR* 23
BD42 0.01% WQWS XRXYILLKC* 14
BD22 0.44% WSCL ILXKXLLXC* 14
BD22 0.14% WSCL ILLKXLYCRA NXC* 18
BD22 0.08% WSCL ILXKXLHYC* 14
BD22 0.07% WSCL ILXXXLLLC* 14
BD22 0.07% WSCL ILXKXLHHSA NSC* 18
BD22 0.05% WSCL ILXRXLHHC* 14
BD22 0.07% WSCL ILXKXLFSC* 14
BD22 0.05% WSCL ILXKXLLLKG SXC* 18
BD22 0.05% WSCL ILXKXLIKC* 14
BD32 0.20% WSCL ILXKXLLRC* 14
BD42 0.06% WSCL ILXKXLLIC* 14
BD22 0.37% WACL ILXKXLLIC* 14
BD22 0.30% WACL ILXKXLHHWK FSC* 18
BD22 0.19% WSCV LLXKXLLLC* 14
BD22 0.08% WSCV LLXKXLLTC* 14
BD22 0.07% WACV LLXKXLLLC* 14
BD22 0.04% WSCV LLXKXLLYC* 14
BD22 0.14% WXCL ILXKXLLYC* 14
BD22 0.13% WSCI LLXKXLLLGR LQC* 18
BD22 0.23% WHCL ILXKXLLVC* 14
BD22 0.07% WHCL ILXKXLLSC* 14
BD22 0.04% WHCL ILXKXLLLCS TNC* 18
BD22 0.07% WRCL ILXKXLLSSC * 15
BD22 0.06% WRCL ILXKXLLSGN LNC* 18
BD22 0.05% WRCL ILXKXLLXLN PRC* 18
BD22 0.28% WTCL ILXKXLILC* 14
BD22 0.07% WDCL ILXKXLILC* 14
BD22 0.04% WRCL ILXRXLILGS HSC* 18
BD22 0.05% WSCL LLXKXLLISS NSC* 18
BD32 0.06% WHTWSCX XYYLHHRC* 16
BD22 0.05% WSCL LLXKXLLKC* 14
BD22 0.17% WSCT ILXKXLIHC* 14
BD22 0.12% WSCN ILXXXLIHC* 14
BD22 0.11% WHCT ILXAXLIHC* 14
BD22 0.04% WSCT ILXAXLLKC* 14
BD22 0.38% WKCT XLXALLIHC* 14
BD22 0.14% WKCT XLXAHLIHGC * 15
BD22 0.06% WKCL XLXXILIHC* 14
BD22 0.05% WKCL LLXKXLLIC* 14
BD42 0.01% WTYY LXYRLLXC* 13
BD42 0.01% WXIL DHXLQKYKKC GSGS* 19
BD42 0.01% WXIL DHXLQKYKKC GSGSGA* 21
BD42 2.73% WXIL DHXLQKYKKC * 15
BD42 0.01% WXIL DHXLQKYKKC GLGLGLR* 22
BD42 0.01% WXIL DHXLQKDKKC * 15
BD42 0.01% WXIL DHXLQKHKKC * 15
BD42 0.03% WXKY QCRXNLNLLA TCGSGS* 21
BD42 0.01% WXKY QCRXNLNLLA TCGSGSWS* 23
BD42 0.01% WXKY QCRXNLNLLA TCGSGSGA* 23
BD42 0.01% WXKY QCRXNLNLLA TCGSGSRS* 23
BD42 0.01% WXKY QCRXNLNLLA TCGSGPGS* 23
BD42 3.61% WXKY QCRXNLNLLA TC* 17
BD42 0.01% WXKN QCRXNLNLLA TC* 17
BD42 0.01% WIKY QCRXNLNLLA TC* 17
BD42 1.94% WKRILL LNXYYYWLRV C* 18
BD42 0.01% WKRILL LDXYYYWLRV C* 18
BD42 0.01% WKRLLL LNXYYYWLRV C* 18
BD42 0.01% WKRILL LNXYYYWLRV CGSGSGT* 24
BD42 0.01% WKRILL LNXYYYWLRV CGSGSWS* 24
BD42 0.01% WKRILL LNXDYYWLRV C* 18
BD42 0.01% WKRILL LNXYYYWLRA C* 18
BD42 0.01% WKRILL LNXYYYWLRG C* 18
BD42 0.01% WXSW RXFSLNICGS GSGP* 19
BD42 0.01% WXSW RXFSLNICGS GSWS* 19
BD42 0.35% WXSW RXFSLNIC* 13
BD42 0.01% WXSW GXHFLLVCGS GSGP* 19
BD42 0.01% WXSW GXHFLLVCGS GS* 17
BD42 0.99% WXSW GXHFLLVC* 13
BD42 0.01% WINH RXWFLWXCGS GSGP* 19
BD32 0.03% WQYY WXWRLKXC* 13
BD32 0.02% WXFY QXYRLKIC* 13
BD32 0.04% WIYY XXWRIRXC* 13
BD42 0.04% WIYS LFLSXNVRIR ITC* 18
BD32 0.03% WSDECWL RXXRSLLRKC * 18
BD42 20.46% WXNWCWL XRXLLLRC* 16
BD42 0.02% WNWCWL XRXLLLRC* 15
BD42 0.02% WLNWCWL XRXLLLRC* 16

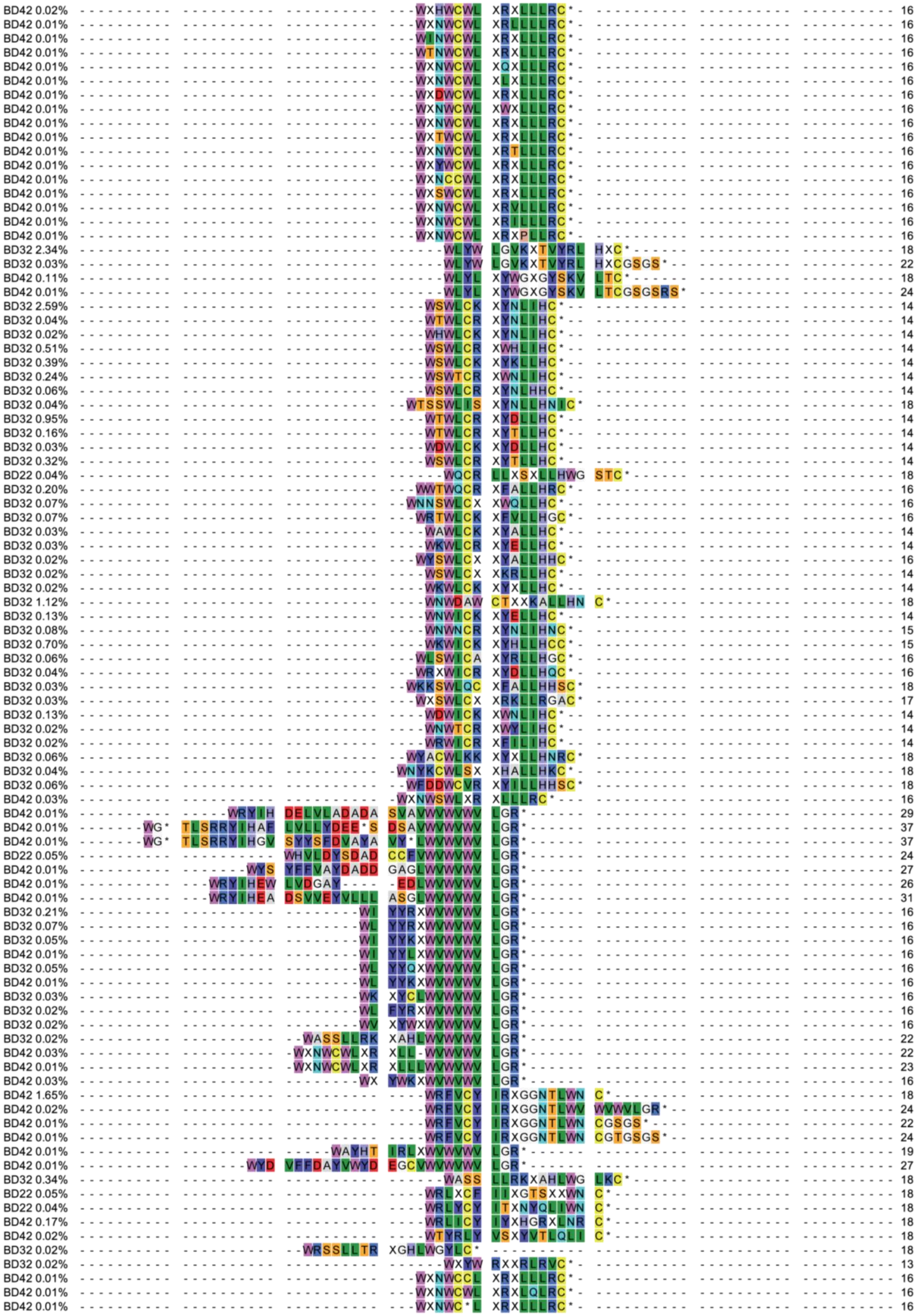

BD42 0.02% 16
BD42 0.01% 16
BD42 0.01% 16
BD42 0.01% 16
BD42 0.01% 16
BD42 0.01% 16
BD42 0.01% 16
BD42 0.01% 16
BD42 0.01% 16
BD42 0.01% 16
BD42 0.01% 16
BD42 0.01% 16
BD42 0.01% 16
BD42 0.01% 16
BD42 0.01% 16
BD42 0.01% 16
BD42 0.01% 16
BD32 2.34% 18
BD32 0.03% 22
BD42 0.11% 18
BD42 0.01% 24
BD32 2.59% 14
BD32 0.04% 14
BD32 0.02% 14
BD32 0.51% 14
BD32 0.39% 14
BD32 0.24% 14
BD32 0.06% 14
BD32 0.04% 18
BD32 0.95% 14
BD32 0.16% 14
BD32 0.03% 14
BD32 0.32% 14
BD22 0.04% 18
BD32 0.20% 16
BD32 0.07% 16
BD32 0.07% 16
BD32 0.03% 14
BD32 0.03% 14
BD32 0.02% 16
BD32 0.02% 14
BD32 0.02% 14
BD32 1.12% 18
BD32 0.13% 14
BD32 0.08% 15
BD32 0.70% 15
BD32 0.06% 16
BD32 0.04% 16
BD32 0.03% 18
BD32 0.03% 17
BD32 0.13% 14
BD32 0.02% 14
BD32 0.02% 14
BD32 0.06% 18
BD32 0.04% 18
BD32 0.06% 18
BD42 0.03% 16
BD42 0.01% 29
BD42 0.01% 37
BD42 0.01% 37
BD22 0.05% 24
BD42 0.01% 27
BD42 0.01% 26
BD42 0.01% 31
BD32 0.21% 16
BD32 0.07% 16
BD32 0.05% 16
BD42 0.01% 16
BD32 0.05% 16
BD42 0.01% 16
BD32 0.03% 16
BD32 0.02% 16
BD32 0.02% 16
BD32 0.02% 22
BD42 0.03% 22
BD42 0.01% 23
BD42 0.03% 16
BD42 1.65% 18
BD42 0.02% 24
BD42 0.01% 22
BD42 0.01% 24
BD42 0.01% 19
BD42 0.01% 27
BD32 0.34% 18
BD22 0.05% 18
BD22 0.04% 18
BD42 0.17% 18
BD42 0.02% 18
BD32 0.02% 18
BD32 0.02% 13
BD42 0.01% 16
BD42 0.01% 16
BD42 0.01% 16

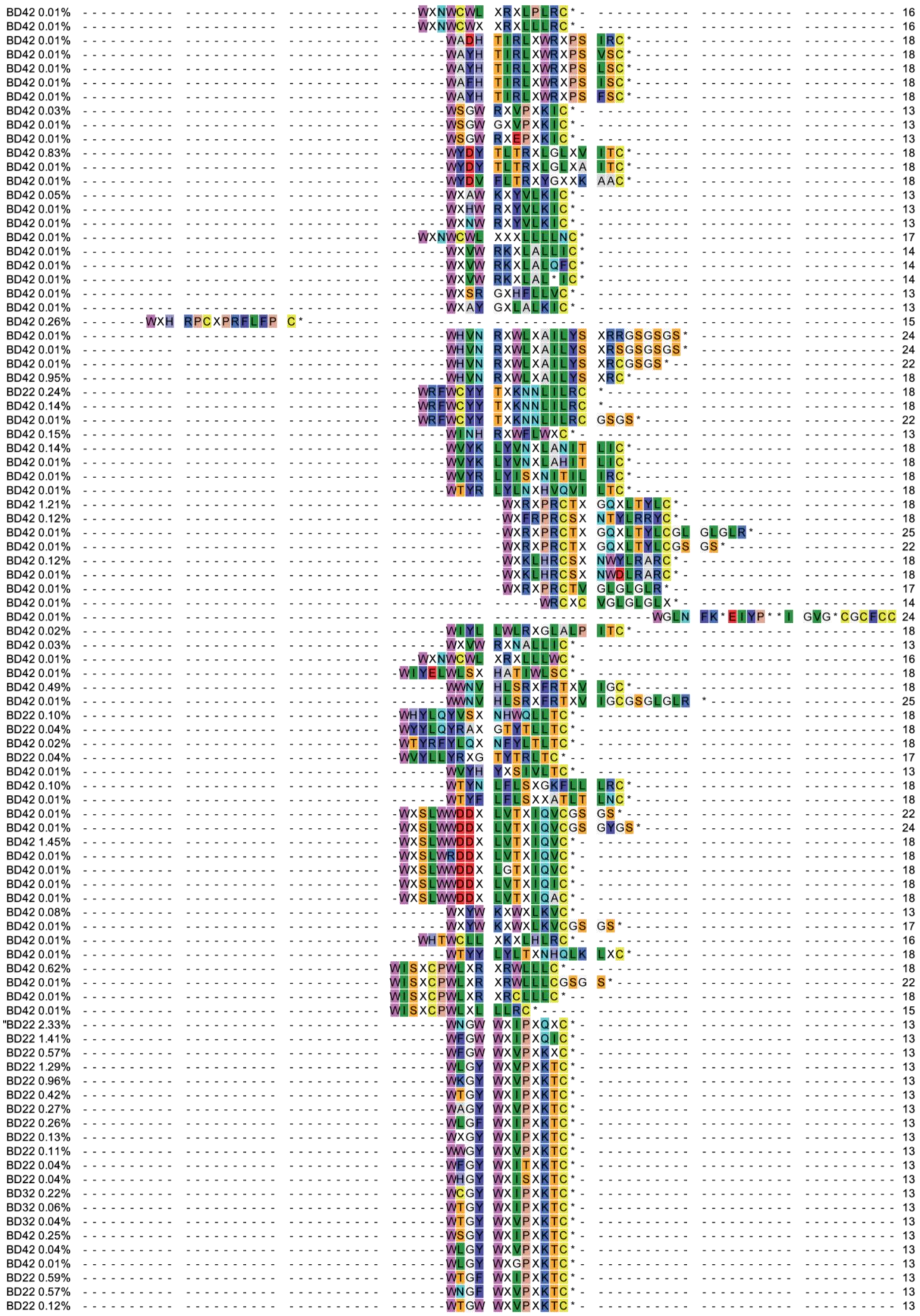
BD42 0.01% WXNWCWL XRXLPLRC* 16
BD42 0.01% WXNWCWX XRXLLLRC* 16
BD42 0.01% WADH TIRLXWRXPS IRC* 18
BD42 0.01% WAYH TIRLXWRXPS VSC* 18
BD42 0.01% WAYH TIRLXWRXPS LSC* 18
BD42 0.01% WAFH TIRLXWRXPS ISC* 18
BD42 0.01% WAYH TIRLXWRXPS FSC* 18
BD42 0.03% WSGW RXVPXKIC* 13
BD42 0.01% WSGW GXVPXKIC* 13
BD42 0.01% WSGW RXEPXKIC* 13
BD42 0.83% WYDY TLTRXLGLXV ITC* 18
BD42 0.01% WYDY TLTRXLGLXA ITC* 18
BD42 0.01% WYDV FLTRXYGXXK AAC* 18
BD42 0.05% WXAW KXYVLKIC* 13
BD42 0.01% WXHW RXYVLKIC* 13
BD42 0.01% WXNW RXYVLKIC* 13
BD42 0.01% WXNWCWL XXXLLLNC* 17
BD42 0.01% WXVW RKXLALLIC* 14
BD42 0.01% WXVW RKXLALQFC* 14
BD42 0.01% WXVW RKXLAL*IC* 14
BD42 0.01% WXSR GXHFLLVC* 13
BD42 0.01% WXAY GXLALKIC* 13
BD42 0.26% WXH RPCXPRFLFP C* 15
BD42 0.01% WHVN RXWLXAILYS XRRGSGSGS* 24
BD42 0.01% WHVN RXWLXAILYS XRSGSGSGS* 24
BD42 0.01% WHVN RXWLXAILYS XRCGSGS* 22
BD42 0.95% WHVN RXWLXAILYS XRC* 18
BD22 0.24% WRFWCYY TXKNNLILRC * 18
BD42 0.14% WRFWCYY TXKNNLILRC * 18
BD42 0.01% WRFWCYY TXKNNLILRC GSGS* 22
BD42 0.15% WINH RXWFLWXC* 13
BD42 0.14% WVYK LYVNXLANIT LIC* 18
BD42 0.01% WVYK LYVNXLAHIT LIC* 18
BD42 0.01% WVYR LYISXNITIL IRC* 18
BD42 0.01% WTYR LYLNXHVQVI LTC* 18
BD42 1.21% WXRXPRCTX GQXLTYLC* 18
BD42 0.12% WXFRPRCSX NTYLRRYC* 18
BD42 0.01% WXRXPRCTX GQXLTYLCGL GLGLR* 25
BD42 0.01% WXRXPRCTX GQXLTYLCGS GS* 22
BD42 0.12% WXKLHRCSX NWYLRARC* 18
BD42 0.01% WXKLHRCSX NWDLRARC* 18
BD42 0.01% WXRXPRCTV GLGLGLR* 17
BD42 0.01% WRCXC VGLGLGLX* 14
BD42 0.01% WGLN FK*EIYP**I GVG*CGCFCC 24
BD42 0.02% WIYL LWLRXGLALP ITC* 18
BD42 0.03% WXVW RXNALLIC* 13
BD42 0.01% WXNWCWL XRXLLLWC* 16
BD42 0.01% WIYELWLSX HATIWLSC* 18
BD42 0.49% WWNV HLSRXFRTXV IGC* 18
BD42 0.01% WWNV HLSRXFRTXV IGCGSGLGLR * 25
BD22 0.10% WHYLQYVSX NHWQLLTC* 18
BD22 0.04% WYYLQYRAX GTYTLLTC* 18
BD42 0.02% WTYRFYLQX NFYLTLTC* 18
BD22 0.04% WVYLLYRXG TYTRLTC* 17
BD42 0.01% WVYH YXSIVLTC* 13
BD42 0.10% WTYN LFLSXGKFLL LRC* 18
BD42 0.01% WTYF LFLSXXATLT LNC* 18
BD42 0.01% WXSLWWDDX LVTXIQVCGS GS* 22
BD42 0.01% WXSLWWDDX LVTXIQVCGS GYGS* 24
BD42 1.45% WXSLWWDDX LVTXIQVC* 18
BD42 0.01% WXSLWRDDX LVTXIQVC* 18
BD42 0.01% WXSLWWDDX LGTXIQVC* 18
BD42 0.01% WXSLWWDDX LVTXIQIC* 18
BD42 0.01% WXSLWWDDX LVTXIQAC* 18
BD42 0.08% WXYW KXWXLKVC* 13
BD42 0.01% WXYW KXWXLKVCGS GS* 17
BD42 0.01% WHTWCLL XKXLHLRC* 16
BD42 0.01% WTYY LYLTXNHQLK LXC* 18
BD42 0.62% WISXCPWLXR XRWLLLC* 18
BD42 0.01% WISXCPWLXR XRWLLLCGSG S* 22
BD42 0.01% WISXCPWLXR XRCLLLC* 18
BD42 0.01% WISXCPWLXL LLRC* 15
"BD22 2.33% WNGW WXIPXQXC* 13
BD22 1.41% WFGW WXIPXQIC* 13
BD22 0.57% WFGW WXVPXKXC* 13
BD22 1.29% WLGY WXVPXKTC* 13
BD22 0.96% WKGY WXVPXKTC* 13
BD22 0.42% WTGY WXIPXKTC* 13
BD22 0.27% WAGY WXVPXKTC* 13
BD22 0.26% WLGF WXIPXKTC* 13
BD22 0.13% WXGY WXIPXKTC* 13
BD22 0.11% WWGY WXVPXKTC* 13
BD22 0.04% WFGY WXITXKTC* 13
BD22 0.04% WHGY WXISXKTC* 13
BD32 0.22% WCGY WXIPXKTC* 13
BD32 0.06% WTGY WXIPXKTC* 13
BD32 0.04% WTGY WXVPXKTC* 13
BD42 0.25% WSGY WXIPXKTC* 13
BD42 0.04% WLGY WXVPXKTC* 13
BD42 0.01% WLGY WXGPXKTC* 13
BD22 0.59% WTGF WXIPXKTC* 13
BD22 0.57% WNGF WXVPXKTC* 13
BD22 0.12% WTGW WXVPXKTC* 13

| | | |
|---|---|---|
| BD22 0.10% | WTGW WXIAXKTC* | 13 |
| BD22 0.07% | WEGF WXVPXKTC* | 13 |
| BD22 0.56% | WYGT WXVPXKTC* | 13 |
| BD22 0.31% | WHGT WXVPXKTC* | 13 |
| BD22 0.20% | WHGT WXIPXKTC* | 13 |
| BD22 0.17% | WYGW WXIVXKTC* | 13 |
| BD22 0.08% | WYGN WXIPXKTC* | 13 |
| BD22 0.05% | WYGW WXQPXKTC* | 13 |
| BD22 0.50% | WSGW WXVPXKTC* | 13 |
| BD22 0.24% | WHGW WXVSXKTC* | 13 |
| BD22 0.21% | WHGW WXVTXKTC* | 13 |
| BD22 0.21% | WRGW WXIPXKTC* | 13 |
| BD22 0.05% | WFGW WXKPXKTC* | 13 |
| BD22 0.05% | WRGW WXVPXKTC* | 13 |
| BD22 0.04% | WFGW WXVPXKTC* | 13 |
| BD32 0.02% | WCGW WXIAXKTC* | 13 |
| BD22 0.56% | WNGY WXIPXRTC* | 13 |
| BD22 0.51% | WSGW WXIPXXTC* | 13 |
| BD22 0.46% | WYGW WXVPXXQC* | 13 |
| BD22 0.37% | WNGW WXIAXKRC* | 13 |
| BD22 0.14% | WHGW WXIAXKRC* | 13 |
| BD22 0.10% | WHGW WXVPXKRC* | 13 |
| BD22 0.08% | WHGW WXINXKRC* | 13 |
| BD22 0.04% | WNGS WXIPXKRC* | 13 |
| BD22 0.31% | WSGW WXTPXKKC* | 13 |
| BD22 0.07% | WSGW WXVIXKKC* | 13 |
| BD22 0.82% | WSGY WXVPXRLC* | 13 |
| BD22 0.21% | WNGY WXIPXRLC* | 13 |
| BD22 0.18% | WSGY WXIPXRLC* | 13 |
| BD22 0.07% | WAGY WXIPXRLC* | 13 |
| BD22 0.05% | WCGY WXIPXRLC* | 13 |
| BD22 0.05% | WSGW WXIPXRLC* | 13 |
| BD22 0.04% | WFGY WXXPXRLC* | 13 |
| BD22 0.04% | WCGY WXVAXRLC* | 13 |
| BD22 0.04% | WSGT WXIPXRLC* | 13 |
| BD42 0.03% | WSGW WXIPXRLC* | 13 |
| BD22 0.29% | WNGW WXVPXRLC* | 13 |
| BD22 0.21% | WDGW WXIPXRLC* | 13 |
| BD22 0.09% | WFGW WXIPXRLC* | 13 |
| BD22 0.08% | WHGW WXISXRLC* | 13 |
| BD22 0.04% | WNGW WXITXRLC* | 13 |
| BD22 0.20% | WNGW WXIAXKLC* | 13 |
| BD22 0.12% | WNGT WXVPXKLC* | 13 |
| BD22 0.11% | WFGW WXQPXKLC* | 13 |
| BD22 0.08% | WNGT WXIPXKLC* | 13 |
| BD22 0.07% | WHGW WXQPXKLC* | 13 |
| BD22 0.05% | WNGY WXIVXKLC* | 13 |
| BD22 0.04% | WQGW WXISXKLC* | 13 |
| BD22 0.04% | WDGW WXKPXKLC* | 13 |
| BD42 0.01% | WNGT WXIPXKLC* | 13 |
| BD22 0.12% | WSGY WXIAXKLC* | 13 |
| BD22 0.08% | WKGY WXVPXKLC* | 13 |
| BD22 0.10% | WHGS WXIPXKLC* | 13 |
| BD22 0.18% | WYGW WXVAXKQC* | 13 |
| BD22 0.07% | WYGN WXIPXKQC* | 13 |
| BD22 0.18% | WNGW WXVPXQXC* | 13 |
| BD22 0.17% | WNGY WXVPXXLC* | 13 |
| BD22 0.17% | WAGW WXIAXQKC* | 13 |
| BD22 0.04% | WNGW WXXPXQKC* | 13 |
| BD22 0.14% | WSGW WXVPXAQC* | 13 |
| BD22 0.13% | WGGW WXVPXRXC* | 13 |
| BD22 0.13% | WNGW WXVPXSLC* | 13 |
| BD22 0.12% | WSGW WXIPXIRC* | 13 |
| BD22 0.07% | WNGW WXLPXIRC* | 13 |
| BD22 0.25% | WFGF WXIPXKKC* | 13 |
| BD22 0.09% | WYGF WXIIXKKC* | 13 |
| BD22 0.05% | WGGF WXIPXKKC* | 13 |
| BD22 0.04% | WFGN WXIVXKKC* | 13 |
| BD22 0.04% | WFGS WXIVXKKC* | 13 |
| BD22 0.18% | WYGT WXIVXKKC* | 13 |
| BD22 0.12% | WYGY WXISXKKC* | 13 |
| BD22 0.11% | WNGT WXIPXKKC* | 13 |
| BD22 0.16% | WVGY WXIPXKKC* | 13 |
| BD22 0.10% | WHGY WXISXKKC* | 13 |
| BD22 0.09% | WCGY WXTPXKKC* | 13 |
| BD22 0.04% | WKGY WXIVXKKC* | 13 |
| BD42 0.07% | WTGY WXIPXKKC* | 13 |
| BD22 0.11% | WNGY WXIPXALC* | 13 |
| BD22 0.08% | WNGY WXVPXRXC* | 13 |
| BD22 0.08% | WNGV WXIPXKXC* | 13 |
| BD22 0.08% | WNGT WXIPXXRC* | 13 |
| BD22 0.08% | WNGW WXIPXVXC* | 13 |
| BD22 0.07% | WNGW WXINX*KC* | 13 |
| BD22 0.07% | WNGY WXIPXKSC* | 13 |
| BD22 0.06% | WYGW WXTPXKXC* | 13 |
| BD22 0.05% | WTGW WXIAXKVC* | 13 |
| BD22 0.05% | WNGY WXVPXKAC* | 13 |
| BD22 0.05% | WNGW WXKPXQLC* | 13 |
| BD22 0.05% | WFGW WXIPXSHC* | 13 |
| BD22 0.04% | WFGW WXIAXKAC* | 13 |
| BD22 0.04% | WNGW WXIPXVSC* | 13 |
| BD22 0.04% | WAGW WXIVXRKC* | 13 |
| BD32 0.14% | WNGCDPWLXX XFRLLHC* | 18 |

| | | |
|---|---|---|
| BD32 0.14% | WNGW WXIPXKXC* | 13 |
| BD32 0.05% | WSGW WXIPXRIC* | 13 |
| BD42 0.15% | WNGQRXWLXA IFYAVKC* | 18 |
| BD22 0.25% | WWII PXKLAAAC* | 13 |
| BD22 0.11% | WWII PXRLAAKC* | 13 |
| BD22 0.11% | WWII PXRVAAAC* | 13 |
| BD22 0.05% | WWII PXRLAANC* | 13 |
| BD22 0.05% | WWII PXXLAAIC* | 13 |
| BD22 0.04% | WWII PXVLAARC* | 13 |
| BD22 0.04% | WWII PXQLAAKC* | 13 |
| BD22 0.04% | WWII PXRLAADC* | 13 |
| BD22 0.23% | WWII PXKXAAKC* | 13 |
| BD22 0.12% | WWII PXKIAAKC* | 13 |
| BD22 0.11% | WWII PXQKAAKC* | 13 |
| BD22 0.09% | WWII PXRKAAKC* | 13 |
| BD22 0.05% | WWII PXXKAAKC* | 13 |
| BD22 0.04% | WWII PX*IAAKC* | 13 |
| BD22 0.21% | WWII PXQKAAYC* | 13 |
| BD22 0.07% | WWIV PXKKAATC* | 13 |
| BD22 0.05% | WWII PXIKAANC* | 13 |
| BD22 0.05% | WWII PXQKAAIC* | 13 |
| BD22 0.20% | WWII PXKXAATC* | 13 |
| BD22 0.05% | WWII PXRVAATC* | 13 |
| BD22 0.17% | WWII PXXKAHTQC* | 14 |
| BD22 0.14% | WWII PXKKAHSC* | 13 |
| BD22 0.08% | WWII PXKKAHIC* | 13 |
| BD22 0.07% | WWII PXKKAHHC* | 13 |
| BD22 0.07% | WWIV PXKKAHDC* | 13 |
| BD22 0.05% | WWII PXVKAHNC* | 13 |
| BD22 0.04% | WWII PXVKAHWC* | 13 |
| BD22 0.04% | WWIV PXKKAHNC* | 13 |
| BD22 0.06% | WWII PXKKASPC* | 13 |
| BD22 0.05% | WWII PXKKAQQC* | 13 |
| BD22 0.05% | WWII PXKKAQDC* | 13 |
| BD22 0.04% | WWII PXKKAYNC* | 13 |
| BD22 0.04% | WWII PXKKAQSC* | 13 |
| BD22 0.04% | WWII PXKKARNC* | 13 |
| BD22 0.16% | WWII PXKRAASC* | 13 |
| BD22 0.14% | WWII PXRIAASC* | 13 |
| BD22 0.05% | WWII PXKIAASC* | 13 |
| BD22 0.05% | WWII PXKRAANC* | 13 |
| BD22 0.04% | WWII PXKRAAFC* | 13 |
| BD22 0.11% | WWII PXKIAANC* | 13 |
| BD22 0.11% | WWII PXKIAAPC* | 13 |
| BD22 0.07% | WWII PXXIAAQC* | 13 |
| BD22 0.13% | WWVI PXKKAHNC* | 13 |
| BD22 0.04% | WWVI PXKKAHVC* | 13 |
| BD22 0.04% | WWVI PXKKARSC* | 13 |
| BD22 0.08% | WWII PXKIAHIC* | 13 |
| BD22 0.23% | WWII PXKXAGC* | 12 |
| BD22 0.08% | WWII PXVKAGC* | 12 |
| BD22 0.05% | WWIV PXKIAGC* | 12 |
| BD42 0.02% | WWIV PXKIAGC* | 12 |
| BD42 0.01% | WWIV PXXVAGC* | 12 |
| BD22 0.05% | WWII PXKIAFHC* | 13 |
| BD22 0.04% | WWII PXKIAYQC* | 13 |
| BD22 0.07% | WWVI PXKIAANC* | 13 |
| BD22 0.05% | WWII PXKXAANC* | 13 |
| BD22 0.04% | WWII PXRLAHHC* | 13 |
| BD22 0.07% | WGFWII VXQRAGYAPC * | 17 |
| BD22 0.11% | WWIV PXKTAAQC* | 13 |
| BD22 0.10% | WWVI PXKTAAQC* | 13 |
| BD22 0.05% | WWII PXKTAALC* | 13 |
| BD22 0.04% | WWII PXKTAHPC* | 13 |
| BD22 0.04% | WWII PXKTAGHC* | 13 |
| BD22 0.15% | WWVI PXXKAGC* | 12 |
| BD22 0.07% | WWVI PXKTAGC* | 12 |
| BD42 0.02% | WWVI PXKIAGC* | 12 |
| BD22 0.04% | WYWVI PXKTAAKHC* | 15 |
| BD22 0.10% | WWII PXXKAGLC* | 13 |
| BD22 0.09% | WWVI PXKIAQAC* | 13 |
| BD22 0.07% | WWVI PXKIAYPC* | 13 |
| BD22 0.04% | WWVV PXKIAASC* | 13 |
| BD22 0.04% | WWVI PXKIAARC* | 13 |
| BD22 0.09% | WWII PXVLAHPC* | 13 |
| BD22 0.06% | WWII PXXLAHNGC* | 14 |
| BD22 0.04% | WWIV PXXLAHYSC* | 14 |
| BD22 0.06% | WWII PXVKAYHC* | 13 |
| BD22 0.33% | WWLI PXKKAISC* | 13 |
| BD22 0.14% | WWLI PXKKAANC* | 13 |
| BD22 0.12% | WYWLI PXKKAALC* | 14 |
| BD22 0.09% | WWLI PXKKAVAC* | 13 |
| BD22 0.09% | WWLI PXKKAFPC* | 13 |
| BD22 0.07% | WWLI PXVKAIPC* | 13 |
| BD22 0.06% | WWLI PXKKAIPC* | 13 |
| BD22 0.06% | WWLI PXKKAHAC* | 13 |
| BD22 0.05% | WWLI PXKKAXPC* | 13 |
| BD22 0.04% | WWLI PXKKAIAC* | 13 |
| BD22 0.04% | WWLI PXKKAINC* | 13 |
| BD22 0.04% | WWLI PXKKAVSC* | 13 |
| BD22 0.04% | WWLI PXKKAXDC* | 13 |
| BD22 0.04% | WWLI PXVKAISC* | 13 |
| BD22 0.18% | WWLI PXKTAIPC* | 13 |

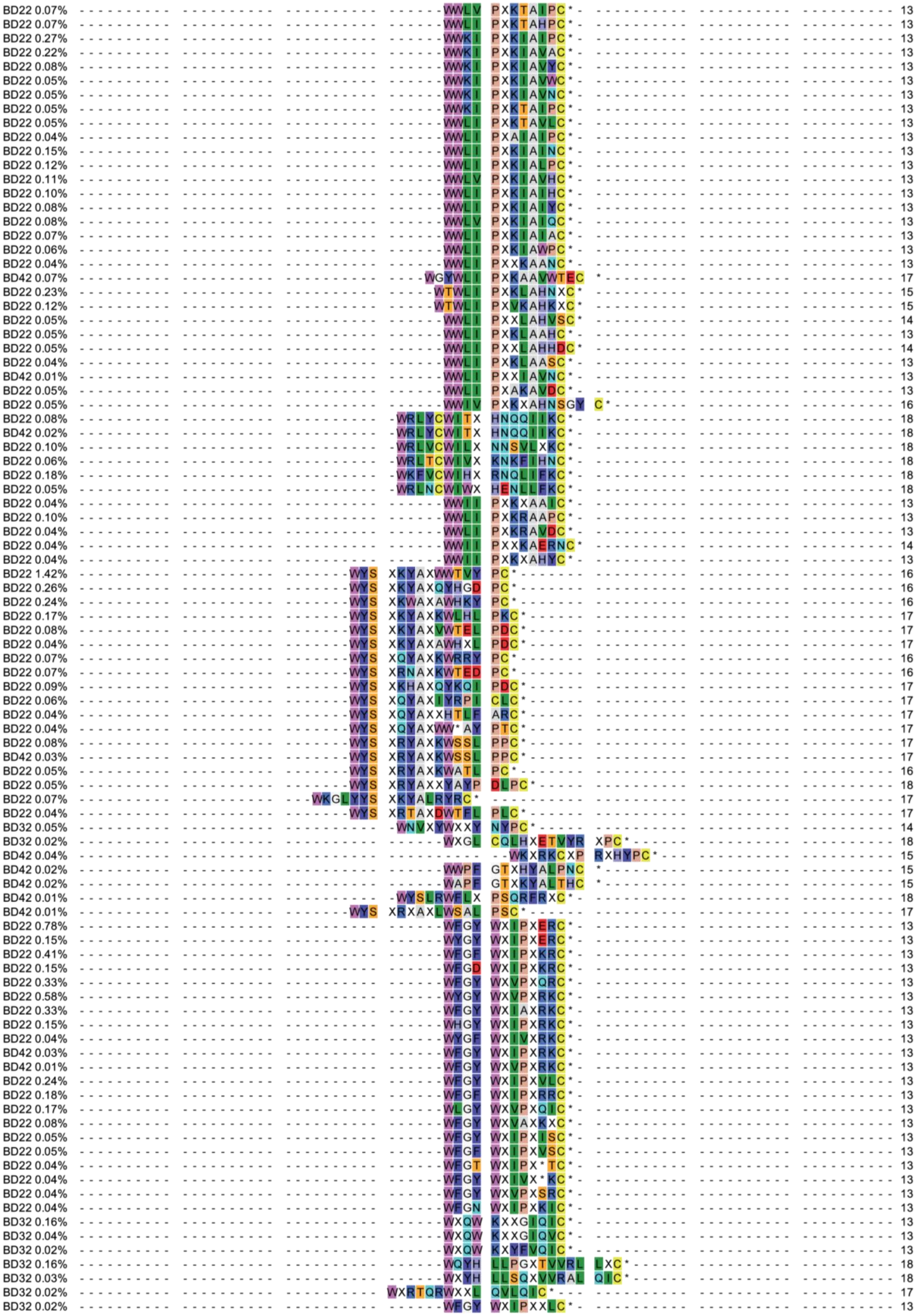

BD22 0.07% 13
BD22 0.07% 13
BD22 0.27% 13
BD22 0.22% 13
BD22 0.08% 13
BD22 0.05% 13
BD22 0.05% 13
BD22 0.05% 13
BD22 0.05% 13
BD22 0.04% 13
BD22 0.15% 13
BD22 0.12% 13
BD22 0.11% 13
BD22 0.10% 13
BD22 0.08% 13
BD22 0.08% 13
BD22 0.07% 13
BD22 0.06% 13
BD22 0.04% 13
BD42 0.07% 17
BD22 0.23% 15
BD22 0.12% 15
BD22 0.05% 14
BD22 0.05% 13
BD22 0.05% 14
BD22 0.04% 13
BD42 0.01% 13
BD22 0.05% 13
BD22 0.05% 16
BD22 0.08% 18
BD42 0.02% 18
BD22 0.10% 18
BD22 0.06% 18
BD22 0.18% 18
BD22 0.05% 18
BD22 0.04% 13
BD22 0.10% 13
BD22 0.04% 13
BD22 0.04% 14
BD22 0.04% 13
BD22 1.42% 16
BD22 0.26% 16
BD22 0.24% 16
BD22 0.17% 17
BD22 0.08% 17
BD22 0.04% 17
BD22 0.07% 16
BD22 0.07% 16
BD22 0.09% 17
BD22 0.06% 17
BD22 0.04% 17
BD22 0.04% 17
BD22 0.08% 17
BD42 0.03% 17
BD22 0.05% 16
BD22 0.05% 18
BD22 0.07% 17
BD22 0.04% 17
BD32 0.05% 14
BD32 0.02% 18
BD42 0.04% 15
BD42 0.02% 15
BD42 0.02% 15
BD42 0.01% 18
BD42 0.01% 17
BD22 0.78% 13
BD22 0.15% 13
BD22 0.41% 13
BD22 0.15% 13
BD22 0.33% 13
BD22 0.58% 13
BD22 0.33% 13
BD22 0.15% 13
BD22 0.04% 13
BD42 0.03% 13
BD42 0.01% 13
BD22 0.24% 13
BD22 0.18% 13
BD22 0.17% 13
BD22 0.08% 13
BD22 0.05% 13
BD22 0.05% 13
BD22 0.04% 13
BD22 0.04% 13
BD22 0.04% 13
BD22 0.04% 13
BD32 0.16% 13
BD32 0.04% 13
BD32 0.02% 13
BD32 0.16% 18
BD32 0.03% 18
BD32 0.02% 17
BD32 0.02% 13

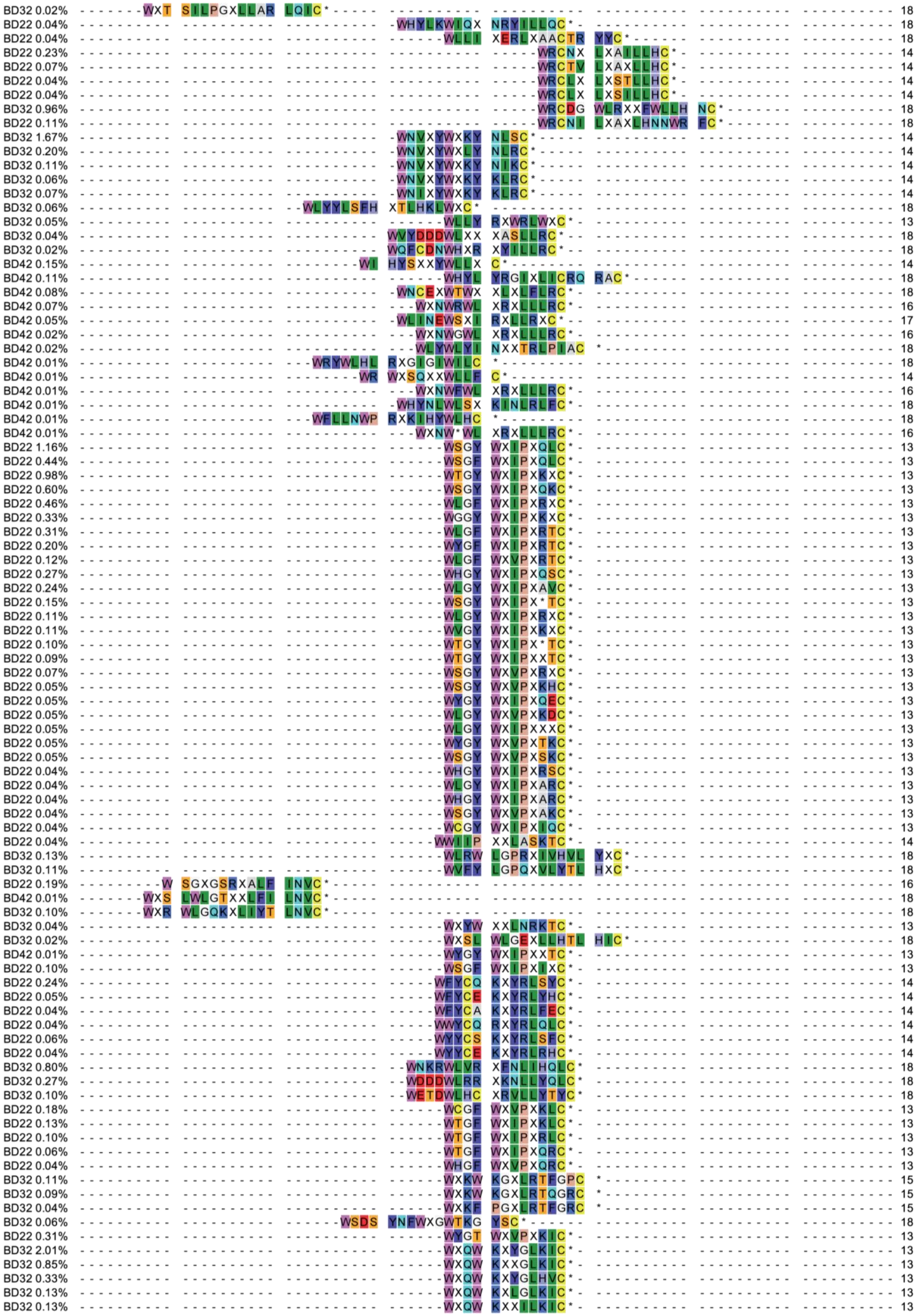
BD32 0.02% 18
BD22 0.04% 18
BD22 0.04% 18
BD22 0.23% 14
BD22 0.07% 14
BD22 0.04% 14
BD22 0.04% 14
BD32 0.96% 18
BD22 0.11% 18
BD32 1.67% 14
BD32 0.20% 14
BD32 0.11% 14
BD32 0.06% 14
BD32 0.07% 14
BD32 0.06% 18
BD32 0.05% 13
BD32 0.04% 18
BD32 0.02% 18
BD42 0.15% 14
BD42 0.11% 18
BD42 0.08% 18
BD42 0.07% 16
BD42 0.05% 17
BD42 0.02% 16
BD42 0.02% 18
BD42 0.01% 18
BD42 0.01% 14
BD42 0.01% 16
BD42 0.01% 18
BD42 0.01% 18
BD42 0.01% 16
BD22 1.16% 13
BD22 0.44% 13
BD22 0.98% 13
BD22 0.60% 13
BD22 0.46% 13
BD22 0.33% 13
BD22 0.31% 13
BD22 0.20% 13
BD22 0.12% 13
BD22 0.27% 13
BD22 0.24% 13
BD22 0.15% 13
BD22 0.11% 13
BD22 0.11% 13
BD22 0.10% 13
BD22 0.09% 13
BD22 0.07% 13
BD22 0.05% 13
BD22 0.05% 13
BD22 0.05% 13
BD22 0.05% 13
BD22 0.05% 13
BD22 0.05% 13
BD22 0.04% 13
BD22 0.04% 13
BD22 0.04% 13
BD22 0.04% 13
BD22 0.04% 13
BD22 0.04% 14
BD32 0.13% 18
BD32 0.11% 18
BD22 0.19% 16
BD42 0.01% 18
BD32 0.10% 18
BD32 0.04% 13
BD32 0.02% 18
BD42 0.01% 13
BD22 0.10% 13
BD22 0.24% 14
BD22 0.05% 14
BD22 0.04% 14
BD22 0.04% 14
BD22 0.06% 14
BD22 0.04% 14
BD32 0.80% 18
BD32 0.27% 18
BD32 0.10% 18
BD22 0.18% 13
BD22 0.13% 13
BD22 0.10% 13
BD22 0.06% 13
BD22 0.04% 13
BD32 0.11% 15
BD32 0.09% 15
BD32 0.04% 15
BD32 0.06% 18
BD22 0.31% 13
BD32 2.01% 13
BD32 0.85% 13
BD32 0.33% 13
BD32 0.13% 13
BD32 0.13% 13

| | | | |
|---|---|---|---|
| BD32 | 0.06% | WXQW KXYFTKIC* | 13 |
| BD32 | 0.05% | WXQW KXYILKIC* | 13 |
| BD32 | 0.04% | WXKW KXYGLKIC* | 13 |
| BD32 | 0.03% | WXQW KXRFLKIC* | 13 |
| BD42 | 0.03% | WXQW KXYFIKIC* | 13 |
| BD32 | 0.25% | WXQW KXHFLKVC* | 13 |
| BD32 | 0.21% | WXQW KXHFLNIC* | 13 |
| BD32 | 0.19% | WLQW KXYFLKXC* | 13 |
| BD32 | 0.07% | WXQW KXYSLKVC* | 13 |
| BD32 | 0.06% | WXQW KXXGLNIC* | 13 |
| BD42 | 0.01% | WLQW KXYILKXC* | 13 |
| BD32 | 3.10% | WXK AILPGXILKT LHIC* | 18 |
| BD32 | 0.98% | WXR HILPGXRLKT LKIC* | 18 |
| BD32 | 0.19% | WXRW KXFGLKIC* | 13 |
| BD32 | 0.03% | WXRW KXYALKIC* | 13 |
| BD32 | 0.18% | WXYH LGAXXSVHSL KIC* | 18 |
| BD32 | 0.06% | WXFH LGPAXVLWSV HIC* | 18 |
| BD32 | 0.10% | WXTR XWXRALKIC* | 14 |
| BD32 | 0.06% | WXFY RXWGLKIC* | 13 |
| BD32 | 0.06% | WXSW KXYFLKIC* | 13 |
| BD32 | 0.03% | WXQW RXHILKIC* | 13 |
| BD42 | 0.13% | WXHW XXYTLKIC* | 13 |
| BD42 | 0.01% | WVPX KIC* | 8 |
| BD32 | 1.17% | WXHY LAAKXXILRL CVC* | 18 |
| BD32 | 0.02% | WXHY LAAKXXILRL C* | 16 |
| BD32 | 0.16% | WXG NCRIHXGAFI XRLC* | 18 |
| BD42 | 0.01% | WNSW LVPXVSVSCR LC* | 17 |
| BD32 | 0.04% | WTSQ LLQKXASLFG YTC* | 18 |
| BD32 | 0.02% | WTSD LLQRXGRLFG FHC* | 18 |
| BD32 | 0.02% | WRSR LLQXXARLFG XVC* | 18 |
| BD42 | 0.03% | WX PWGTXKWALE RC* | 15 |
| BD22 | 0.62% | WHLXCWYDX KANRIYKC* | 18 |
| BD22 | 0.28% | WRYYCWYVX HGTLISNC* | 18 |
| BD22 | 0.06% | WYDX STAXRWKRYP AC* | 17 |
| BD22 | 0.05% | WRFTCWIIX ANRXICSC* | 18 |
| BD22 | 0.04% | WRFVCWIIX KGHTLPNC* | 18 |
| BD22 | 0.04% | W ILHRILDXYR YYQYKC* | 18 |
| BD32 | 0.03% | WLYW QXYRIYXC* | 13 |
| BD42 | 0.02% | WLFK TXRIYAPC* | 13 |
| BD42 | 0.01% | WXTCWYLT XLIAVLRVC* | 18 |
| BD32 | 0.02% | WXWRGWXGX WQKC* | 14 |
| BD22 | 0.16% | WCGF WXIPXKXC* | 13 |
| BD22 | 0.12% | WYGF WXVPXRVC* | 13 |
| BD22 | 0.04% | WYGT WXIPXRVC* | 13 |
| BD22 | 0.10% | WDGF WXIPXRXC* | 13 |
| BD22 | 0.09% | WCGF WXIPXVRC* | 13 |
| BD42 | 0.01% | WYGF WXIPXIRC* | 13 |
| BD32 | 1.09% | WSKI WSVXXYNLIH RKC* | 18 |
| BD22 | 0.04% | WNCHL LXLXLHNGFR CC* | 18 |
| BD22 | 0.04% | WDGT WXIPXQXC* | 13 |
| BD22 | 0.04% | WDGT WXIPXRRC* | 13 |
| BD42 | 0.17% | WHPY GTXKWALPWC * | 15 |
| BD42 | 0.06% | WTPY GTXKKALPHC * | 15 |
| BD42 | 0.01% | WYPY GTXKWALTIC * | 15 |
| BD42 | 0.01% | WYPY GTXQKALWYC * | 15 |
| BD42 | 0.06% | WWYPY GTXAKAISCN C* | 17 |
| BD42 | 0.16% | WWWPY GTXFLAICKC * | 16 |
| BD42 | 0.01% | WWPY GTXXHARAIC * | 15 |
| BD42 | 0.02% | WWPY GTXXILINC* | 14 |
| BD42 | 0.01% | WWPY GTXXHALDXC * | 15 |
| BD42 | 0.03% | WFP YGTXHKWLXY C* | 15 |
| BD22 | 0.53% | WQYLLFSRX GXPTFLYC* | 18 |
| BD22 | 0.23% | WRYPXWRTX TQFRTLYC* | 18 |
| BD22 | 0.14% | WKCTX LXATLLFC* | 14 |
| BD22 | 0.10% | WRYL LGSXLNWITX QC* | 17 |
| BD22 | 0.08% | WQCL XLXAHLLFC* | 14 |
| BD22 | 0.06% | WRCSI LXAXLLFC* | 14 |
| BD22 | 0.04% | WRCSI LXKXLLFC* | 14 |
| BD42 | 0.04% | WRCII LXKXLLFC* | 14 |
| BD22 | 0.04% | W AVKPKXKCSL LFC* | 15 |
| BD22 | 0.04% | WRLY CFYVXRHQYV YRC* | 18 |
| BD22 | 0.04% | WYYLLYRX SNYXRLKC* | 17 |
| BD42 | 0.01% | WRYLLYIHX KARLQLKC* | 18 |
| BD32 | 1.09% | WKXYCL WRXQFAWKLY C* | 18 |
| BD32 | 0.08% | WRSY LLWKXSLQHT IKC* | 18 |
| BD32 | 0.06% | WQYH LGSQXVLFKL FXC* | 18 |
| BD32 | 0.04% | WQYY RXWXLHXC* | 13 |
| BD32 | 0.02% | WQYY RXWXLKXC* | 13 |
| BD32 | 0.02% | WQYY XXWRIKXC* | 13 |
| BD42 | 0.19% | WXTNPW*XW RYLLVC* | 16 |
| BD42 | 0.06% | WQYW RXFQLYXC* | 13 |
| BD42 | 0.03% | WISW KXFYLLXC* | 13 |
| BD42 | 0.01% | WFWX GQXXALLFC* | 14 |
| BD42 | 0.01% | WXYW XXFSRKRC* | 13 |
| BD42 | 0.01% | WIWL IPXRXTFCTF LHC* | 18 |
| BD22 | 0.49% | WTYLKW LXKVRIWVPC * | 17 |
| BD22 | 0.05% | WTYLKW XTKTRIWC* | 15 |
| BD22 | 0.12% | WTYLQW KXKHRIWIPC * | 17 |
| BD22 | 0.10% | WTYLQW XTKIRTWC* | 15 |
| BD42 | 0.49% | WYIXPSX XFXYRIWXC* | 17 |
| BD42 | 0.01% | WYIXPSX XFXYRIWXCG SGS* | 21 |
| BD32 | 0.02% | WIYH LGPKXAIISL FXC* | 18 |
| BD42 | 0.05% | WTYK YHLSXYISLT IVC* | 18 |

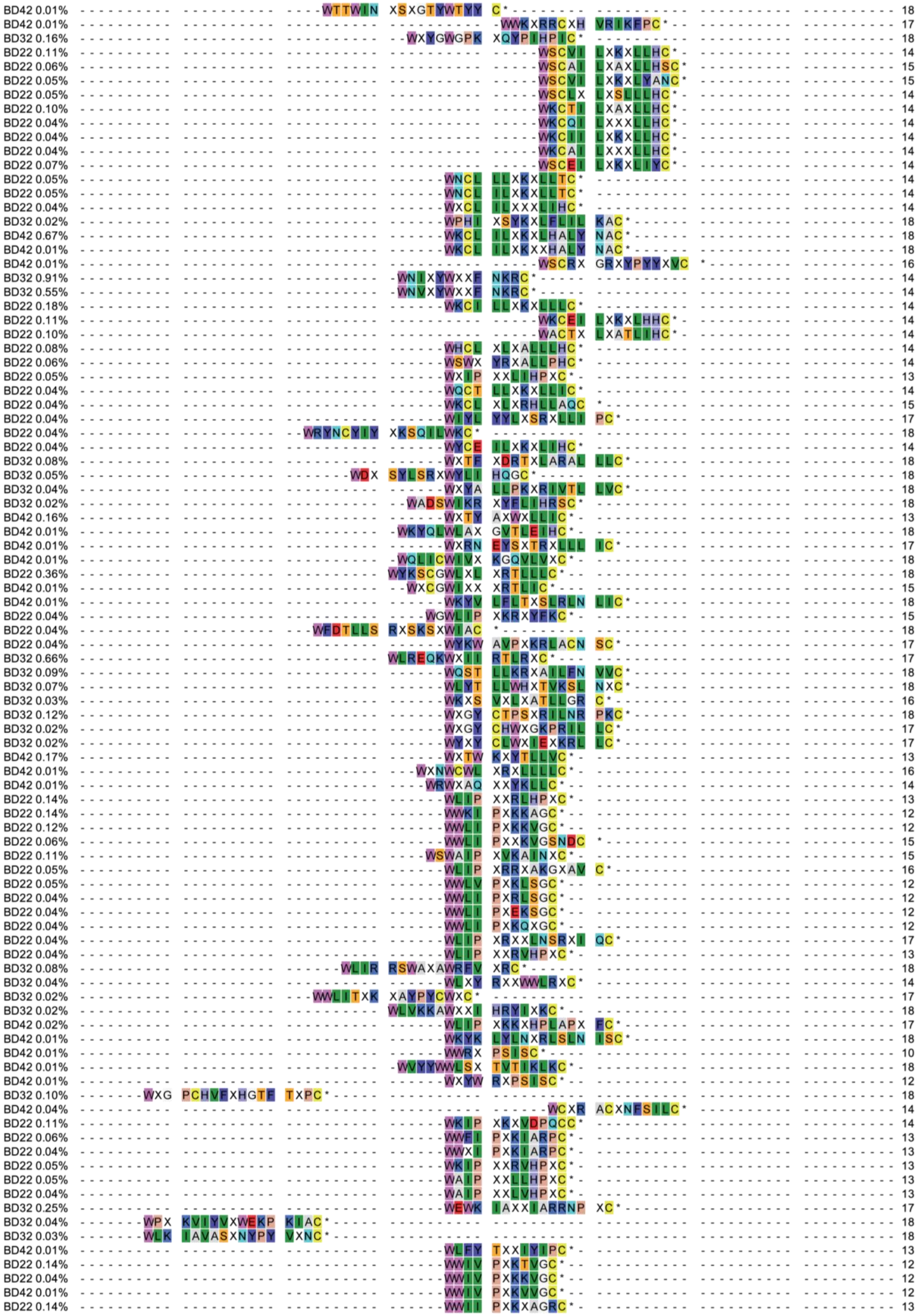

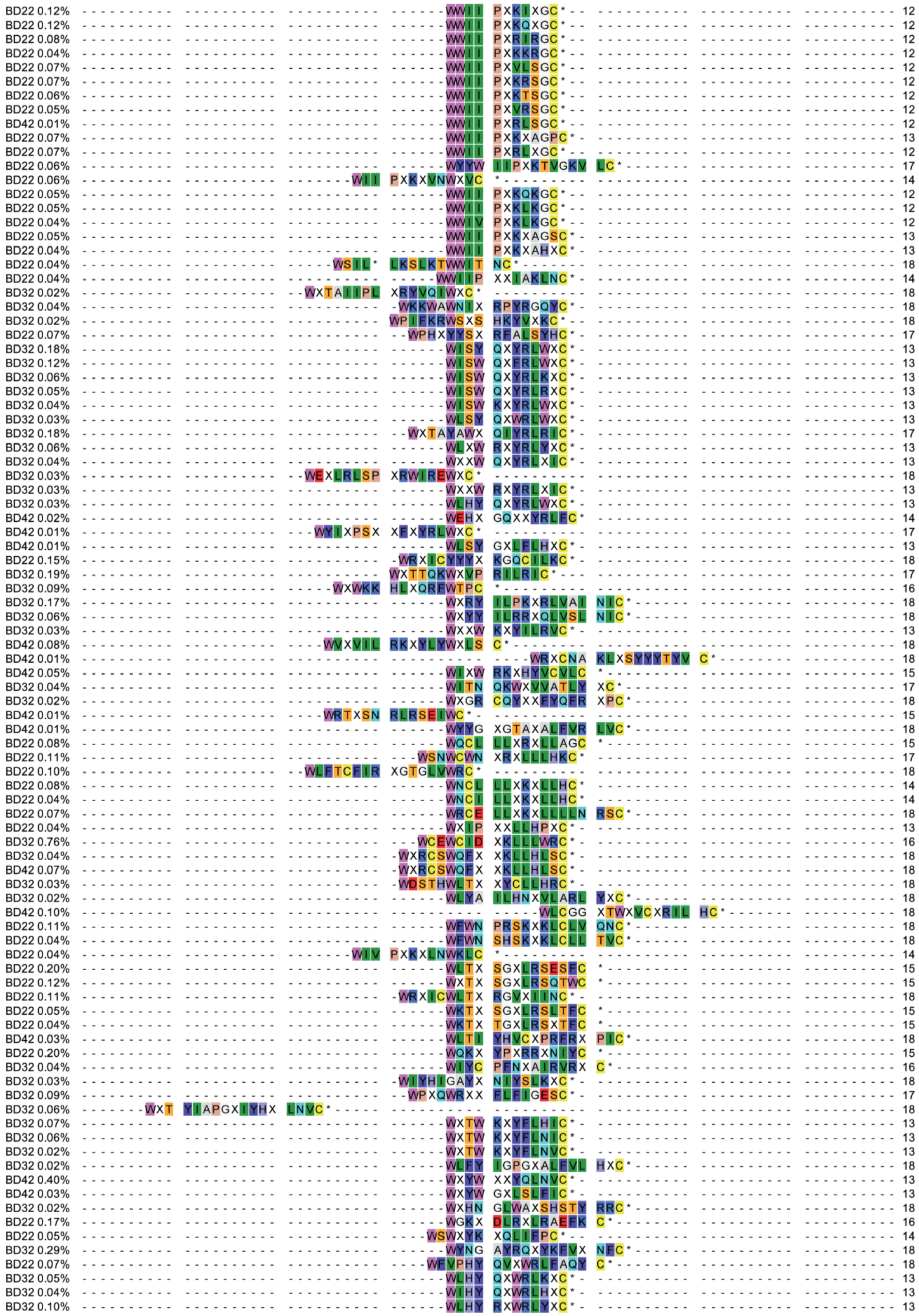

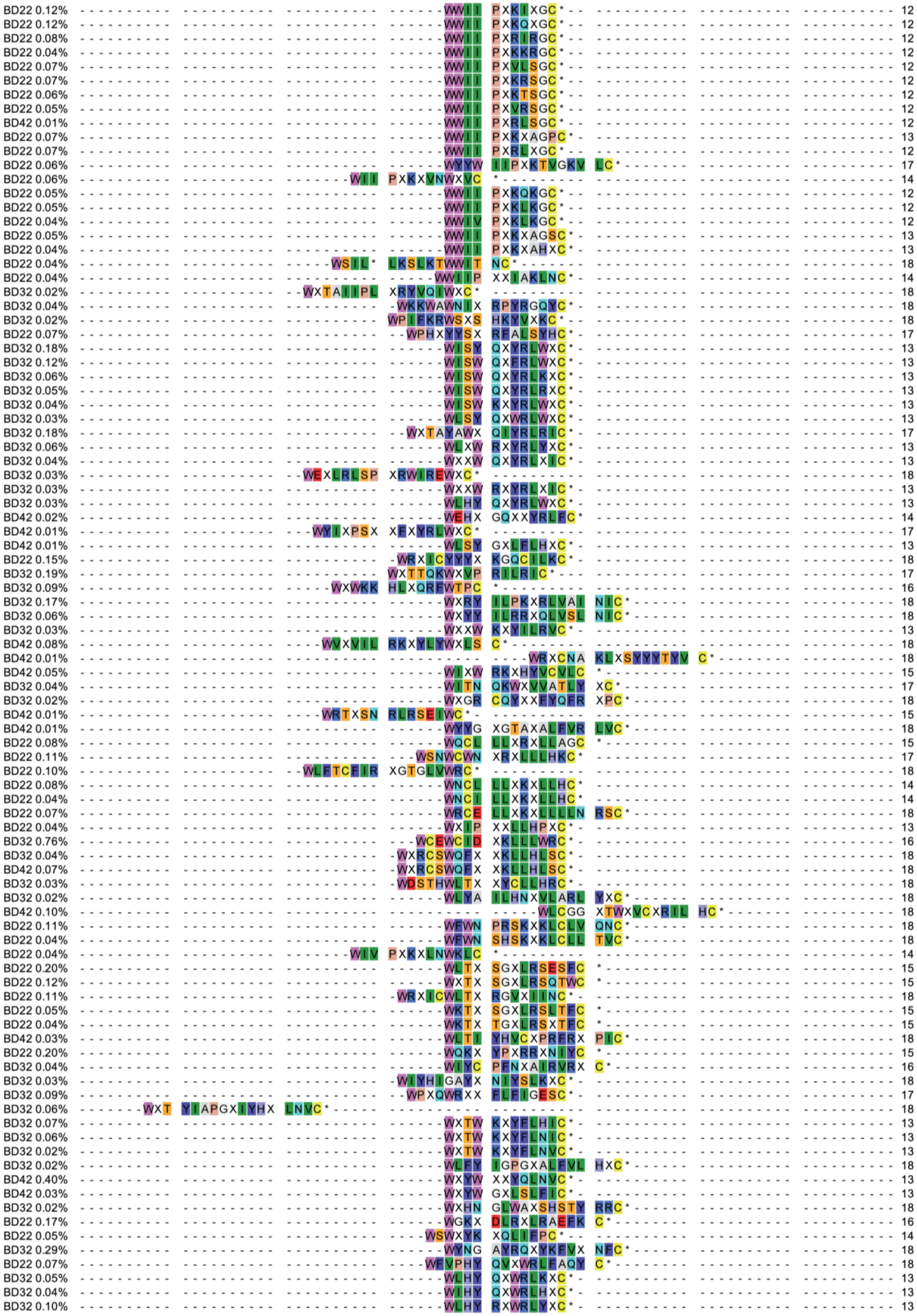

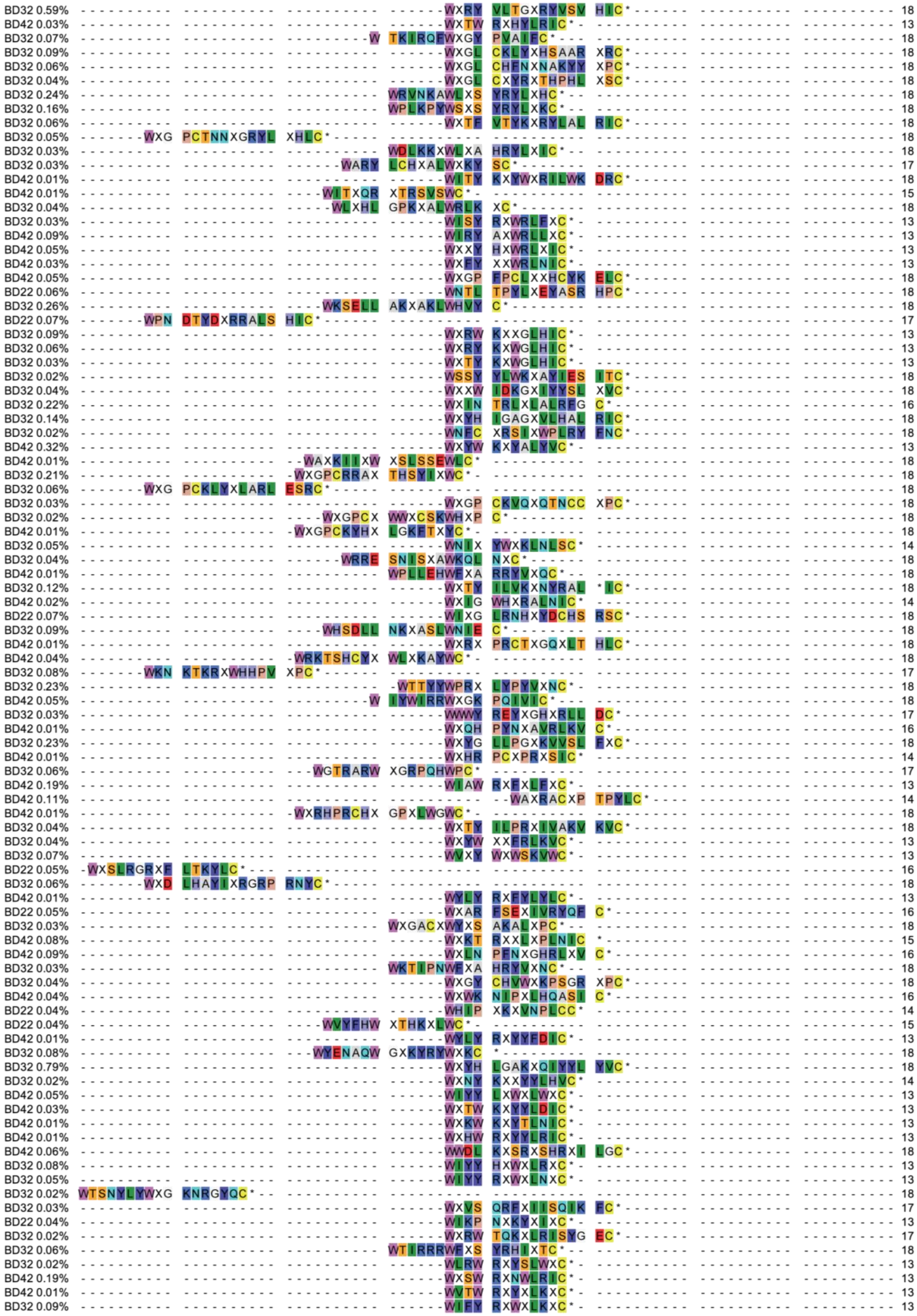

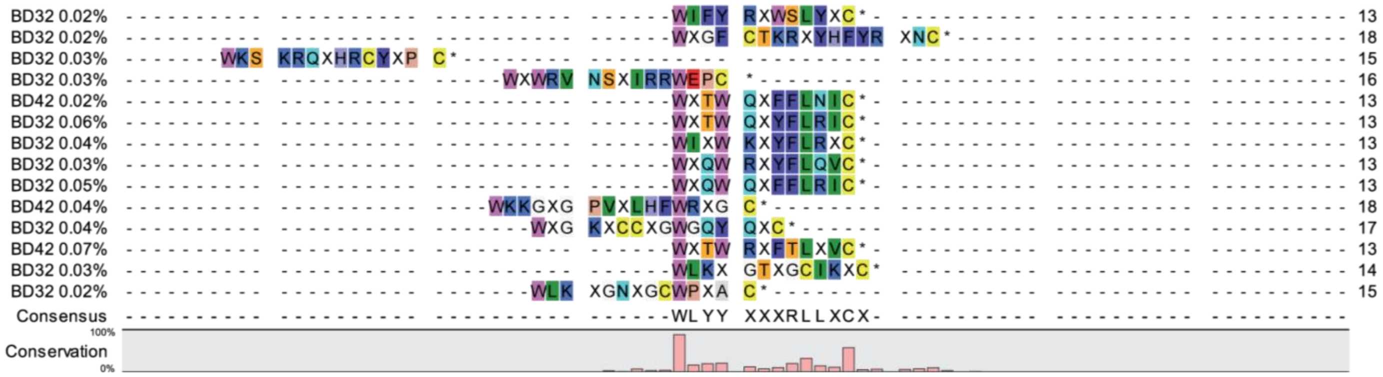
BD32 0.02%
BD32 0.02%
BD32 0.03%
BD32 0.03%
BD42 0.02%
BD32 0.06%
BD32 0.04%
BD32 0.03%
BD32 0.05%
BD42 0.04%
BD32 0.04%
BD42 0.07%
BD32 0.03%
BD32 0.02%
Consensus
WLYY XXXRLLXCX
Conservation
100%
0%
13
18
15
16
13
13
13
13
13
18
17
13
14
15

**Supplementary Table 5.** Data collection and refinement statistics for the crystal structure of BRD4-BD2 in complex with **4.2E** (PDB ID: 8CV4) solved using molecular replacement.

| | **BRD4-BD2:4.2E** |
|---|---|
| **Data collection** | |
| Space group | P 21 21 21 |
| Cell dimentions | |
| *a, b, c* (Å) | 53.58, 65.96, 76.39 |
| *α, β, γ* (°) | 90.00, 90.00, 90.00 |
| Resolution (Å) | 1.93 (1.93-1.97) |
| $R_{merge}$ | 0.079 (0.749) |
| $I/\sigma I$ | 19.6 (3.4) |
| CC(1/2) | 1.000 (0.907) |
| Completeness (%) | 99.6 (94.4) |
| Redundancy | 13.1 (6.6) |
| **Refinement** | |
| Resolution (Å) | 1.93 |
| No. reflections | 20881 |
| $R_{work}/R_{free}$ | 0.2090/0.2545 |
| No. atoms | 2389 |
| Protein | 1884 |
| Ligand/ion | 387 |
| Water | 118 |
| *B*-factors | 29.00 |
| R.m.s. deviations | |
| Bond lengths (Å) | 0.0071 |
| Bond angles (°) | 1.0026 |

*Values in the parantheses are for the highest resolution shell.
All data were collected on a single crystal.

**Supplementary Table 6.** Data collection and refinement statistics for the crystal structure of BRD4-BD2 in complex with **4.2D** (PDB ID: 8CV6) solved using molecular replacement.

| | BRD4-BD2:4.2D |
|---|---|
| **Data collection** | |
| Space group | P 1 21 1 |
| Cell dimentions | |
| *a, b, c* (Å) | 29.44, 59.64, 36.27 |
| *α, β, γ* (°) | 90.00, 100.01, 90.00 |
| Resolution (Å) | 1.7 (1.7-1.73) |
| $R_{merge}$ | 0.073 (0.566) |
| $I/\sigma I$ | 15.9 (3.0) |
| CC(1/2) | 0.999 (0.851) |
| Completeness (%) | 97.8 (93.0) |
| Redundancy | 7.1 (6.6) |
| **Refinement** | |
| Resolution (Å) | 1.7 |
| No. reflections | 29500 |
| $R_{work}/R_{free}$ | 0.2115/0.2614 |
| No. atoms | 1195 |
| Protein | 932 |
| Ligand/ion | 161 |
| Water | 102 |
| *B*-factors | 19.00 |
| R.m.s. deviations | |
| Bond lengths (Å) | 0.0070 |
| Bond angles (°) | 0.8270 |

*Values in the parantheses are for the highest resolution shell.
All data were collected on a single crystal.

**Supplementary Table 7.** Data collection and refinement statistics for the crystal structure of BRD3-BD2 in complex with **4.2D** (PDB ID: 8CV5) solved using molecular replacement.

| | BRD3-BD2:4.2D |
|---|---|
| **Data collection** | |
| Space group | P 32 2 1 |
| Cell dimentions | |
| *a, b, c* (Å) | 76.07, 76.07, 57.91 |
| *α, β, γ* (°) | 90.00, 90.00, 120.00 |
| Resolution (Å) | 1.47 (1.47-1.49) |
| $R_{merge}$ | 0.083 (0.906) |
| $I/\sigma I$ | 22.8 (3.5) |
| CC(1/2) | 1.000 (0.882) |
| Completeness (%) | 99.9 (99.0) |
| Redundancy | 20.3 (19.7) |
| **Refinement** | |
| Resolution (Å) | 1.47 |
| No. reflections | 33292 |
| $R_{work}/R_{free}$ | 0.1800/0.1967 |
| No. atoms | 1248 |
| Protein | 929 |
| Ligand/ion | 161 |
| Water | 158 |
| *B*-factors | 18.00 |
| R.m.s. deviations | |
| Bond lengths (Å) | 0.0059 |
| Bond angles (°) | 1.0453 |

*Values in the parantheses are for the highest resolution shell.
All data were collected on a single crystal.
